# Neck Geometry and Shape Compactness as Additional Haemodynamic Modulators in Abdominal Aortic Aneurysms

**DOI:** 10.64898/2026.02.27.708461

**Authors:** Vijay Nandurdikar, Aryan Tyagi, Tejas Canchi, Alejandro F. Frangi, Alistair Revell, Ajay B. Harish

**Author notes:** **Author for correspondence:** Ajay B. Harish.

## Abstract

We present an automated, constraint-aware framework to generate demographically stratified virtual populations of AAAs and quantifies, at cohort scale, how neck geometry, shape compactness, and maximum diameter jointly modulate AAA haemodynamics. Using 258 CTA-derived cases, we generated 182 validated AAA geometries and ran 364 simulations, extracting 10 geometric descriptors and correlated with six haemodynamic biomarkers. The automated framework blends statistically grounded sampling, anatomical plausibility and regional morphing to provide a scalable route for reproducible computational fluid dynamics (CFD) to uncover geometry-biomarker relations at cohort scale. Proximal neck diameter and sphericity were the dominant determinants of wall shear magnitude. Neck diameter increased mean wall shear stress (WSS) (*r* ≈ 0.49) while reducing peak WSS_0.95_ (*r* ≈ −0.31) and low-time-averaged wall shear stress (TAWSS) area (*r* ≈ −0.27), whereas sphericity, a largely under-explored AAA shape descriptor, was the strongest inverse correlate of mean WSS (*r* ≈ −0.57). Maximum diameter minimally affected peak shear (*r* ≈ +0.06) but was associated with reduced mean WSS (*r* ≈ −0.41). Convexity, another such descriptor, was the dominant correlate of oscillatory shear, increasing OSI (*r* ≈ +0.46) while modestly lowering mean WSS (*r* ≈ −0.28). The framework reveals neck calibre and shape compactness, not maximum diameter alone, as dominant modulators of AAA haemodynamics.

## 1. Introduction

Abdominal Aortic Aneurysm (AAA) is a localised and irreversible dilation of the abdominal aorta, typically defined as an expansion exceeding 1.5 times the maximum transverse diameter measured at the centre of the aneurysmal sac (18-20 mm) [1, 2]. It predominantly affects older adults, with a higher prevalence in males and individuals with a history of smoking or cardiovascular disease [3, 4]. Population based analyses shows that approximately 32% of patients die before reaching hospital, and when these pre-hospital deaths are included the overall case fatality remains 78–83% [5].

Elective surgical intervention, such as Endovascular Aneurysm Repair (EVAR) or open surgery, is generally recommended when the maximum aneurysm transverse diameter (*D*_max_) exceeds 55 mm in men or 50 mm in women [6]. However, multiple studies have shown that AAA’s can rupture at even smaller diameters, particularly in women, challenging the adequacy of size based criteria [7–9] as a risk metric. Data from the Vascular Quality Initiative (VQI) registry reveal that over 11% of ruptured AAA’s occurred below the surgical threshold [10], and mean rupture diameters as low as 42 mm have been reported. Conversely, some large AAAs do not rupture and may not require immediate repair, including in elderly or patients with other co-morbidities [11, 12]. Studies have also suggested that the current size thresholds for surgery may be lower than necessary, and that many large aneurysms, especially those near the cut-off, can be safely monitored without immediate intervention [5, 13, 14]. Elective repair itself involves risk, with 30-day mortality ranging from 1-2% for EVAR and 3-4% for open surgery. Randomised trials such as UKSAT and ADAM have also found no long-term survival benefit for early repair of small AAA’s [15, 16]. In summary, these findings suggest that diameter alone may not provide a guide for treatment decisions.

In the recent years, in addition to maximum diameter, several other geometric parameters including, but not limited to, tortuosity, curvature, neck angles and diameters, sac length, and aneurysm volume, have been studied. Computational fluid dynamics (CFD) has been used to study their association with haemodynamic biomarkers like wall shear stress (WSS), time-averaged WSS (TAWSS), and oscillatory shear index (OSI) [17–19]. These studies have demonstrated that shape-driven features can significantly modulate flow behaviour and wall stress distributions. However, most of them are based on relatively small patient datasets and focus primarily on conventional geometric descriptors. More complex shape measures such as average radius, sphericity, and convexity remain largely under-explored in the AAA literature [20–22]. Further, correlation analyses rarely incorporate population stratification (e.g., by gender or age), despite emerging evidence that anatomical and haemodynamic differences exist between subgroups [23, 24]. These gaps underscore the importance of broader, demographically aware investigations that combine conventional and advanced geometric features. Most CFD-based AAA studies rely on small cohorts, often limited to tens or low hundreds of patients thus restricting statistical power and prevents meaningful subgroup analyses [25]. Further, these datasets are predominantly male-dominated such as the ENGAGE registry, where only 11% of the 1,263 patients were female, hindering generalisability to women despite their higher rupture risk at smaller diameters [26]. The Kyriakou et al.[27] dataset employed in the present work, comprising 258 EVAR-planning CTA scans (14% female), reflects the same demographic skew. Ethical restrictions, segmentation effort, and privacy concerns further complicate the sharing of clinical data and limit access to large, balanced cohorts [28].

These limitations on the lack of data has led to studies that explored synthetic data generation methods like generative AI approaches such as generative adversarial networks (GAN), variational autoencoders (VAE), diffusion models, and graph-based networks [29]. These methods have demonstrated the ability to synthesize anatomically plausible geometries and capture complex shape variability. For instance, the recently released AAA-100 dataset was curated to support such efforts by providing watertight vascular models derived from pre-operative computed tomography angiography (CTA) scans [30]. The survey by Lin et. al. [31] summarises the growing interest in medical community for the automation of image to simulation-ready mesh reconstruction through generative and implicit methods. These developments represent significant progress toward fully data-driven in-silico approaches. However, many ML-based shape generators remain inherently black-box in nature and often lack anatomical interpretability or user-level control over specific geometric parameters. As noted in recent literature, even advanced architectures like VAE’s and diffusion models may struggle to enforce topological correctness or physiologically meaningful variation without explicit priors or segmentation supervision [31, 32]. Consequently, while these models are powerful, complementary approaches that allow controlled geometric perturbation and parametrisation are still needed for tasks involving hypothesis testing or systematic investigation of shape-function relationships and physiological constraints [33].

Interactive frameworks like svMorph [34] and Harvis [35] offer solutions with precise geometric manipulation, but the current versions are not fully automated and require manual intervention. In svMorph, the operator must pin and drag individual centreline control points and fine-tune an influence radius for each local change. Harvis requires a virtual reality headset and hand held controllers to sculpt the vessel surface in real time. In the current form of the two platforms, every edit is interactive and neither platform exposes a batch processing interface, producing the hundreds of variants needed for population phenotype-based trials is prohibitively labour intensive. This has motivated increasing interest in frameworks that can synthesize large, diverse, and physiologically acceptable AAA geometries in an automated, scalable, and constraint-aware fashion. Such frameworks can enable in-silico, systematic investigation of geometry–hemodynamic relationship, and machine learning models.

This work makes four major contributions, namely

i. Development of a constraint-aware, demographically stratified virtual cohort generator
ii. Cohort-scale CFD investigation linking conventional geometric descriptors (i.e. neck diameters, maximum diameter, distal diameter, tortuosity) and previously under-explored compactness descriptors (sphericity, convexity, fractal dimension) to CFD biomarkers under two inlet velocity profiles
iii. Demonstration that neck diameter and shape compactness dominate WSS and OSI patterns at population scale
iv. A reproducible, openly-released pipeline (OpenFOAM^®^ cases and a subset of geometries)

## 2. Materials and Methods

The following subsections describe each stage of the methodological pipeline including statistical distribution derivation, synthetic geometry generation and CFD workflow.

### 2(a) Dataset description

The dimensional records of 258 EVAR patients reported by Kyriakou et al. [27] (222 males, 34 females, 2 unspecified) are used in this work. All patient data corresponds to pre-operative CTA acquired for EVAR planning and the patients underwent EVAR. Diameters were measured on CTA slices orthogonal to the centreline and calculated to the intima (inner vessel wall), with stated accuracy of ±1 mm (±1.5 mm for maximum diameter). The measurements were performed by trained M2S clinical imaging technicians, and the study authors independently verified reproducibility on a subset of scans.

#### (2(a)(i)) Parameter selection

Among the parameters extracted from the dataset, four anatomical parameters were identified as critical determinants of AAA geometry based on their established clinical significance in EVAR planning. They consist of aneurysm diameter measurements at specific locations, as illustrated in Figure 1 and defined in Table 1. Aortic diameter is known to substantially influence haemodynamic behaviour in AAA [36–38], motivating its characterisation at these locations.

**Table 1.**
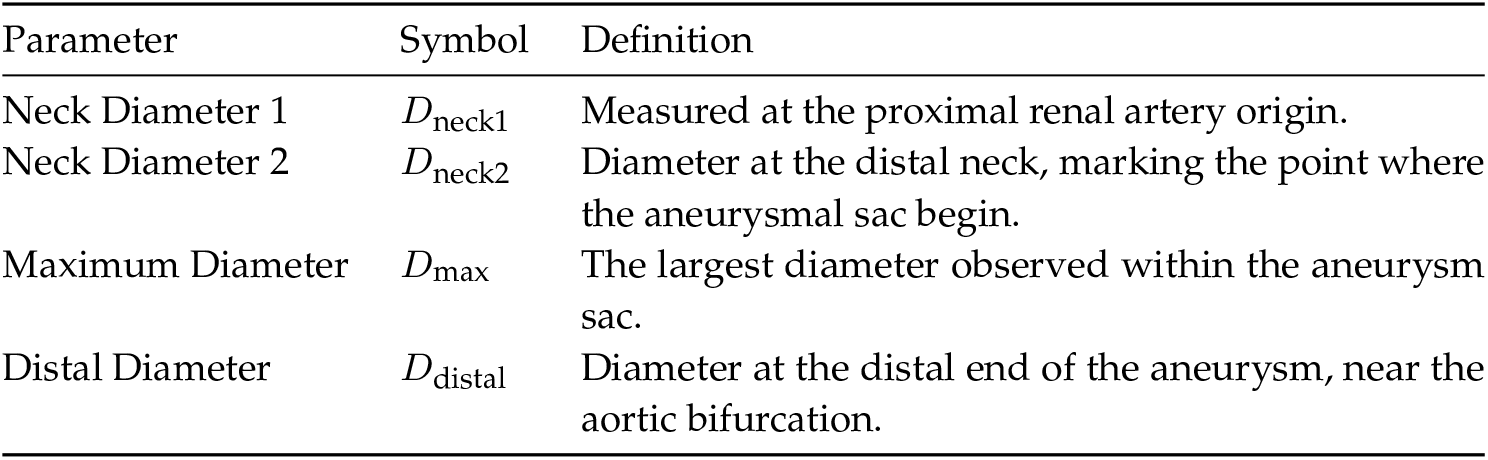
Key anatomical parameters used for AAA geometric characterisation.

| Parameter | Symbol | Definition |
| --- | --- | --- |
| Neck Diameter 1 | $D_{\text{neck1}}$ | Measured at the proximal renal artery origin. |
| Neck Diameter 2 | $D_{\text{neck2}}$ | Diameter at the distal neck, marking the point where the aneurysmal sac begin. |
| Maximum Diameter | $D_{\text{max}}$ | The largest diameter observed within the aneurysm sac. |
| Distal Diameter | $D_{\text{distal}}$ | Diameter at the distal end of the aneurysm, near the aortic bifurcation. |

**Figure 1.**
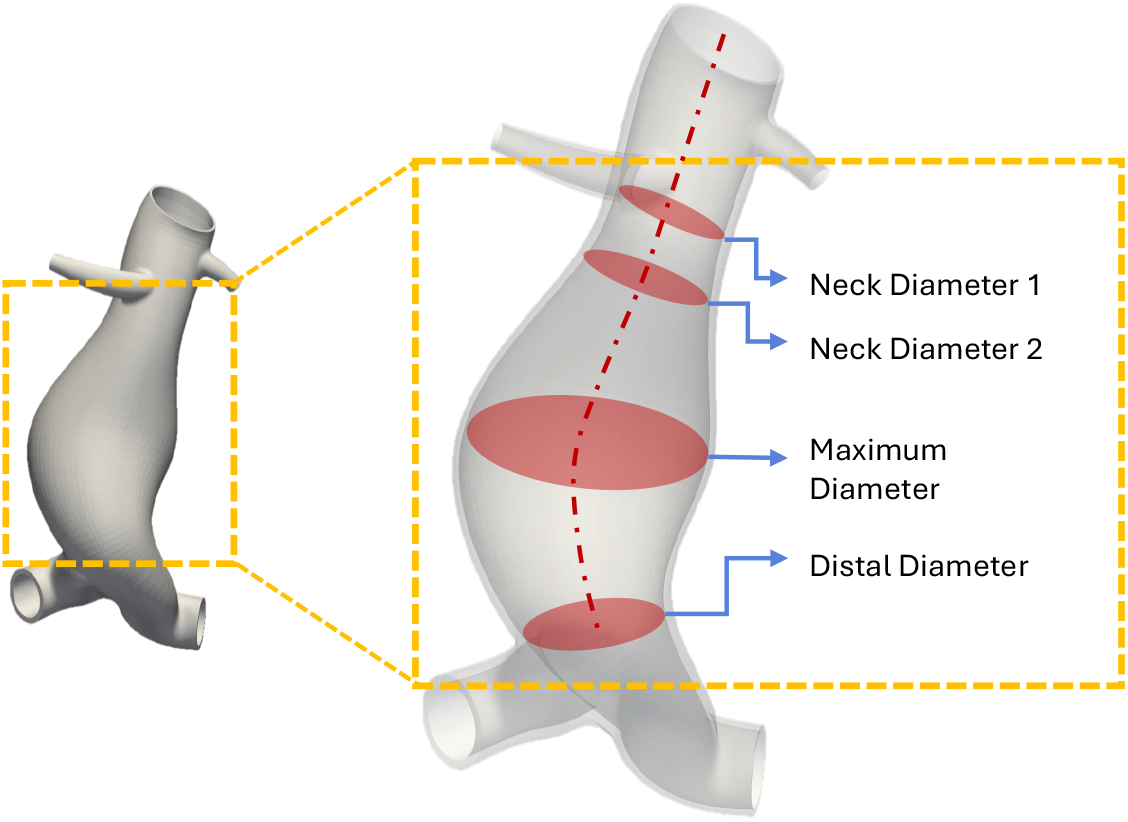
Anatomical landmarks used for geometric parameterisation of AAA: Neck diameter 1, neck diameter 2, maximum diameter, and distal diameter.

These locations span the longitudinal extent of the aorta from the proximal neck, adjacent to the renal arteries, through the aneurysm sac to the aortic bifurcation. This selection reflects EVAR planning convention where proximal neck diameters determine the fixation and sealing zone, the maximum diameter governs repair eligibility, and the distal diameter characterises the landing zone, providing a compact but clinically grounded description of the aneurysm along its axis. In the present work, additional geometric parameters were also derived from these landmarks, as discussed later in Section 2(b)(v).

#### (2(a)(ii)) Demographic stratification

The dataset spans patients aged 51–88 years. The proposed framework stratified the dataset by gender and four age brackets, namely 50–59, 60–69, 70–79, and 80+ years. The grouping highlights gender and age related differences in AAA geometry and supports modelling within each subgroup. The framework allows researchers to adjust the age-band boundaries to suit alternative clinical thresholds or study designs. This stratification therefore characterises geometry within an EVAR treated cohort and supports generation of clinically realistic intervention-stage morphologies. However, it does not claim to reproduce the full spectrum of AAA anatomy in the general screening/surveillance population.

#### (2(a)(iii)) Distribution fitting

Probability distributions were fitted to the selected anatomical parameters to quantify the observed variability in the AAA geometry. Normal, lognormal, gamma, and Weibull distributions were selected based on their common usage in biomedical modelling [39–42]. Right-skewed log-normal patterns are particularly well-precedented in AAA context where stochastic models of aneurysm growth [43] and AAA wall thickness measurements [44] both follow log-normal distributions. Maximum likelihood estimation (MLE) [45] was used for parameter estimation, and distributional fit was evaluated using the Kolmogorov–Smirnov (KS) test, p-values, and the sum of squared errors (SSE) [46]. For some subgroups (e.g. females aged 50–59 and 80+ years), the sample size was insufficient to support reliable distribution fitting. These strata were therefore excluded from parametric modelling. However, future work could address this limitation using hierarchical (partial-pooling) approaches. Accordingly, results should be interpreted with caution when generalising to under-represented subpopulations.

The best fitting distributions have been used to generate statistical variants as shown discussed in Section 2(b)(ii) while the results, criteria hierarchy and sensitivity on the distribution choices are discussed in Section 3(a).

### 2(b) Geometric modelling

#### (2(b)(i)) Base geometry generation

The base geometry generation was done in two parallel steps, including centreline generation and cross-sectional profile generation.

##### Centreline generation

The base centreline was generated using interpolating splines, defined through a set of anatomical coordinates extracted from patient specific geometries. The framework is flexible, allowing users to substitute these coordinates with points from other datasets or patient specific geometries. Additional control points can also be included to modulate the curvature or make use of the centreline points as needed for different anatomical shapes.

Interpolating splines were chosen for their ability to pass exactly through the specified points, in contrast to B-splines or Bézier curves, which approximate the shape based on control points [47]. The inlet plane (at *y* =0 mm, inlet) is placed 30 mm proximal to the first anatomical landmark *D*_neck1_ (at *y* = 30 mm, the distal renal artery level [27]), creating a 30 mm extension of constant circular cross-section equal to *D*_neck1_ (coordinates listed in Table 4, Appendix A). This extension facilitates the application of fully developed velocity profiles. The resulting centreline preserves anatomical alignment and supports consistent geometric reconstruction for downstream analysis.

##### Cross-sectional profiles

Cross-sectional diameters were initialised at anatomically defined points and additional control locations along the aorta. Parameters such as the inlet diameter and intermediate control point diameters were specified explicitly and remain unchanged during subsequent sampling process.

Two approaches for plane orientation were considered when generating the cross-sectional profiles: aligning the planes parallel to the primary axis, which mirrors the methodology used in generating CTA scans, and secondly orienting the planes perpendicular to the centreline, as per the data available from existing literature [27]. The latter option was adopted to ensure consistency with the patient specific data, necessitating perpendicular orientation of the cross-sectional profiles to the centreline.

#### (2(b)(ii)) Statistical variant generation

Statistical variants of AAA geometries were generated by sampling diameter values from the fitted probability distributions described in Section 2(a)(iii). These distributions were stratified by age and gender to reflect demographic specific anatomical variability.

##### Sampling procedure

Each anatomical diameter, *D*_neck1_, *D*_neck2_, *D*_max_, and *D*_distal_ were sampled using random variate sampling. In order to ensure anatomical realism, sampling employed a rejection-based approach,

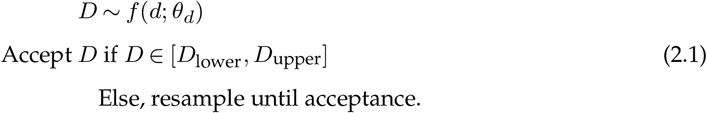

where, *f*(*d*; *θ*_*d*_) denotes the probability density function with demographic specific parameters *θ*_*d*_, [*D*_lower_, *D*_upper_] defines the clinically plausible range. If a valid sample was not obtained after a set number of iterations, the process defaulted to the distribution mean. All parameters were sampled simultaneously to form complete geometries.

##### Anatomical constraints

Physiological plausibility is enforced by applying a set of anatomical constraints immediately after diameter sampling. First, the sac is required to scale appropriately with respect to the necks, such that the maximum aneurysm diameter exceeds both the neck diameters (*D*_max_ > *D*_neck1_, *D*_max_ > *D*_neck2_). This ensures that the resulting geometry captures the characteristic dilation of aneurysmal segments. Second, all diameters must be positive (*D*_*i*_ > 0). The framework also supports optional user defined criteria to accommodate specific anatomical requirements; for example, a minimum distal diameter threshold of *D*_distal_ ≥ 7 mm can be imposed to reflect endograft delivery constraints [48]. Only geometries satisfying both the core conditions and any supplementary rules are retained for subsequent perturbation and morphing. However these constraints do not capture the joint relationships among all the diameters, which are enforced at a later stage by the population bounds analysis (Section 2(b)(v)).

#### (2(b)(iii)) Surface generation

The abdominal aortic surface was constructed by continuously triangulating successive cross-sectional profiles along the centreline. The resulting vertex-centred triangular mesh offers independent control of resolution in the longitudinal and circumferential directions through two parameters, namely the number of spline points along the centreline and the number of vertices per cross-section, respectively. This approach yields a geometrically smooth, *C*^1^-continuous surface; however, it lacks the finer undulations characteristic of in vivo vessel walls. Cross-sections are oriented perpendicular to the local centreline tangent using a parallel-transport frame, which propagates an orthonormal basis along the centreline without accumulating twist, ensuring consistent cross-section orientation along curved paths. This construction produces a non-self-intersecting surface provided that *R*(*s*) > *r*(*s*) at every arc-length station *s*, where *R*(*s*) is the local centreline radius of curvature and *r*(*s*) is the local vessel radius. A technical description and capability illustration are given in Appendix C. This constraint limits the maximum centreline angulation that can be synthesised.

Physiological wall roughness was approximated by introducing stochastic perturbations to the nodal coordinates using Latin Hypercube Sampling (LHS). For a vertex **v**(*x, y, z*), the perturbed position **v′** is defined as:

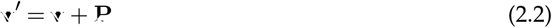

where **P** is a random vector with components *P*_*i*_ ∼ LHS([−*r*, +*r*]), ∀*i* ∈ {*x, y, z*}. The perturbation range *r* could be chosen based on wall roughness magnitudes reported in clinical imaging studies. Stratified sampling via LHS ensured uniform coverage of the perturbation space and mitigated artificial clustering common with Monte Carlo approaches [49]. Perturbations were restricted for the nodes associated with the inlet and outlet to preserve flat surface patches required for stable CFD simulations.

Although the perturbation scheme improves realism by introducing localised surface irregularities, it cannot reproduce larger scale asymmetries or regional deformations characteristic of pathological AAA shapes. This limitation motivates the complementary, and additionally employed, morphing strategy described in the following section, which introduces broader geometric variation while retaining anatomical plausibility.

#### (2(b)(iv)) Shape morphing

The morphing algorithm introduces larger-scale asymmetries through a spherical control point-based deformation approach. In order to prevent distortions at physiologically critical regions, the aorta is segmented along the centreline into three distinct zones: (i) fixed regions at the inlet and outlet, where no deformation is applied preserving the sampled cross-sectional geometry and maintaining boundary condition integrity; (ii) buffer zones, which serve as smooth transition regions; and (iii) a central morphable region that undergoes full deformation to introduce geometric variability (Figure 2).

**Figure 2.**
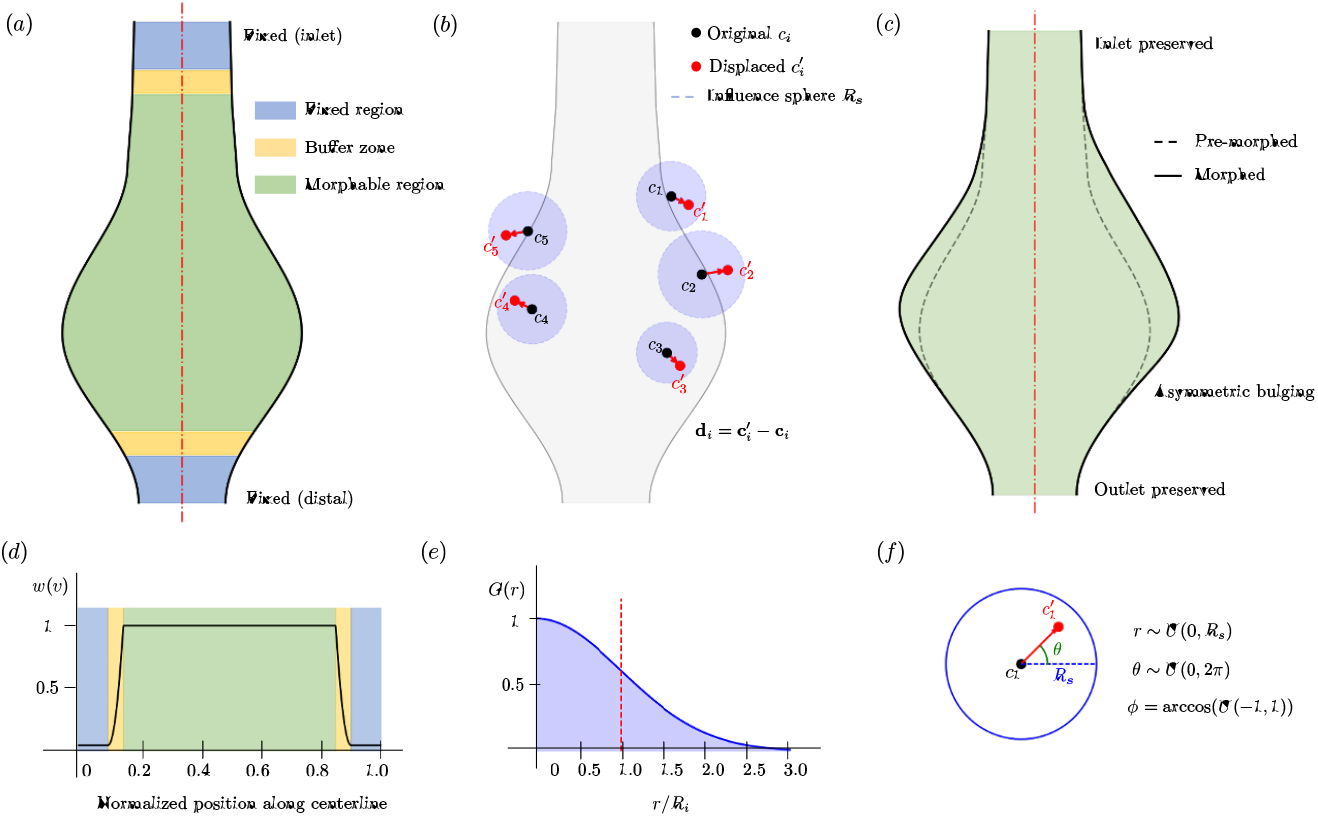
Shape morphing workflow: (a) original geometry with morphing zones, (b) control point deformation, (c) morphed geometry, (d) weighting function, (e) Gaussian kernel, and (f) spherical sampling (2D projection).

##### Regional weighting and transition control

The deformation process employs spatial weighting to ensure smooth transitions between regions. For each vertex **v** in the mesh, a transition weight *w*(**v**) ∈ [0, 1] is assigned based on its position relative to the defined regions

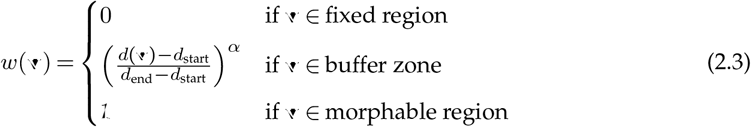

where *d*(**v**) denotes the distance of vertex **v** along the centreline, *d*_start_ and *d*_end_ define the boundaries of the buffer zone, and *α* =2 determines the smoothness of the polynomial transition function.

##### Control point-based deformation

The morphing process introduces local perturbations using *n*−randomly sampled control points within the morphable region. Each control point **c**_*i*_ is allowed to move within a sphere of radius *R*_*s*_, resulting in a new position 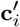 and a displacement vector defined using spherical coordinates

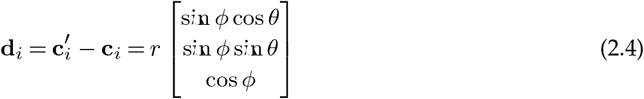

where *r* ∼ *U*(0, *R*_*s*_) is the radial displacement, *ϕ* = arccos(*U*(−1, 1)) is the polar angle, and *θ* ∼ *U*(0, 2*π*) is the azimuthal angle. This ensures uniform sampling within the spherical region.

##### Mesh vertex displacement

Each mesh vertex **v** is displaced by aggregating the influence of all control points, weighted by both spatial proximity and the regional transition weight

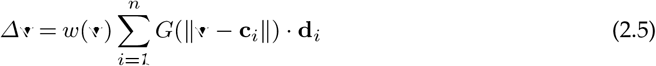

The influence of each control point on surrounding vertices is modelled using a Gaussian kernel [50]

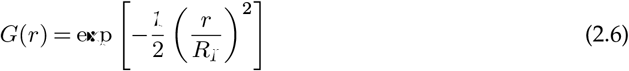

where *R*_*I*_ is the influence radius that determines the spatial extent of deformation around each control point. The updated vertex position is

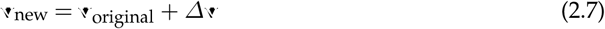

##### Mesh smoothing

Lastly, a Laplacian smoothing step [51] is applied following the deformation to improve mesh quality without distorting introduced geometric variations. This smoothing process operates independently of the Gaussian kernel and is based on local vertex connectivity

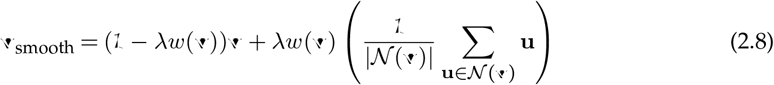

where *N*(**v**) denotes the set of neighbouring vertices of **v** in the mesh, and *λ* is the smoothing factor. The regional weight *w*(**v**) ensures that smoothing is applied only in the deformable and buffer regions, leaving fixed regions unchanged.

All morphed variants share a common base centreline path; the variability introduced by morphing therefore manifests as surface-level asymmetries that translate into derived geometric descriptors (tortuosity, sphericity, convexity) used in the correlation analysis. Inter-patient variation in the axial centreline path is not represented as an independent sampling dimension.

Since the morphing step modifies the geometry, each morphed variant is re-checked against the anatomical constraints defined earlier in Section 2(b)(ii). Additionally, all morphed geometries are validated against statistical distributions using Kullback-Leibler (KL) divergence [52] to ensure statistical similarity with the original patient data distributions. This quantitative validation ensures that generated variations remain within physiologically realistic limits, with lower KL divergence values (< 0.3) indicating better preservation of the original statistical properties.

#### (2(b)(v)) Population bounds analysis

Individual diameter measurements may lie within their respective statistical ranges yet still combine into anatomically unrealistic shapes. For example, pairing a very small neck diameter 2 with a large aneurysmal maximum creates an abrupt transition not generally seen in clinical data. A population bounds analysis was applied to check multivariate consistency to enable screening out cases and accept only geometries whose joint parameters match observed anatomy. The procedure had three steps: convex hull construction, outlier detection, and interior-point validation.

##### Convex Hull construction

Convex hulls were built for three clinically important diameter pairs: (i) neck diameters (*r* = 0.901, *p*< 0.01), (ii) neck diameter 2 - maximum diameter (*r* = 0.111, *p*< 0.05), and (iii) maximum diameter - distal diameter (*r* = 0.208, *p*< 0.01). Each hull represents the convex region containing all patient points and therefore sets a data-driven outer limit for a normal anatomy. In this work, pairwise (i.e. 2D) hulls were considered as opposed to multivariate hull in higher dimensions. This enables interpretability and sample-size stability to enable informed decisions.

A given set of patient data points, represented as *P* = {*p*_1_, *p*_2_,. .., *p*_*n*_} in 2D parameter space, where each point *p*_*i*_ = (*x*_*i*_, *y*_*i*_) represents a diameter pair, the convex hull ℋ is defined as

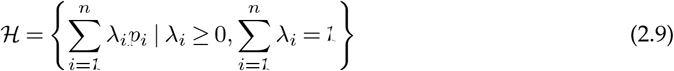

where *λ*_*i*_ are the convex combination coefficients. The boundary vertices of the convex hull are extracted using SciPy® ConvexHull implementation, which employs the Quickhull algorithm [53] for efficient computation.

##### Outlier detection

The presented framework can apply five complementary techniques to flag parameter outliers: 1) modified Z-score analysis identified outliers based on the median absolute deviation, providing robustness against extreme values [54]; 2) the Interquartile Range (IQR) method established boundaries using quartile-based thresholds; 3) Mahalanobis distance calculations accounted for the covariance structure between parameters; 4) DBSCAN clustering identified outliers through density-based spatial clustering; and 5) a manual method incorporated expert-defined exclusion criteria based on clinical knowledge. The framework allows user to choose any method on desired level of stringency.

##### Interior-point validation

Each candidate geometry was subjected to an interior-point test against the convex hull ensemble described in Section 2(b)(v). **d** = {*D*_neck1_, *D*_neck2_, *D*_max_, *D*_distal_} denotes the diameter vector for a generated case, *H*_*ij*_ represents the convex hull formed by the *i*th–*j*th diameter pair in the clinical dataset. A geometry satisfies the anatomical consistency criterion when

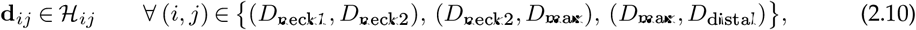

where **d**_*ij*_ is the two-dimensional projection of **d** onto the corresponding parameter pair.

Interior point testing is performed using a winding number algorithm [55] that determines point containment by computing how many times the convex hull boundary winds around the test point. For each hull ℋ _*ij*_, a point is inside if the winding number is non-zero. This approach provides reliable containment testing for the 2D parameter projections. This method is chosen over others, like ray casting, considering its effectiveness in handling self-intersecting polygons.

A geometry is acceptable only if it lies inside all three pairwise hulls, thereby forming the universal interior set. This multivariate filter removed combinations that, while marginally plausible in one projection, produced unrealistic transitions when considered jointly. It is worth noting that although hull-exterior cases might still represent rare but valid anatomies, restricting the CFD analysis to the universal interior set ensured that subsequent hemodynamic trends were not driven by extreme geometries. The retained cohort therefore offers the reliable basis for correlating shape descriptors with wall-shear and oscillatory metrics.

The comparison of morphed geometries that were satisfied atleast one convex hull vs. universal interior criterion is outlined in detail in Appendix Section J(c) and the viability of the morphing validation is given in Section 3(e).

##### Geometric parameterization

In addition to the primary anatomical diameters discussed earlier in Section 2(a)(i), a set of derived metrics are computed from the re-constructed surface mesh and centreline to capture the three-dimensional complexity of AAA morphology. While traditional AAA assessment focuses on conventional geometric descriptors, this study incorporated under-explored parameters that have shown significant predictive value in cerebral aneurysm research. These derived parameters provide a more holistic representation of shape features that may influence intra-luminal flow dynamics.

The total aneurysm volume (*V*) was calculated by performing tetrahedralisation of the surface mesh, capturing the full enclosed lumen. The corresponding surface area (*A*) is used to quantify the endothelial surface in contact with blood flow. The centreline length (*L*_*c*_), defined as the arc-length of the centreline extracted from the morphed surface mesh, is used as a measure of the vessel’s axial elongation, the tortuosity (*τ*) is used to quantify the deviation from a straight configuration using the ratio *L*_*c*_/*L*_s_, where *L*_s_ denotes the Euclidean distance between inlet and outlet points.

Shape compactness was quantified through the sphericity index,

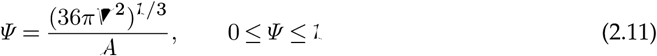

where *V* and *A* denote the aneurysm lumen volume and surface area, respectively [56]. A perfect sphere yields *Ψ* = 1. Recent comprehensive studies on cerebral aneurysms consistently rank sphericity measures among the most important morphological predictors [57]. Surface irregularity was characterised by the convexity ratio

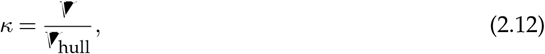

with *V*_hull_ denoting the volume of the convex hull that encloses the sac. Values close to unity indicate a nearly convex surface, whereas lower values reflect pronounced lobulations. Convexity-based metrics, including the undulation index, are moderately validated for cerebral aneurysms and show significant associations with rupture [58]. Currently, there is a limited discussion on the correlation between convexity and aneurysm growth [21] for AAA. Overall lumen size was summarised by the average radius

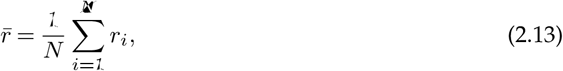

where *r*_*i*_ is the radial distance from the centreline to the *i*-th surface node (*N* vertices in total) along the orthogonal direction.

The present study adapts the established formulations of *Ψ, κ*, and 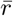 from past studies of cerebral-aneurysm modelling and leverages for synthetic AAA geometry development. Together, all the above parameters enable a comprehensive geometric description of the aneurysm. The pseudo-code of the formulations are provided in Section F. The transition from geometric models to CFD simulations requires the automated generation of case directories with appropriate boundary conditions and mesh specifications. More details on the case generation for OpenFOAM^®^ are provided in Appendix B.

### 2(c) Computational fluid dynamics framework

The blood flow in the abdominal aorta is modelled as an incompressible Newtonian flow. The choice of rheological model remains a point of discussion in aortic CFD. Shear-thinning effects grow with blood residence time and can become non-negligible in slow recirculation zones [59]. Newtonian models nonetheless remain widely used in AAA simulations for their simplicity and computational effieciency [25, 60, 61] and are physically justified in the aorta where shear rates typically exceed 100 s^*−*1^ [62]. Based on prior AAA CFD studies [63–67], the inflow is treated as pulsatile and (predominantly) laminar. This assumption is supported by the characteristic dimensionless numbers in the infrarenal aorta: taking a representative diameter *D*≈2 cm and mean velocity *U* ≈0.10 m s^*−*1^ yields Re = *ρUD*/*µ* ≈ 600 for *ρ*≈1060 kg m^*−*3^ and *µ*≈3.5 mPa s, with peak-systolic values typically a few times higher [68]. In vivo AAA inlet measurements compiled from Doppler ultrasound (the Fraser waveform) further indicate relatively low mean Reynolds numbers at the infrarenal level, with larger but transient systolic peaks [69]. The pulsatility is characterised by the Womersley number 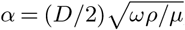, which is *O*(5–12) in the infrarenal/abdominal aorta for physiological heart rates [69, 70]. Given these ranges, a laminar Newtonian model provides an appropriate baseline for capturing the dominant haemodynamic trends in intervention-stage AAA geometries, while recognising that local transitional features may arise in highly disturbed sacs.

In this work, two major haemodynamic metrics are extracted from the CFD simulations, namely Wall shear stress (WSS) and flow oscillatory metrics. WSS is a key hemodynamic quantity representing the tangential force exerted by blood flow on the vessel wall. Variations in WSS have been believed as as potential mechanical biomarker for aneurysm progression, thrombus formation, and wall degradation. In order to characterise these shear forces across the aneurysm surface, specific WSS-related quantities are extracted from each CFD simulation, including mean cycle WSS (WSS_mean_), 95th percentile of WSS magnitude at the peak systolic phase (WSS_95%_) and TAWSS.

Oscillatory flow behaviour within AAA’s is characterized by bidirectional wall shear forces that have been implicated in endothelial dysfunction and pro-inflammatory responses. OSI provides a dimensionless metric to quantify the directional variability of wall shear stress over the cardiac cycle and ranges from 0 (purely unidirectional flow) to 0.5 (completely oscillatory flow).

The governing continuity and Navier–Stokes equations, and all notation used throughout, the physical relevance of these flow metrics and their formal integral definitions are described in the supplementary materials.

#### (2(c)(i)) Boundary conditions

A pulsatile velocity boundary condition was prescribed at the inlet using a population-averaged infrarenal abdominal aortic waveform adapted from Fraser et. al. [71]. Two spatial inlet velocity distributions were defined for each geometry: a uniform plug profile and a fully developed parabolic profile,

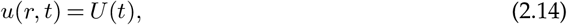

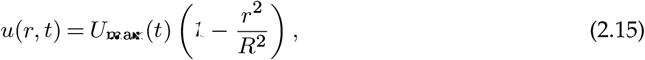

where *r* is the radial coordinate and *R* is the inlet radius. For the parabolic profile, the cross-sectional mean velocity satisfies

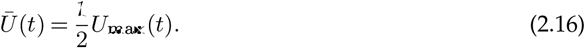

The implementation details of the inlet velocity specification are given in Appendix B. The zero gauge pressure was prescribed at the outlet [72, 73]. The domain terminates above the iliac bifurcation, giving a single outlet. Thus, there is no competing flow-split problem for which a three-element Windkessel model would be required. Since WSS, TAWSS, and OSI are near-wall velocity-gradient quantities governed primarily by the inlet waveform and vessel geometry rather than by the absolute outlet pressure datum, these metrics are less sensitive to this outlet condition. The aortic wall was modelled as rigid with a no-slip boundary condition because fluid–structure interaction studies show that wall compliance only causes minor changes to global hemodynamic metrics-peak wall stresses differ by less than 1% [74] and the overall flow field and wall shear stress patterns are similar to those in rigid-wall simulations [75]. All haemodynamic metrics were evaluated using data from the fourth cardiac cycle, with the preceding three cycles excluded to mitigate initial start-up transients [63, 68, 76]. Cycle-to-cycle convergence is demonstrated quantitatively in Appendix D. All 182 simulations used a fixed time-step of Δ*t* =1 × 10^*−*4^ s (10 000 steps per cardiac cycle). The Courant-number diagnostics extracted from the simulation logs confirm a time-averaged domain-mean Co = 0.022 (range 0.012-0.035 across cases) and a time-averaged domain-maximum Co = 0.36 (range 0.20-0.66). Thus, the bulk of the flow domain therefore operates well within the Co < 1 regime throughout all simulations. The absolute instantaneous peak Co reaches 2.4 in the most geometrically complex case, which is accommodated by pisoFoam’s implicit PISO algorithm [77]. This time-step is also consistent with or finer than temporal resolutions reported in comparable pulsatile AAA haemodynamic studies [78–80].

#### (2(c)(ii)) Geometry–flow correlation analysis

Linear associations between geometric features and hemodynamic responses were quantified using Pearson product–moment correlation coefficients [81]. The Pearson correlation coefficient (*r*) captures both the strength and direction of a relationship, taking values between −1 (perfect inverse correlation) and +1 (perfect direct correlation) and is given as

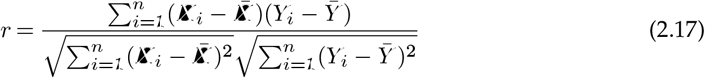

where *n*− observations are made for variables *X* and *Y* and the respective sample means are 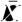 and 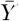.

Before performing the correlation analysis, data were preprocessed to exclude any parameter pairs with fewer than three valid (non-NaN) observations to ensure reliable estimates. Pairs involving missing data were removed listwise. This approach ensured interpretable correlation measures across the geometric and hemodynamic variables The final dataset comprised statistically and anatomically validated AAA geometries, each annotated with 10 geometric descriptors and six hemodynamic metrics derived from CFD simulations.

## 3. Results

A total of 182 synthetic AAA geometries were selected for CFD analysis from the universal-interior set spanning the age groups 60–69, 70–79, and 80–89 for both male and female cohorts. No geometries were generated for the 50–59 age group due to the absence of statistically and anatomically validated samples within this range. Details of the framework configuration can be found in Appendix A. In this work, we have used OpenFOAM v9^®^ for the CFD analysis. The average mesh size used in the work is approximately 1.3M with the average edge lengths in the range of 0.019 to 1.15 mm. A detailed mesh convergence is presented in Appendix E. This work uses a pisoFOAM solver, a transient solver based on the PISO (Pressure Implicit with Splitting of Operators) algorithm [77].

### 3(a) Distribution fitting

The age-wise fitted distributions from Section 2(a)(iii) are summarised in Table 2. For the combined dataset, the log-normal distribution provided the best fit, with the lowest SSE and a high KS p-value (*p* = 0.87), indicating good agreement with the data. In some male age groups, other distributions such as gamma (e.g. males aged 70-79) or Weibull gave slightly better fits. For female subgroups with very few patients (10 or fewer), reliable distribution fitting was not possible, so only summary values are reported. The detailed distribution fitting results for all anatomical diameters are included in Appendix G. In this study, the distributions are mainly used to generate synthetic geometries, as described in Section 2(b)(ii).

**Table 2.**
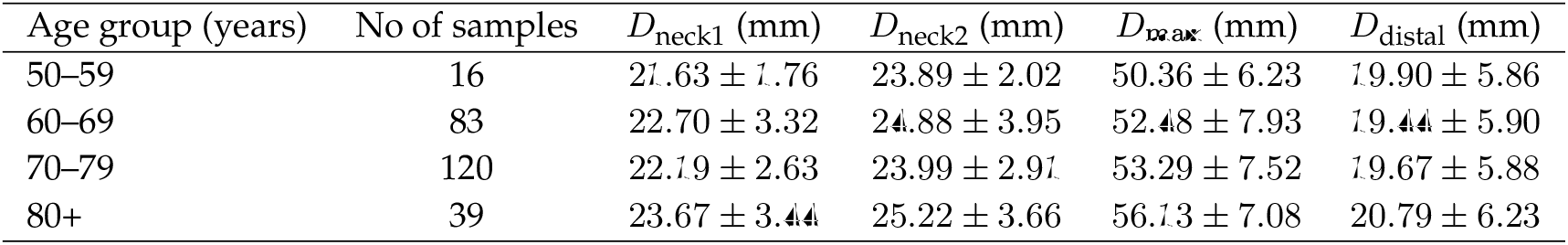
Age wise summary of anatomical diameters for all population. Values are reported as mean ± standard deviation.

**Table 3.** Top five strongest geometry–haemodynamics correlations under plug and parabolic inlet velocity profiles. Correlations are ranked by absolute Pearson correlation coefficient |*r*|.

| Input parameter | Output parameter | $r$ | $p$ |
| --- | --- | --- | --- |
| *Plug inlet profile* |  |  |  |
| Neck diameter 1 | WSS<sub>mean</sub> | 0.838 | < 0.001 |
| Convexity | OSI<sub>mean</sub> | 0.628 | < 0.001 |
| Convexity | Area with OSI > 0.3 (%) | 0.618 | < 0.001 |
| Convexity | WSS<sub>0.95</sub> (systolic) | −0.538 | < 0.001 |
| Convexity | TAWSS<sub>mean</sub> | −0.504 | < 0.001 |
| *Parabolic inlet profile* |  |  |  |
| Sphericity | WSS<sub>mean</sub> | −0.570 | < 0.001 |
| Tortuosity | Area with TAWSS < 0.4 (%) | −0.509 | < 0.001 |
| Neck diameter 1 | WSS<sub>mean</sub> | 0.492 | < 0.001 |
| Convexity | OSI<sub>mean</sub> | 0.455 | < 0.001 |
| Neck diameter 2 | WSS<sub>0.95</sub> (systolic) | −0.432 | < 0.001 |

### 3(b) Demographic variations

Anatomical differences across age and gender groups were clearly observed, highlighting the importance of demographic stratification when modelling AAA geometry. Among male patients, the maximum aneurysm diameter increased with age, from an average of 50.4 mm in the 50–59 age group to 56.3 mm in those aged 80 and above. This suggests that while most patients have moderate aneurysms, a small number reach much larger sizes at older ages, possibly due to long-term degeneration or delayed treatment. Neck diameter 1 showed less change in the mean across age groups but exhibited wider spread with age. The fitted distribution also shifted from Weibull in younger males to gamma or log-normal in older groups. This increasing variability could impact stent graft fixation, which depends on neck geometry. Other diameters, such as neck diameter 2 and distal diameter, showed smaller changes in average values but also shifted toward broader, right-skewed distributions in older patients.

Female patients consistently had smaller vessel diameters across all the considered parameters. The average neck diameter 1 in females was 12.45 mm (log-normal, *p* = 0.954), compared to 16.67 mm in males (Weibull, *p* = 0.813). Although males exhibited greater variability in distal diameter, the limited number of female cases (*n* = 34) makes it harder to generalise the findings. This under-representation is clinically important, as current stent graft designs are often based on male-dominated datasets and may not accommodate female patients and thus increasing the risk of sizing errors during EVAR. This study also highlights the need for more balanced and diverse datasets to further compare these observations.

### 3(c) Morphing effectiveness

The effectiveness of the morphing process in preserving anatomical realism was evaluated by comparing the distributions of morphed variants with patient data across age and gender groups. Figure 3 illustrates one such comparison for the maximum aneurysm diameter in males aged 60-69. The morphed variants closely follow the clinical data, with a KL divergence of 0.0180 indicating strong similarity.

**Figure 3.**
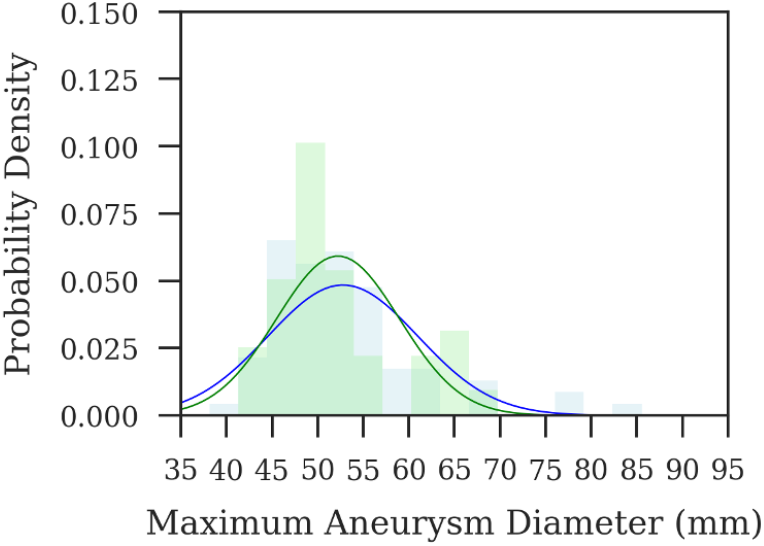
Probability density functions of maximum aneurysm diameter for patient data (blue) and morphed variants (green), males aged 60-69.

More detailed comparisons and divergence values are provided in Appendix J. KL divergence values below 0.3 for maximum diameter suggest that the morphing process retains key population-level characteristics. The higher divergence observed for neck diameter 1 likely results from limited deformation in the inlet region, due to the buffer zone defined by the weighting function in Equation (2.3). These constraints reduce morphological variability near the proximal end, explaining the discrepancy from clinical data.

### 3(d) Parameter detection and outlier analysis

Prior to analysis on the flow-morphology relation, it is pertinent to identify outliers that can distort the overall trends. Seven such cases were manually selected based on visual inspection. These cases represent atypical parameter combinations and are marked as squares in Figure 4. Manual selection in this context is intended for illustrative purposes, helping to reveal how extreme cases influence the observed trends.

**Figure 4.**
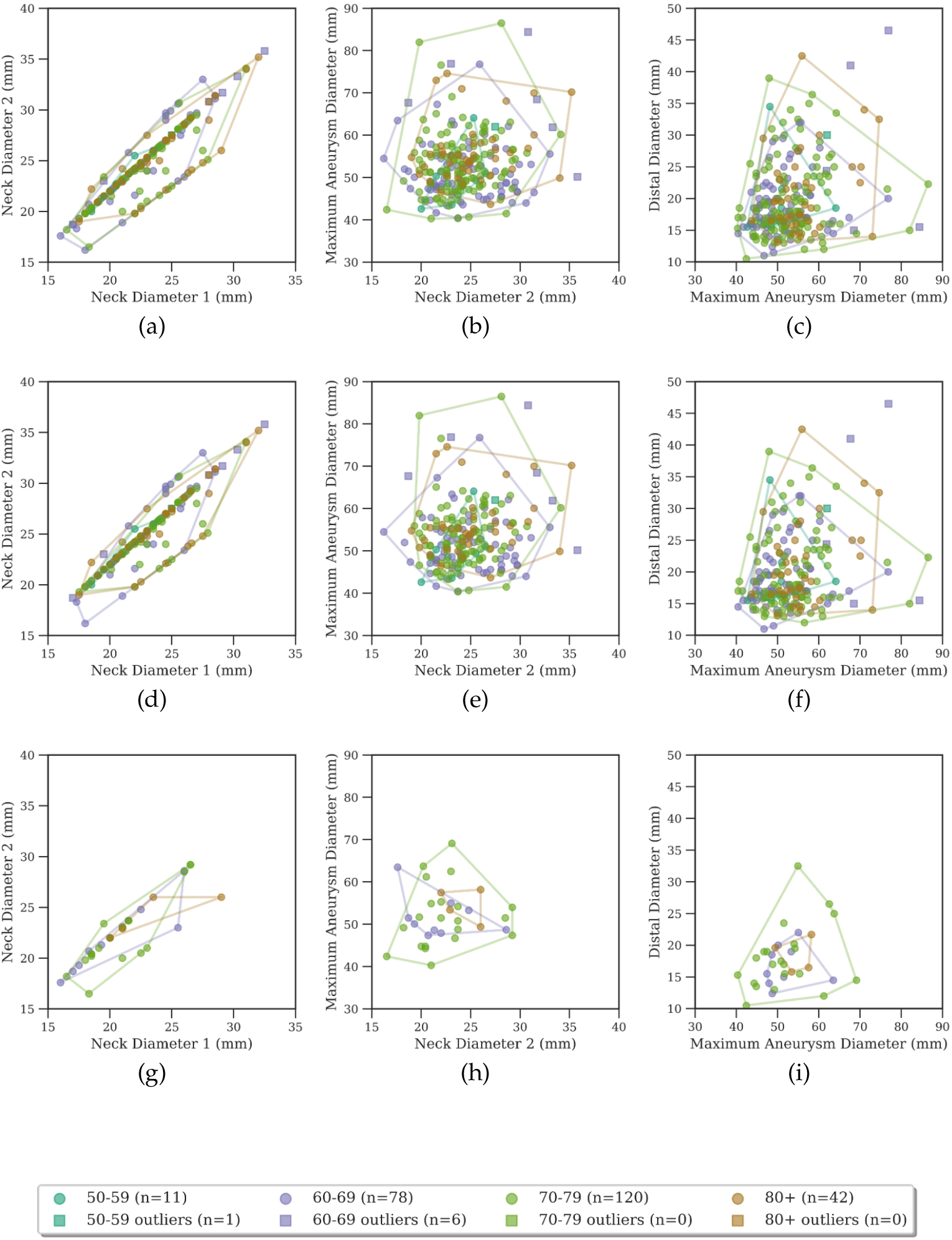
Parameter relationships with manually identified outliers. Top row: All population, Middle row: Male patients, Bottom row: Female patients. (a,d,g) Neck diameter relationships, (b,e,h) Neck diameter vs maximum aneurysm diameter correlations, (c,f,i) Maximum aneurysm diameter vs distal diameter correlations

Figure 4 presents pairwise parameter relationships across the full dataset and stratified by gender and clearer associations emerge, following the exclusion of outliers. The neck diameters show a strong positive correlation (Figure 4a, *r* = 0.901, *p*< 0.01), while the relationship between neck diameter 2 and maximum aneurysm diameters is weak (Figure 4b, *r* = 0.111, *p*> 0.05), and indicates potentially independent sac expansion.

Figure 5 isolates only the convex hulls for each age group, omitting the data points to help visualize the shift in the parameter space with age. A trend toward larger values particularly in maximum aneurysm diameter and distal diameter is observed in older cohorts. This progression likely reflects cumulative remodeling effects, such as wall weakening or dilation. Still, it is important to note that convex hulls are sensitive to sample distribution, and these visual patterns should be interpreted in that context. The framework supports statistical testing for wider generalisation when larger datasets are available.

**Figure 5.**
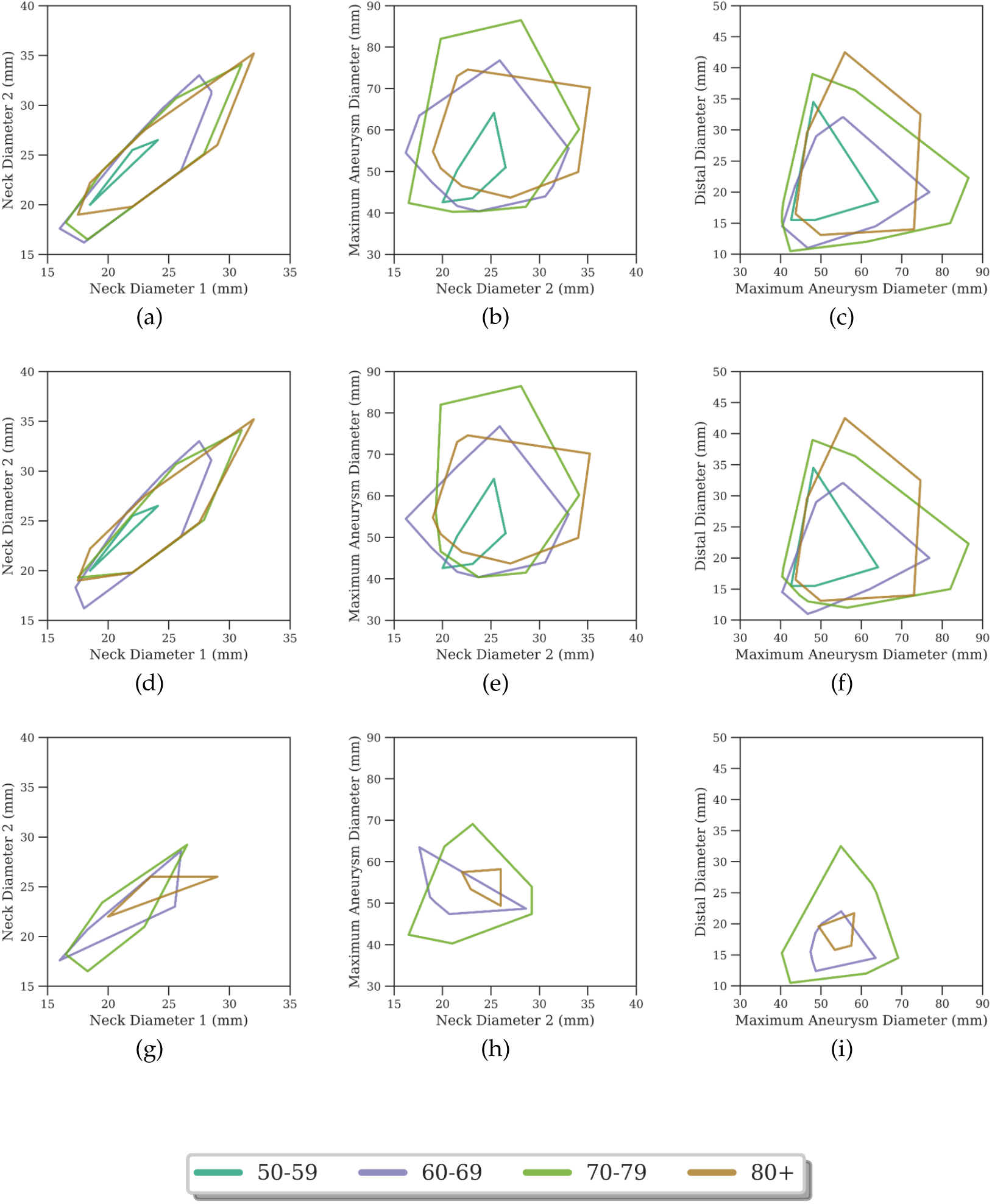
Convex hull analysis for AAA parameter relationships. Top row: All population, Middle row: Male patients, Bottom row: Female patients. (a,d,g) Neck diameter relationships, (b,e,h) Neck diameter vs maximum aneurysm diameter correlations, (c,f,i) Maximum aneurysm diameter vs distal diameter correlations.

### 3(e) Universal interior set analysis

The morphed samples are compared against patient data for female patients aged 70-79 in Figure 6. While some individual geometries extended beyond single-parameter boundaries, most of the generated cases stayed within observed population bounds. A stricter criterion was applied to identify the universal interior set-geometries that remained within the convex hull across all parameter pairs simultaneously. These cases are shown in Figure 7.

**Figure 6.**
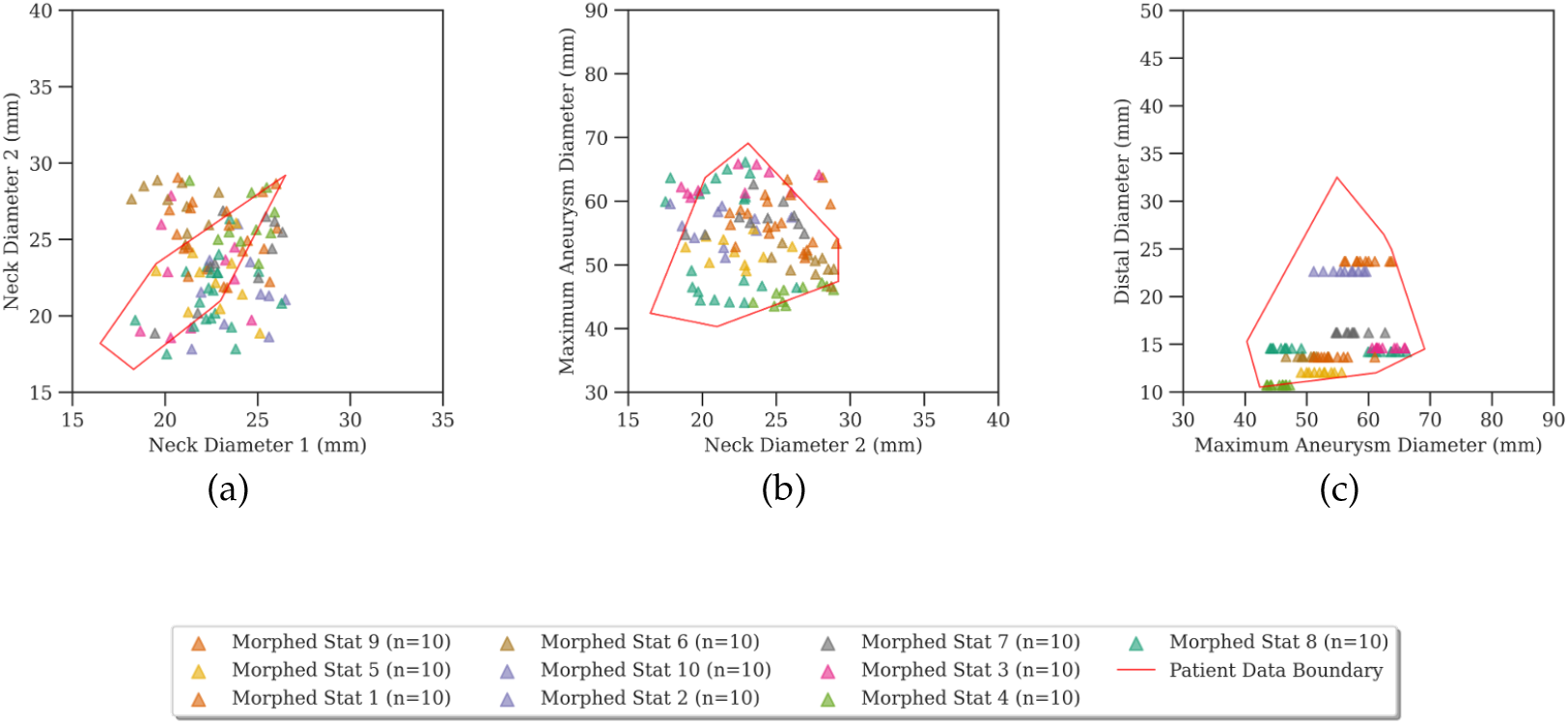
Parameter space comparison between patient data and morphed geometries for female patients aged 70-79. The convex hull of patient data (red) is overlaid with morphed geometry points. Subplots represent relationships between neck diameter points, neck-to-aneurysm transitions, and aneurysm-to-distal transitions. The legend in the second row corresponds to this specific age group.

**Figure 7.**
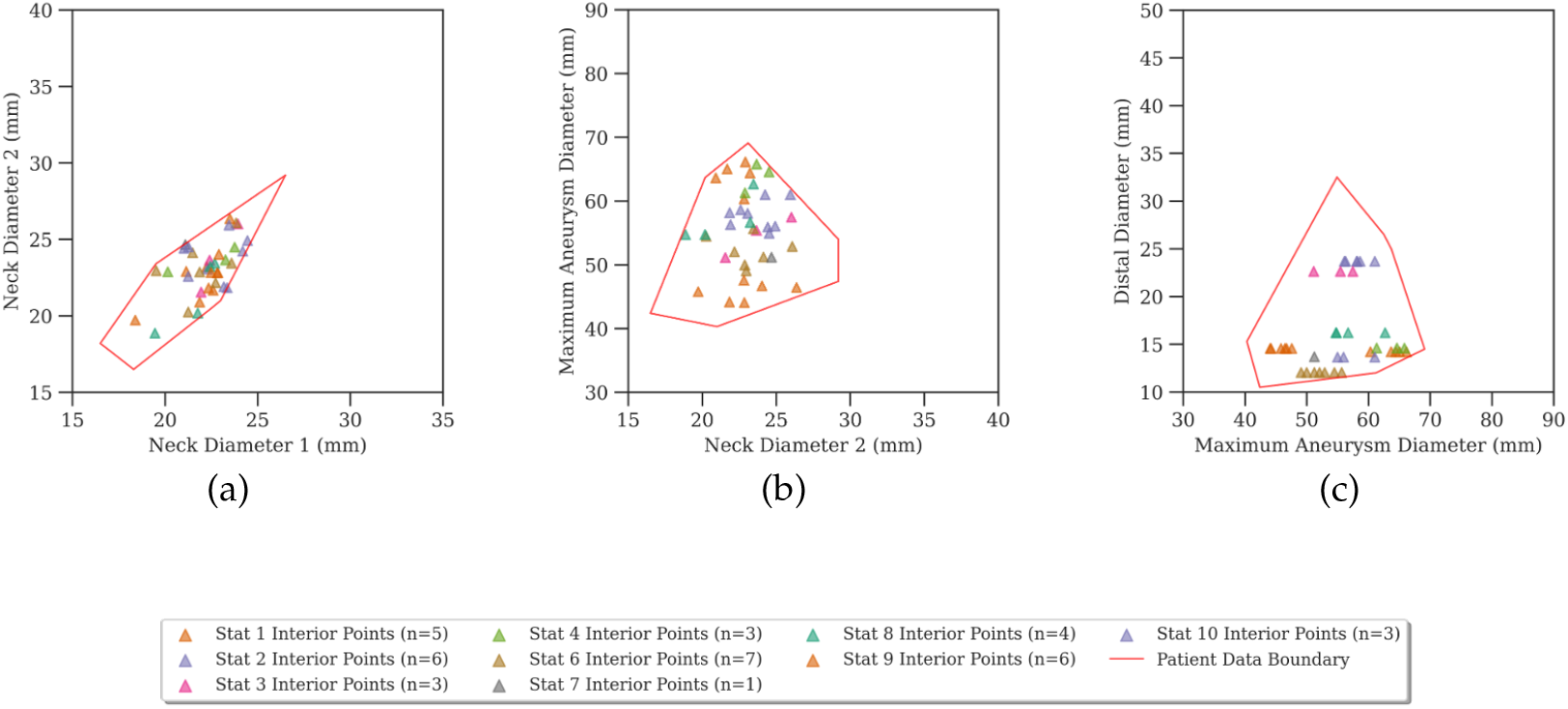
Universal interior set analysis for female patients aged 70-79. Valid morphed geometries are shown within the convex hull of patient data (a) Neck Diameter 1 vs. Neck Diameter 2, (b) Neck Diameter 2 vs. Maximum Aneurysm Diameter, (c) Maximum Aneurysm Diameter vs. Distal Diameter.

Across this age–gender group, 65% of morphed geometries were interior to at least one convex hull, and 18.2% satisfied the universal interior criterion. Overall, 400 geometries were generated and 182 high confidence geometries were selected for downstream CFD simulations to ensure physiological relevance. This ensures pipeline yield of approximately ∼ 45%. Additional plots for other demographic groups are provided in Appendix J(c).

### 3(f) Geometry–Hemodynamics Relationship Analysis

Figure 8 maps the Pearson correlations between eleven geometric descriptors and six haemodynamic biomarkers across 364 simulations. Statistical significance was evaluated using the p-value corresponding to each correlation coefficient, which tests the null hypothesis of zero correlation. The following thresholds were used to indicate levels of significance: ∗ for *p*< 0.05, ∗∗ for *p*< 0.01, and ∗∗∗ for *p*< 0.001.

**Figure 8.**
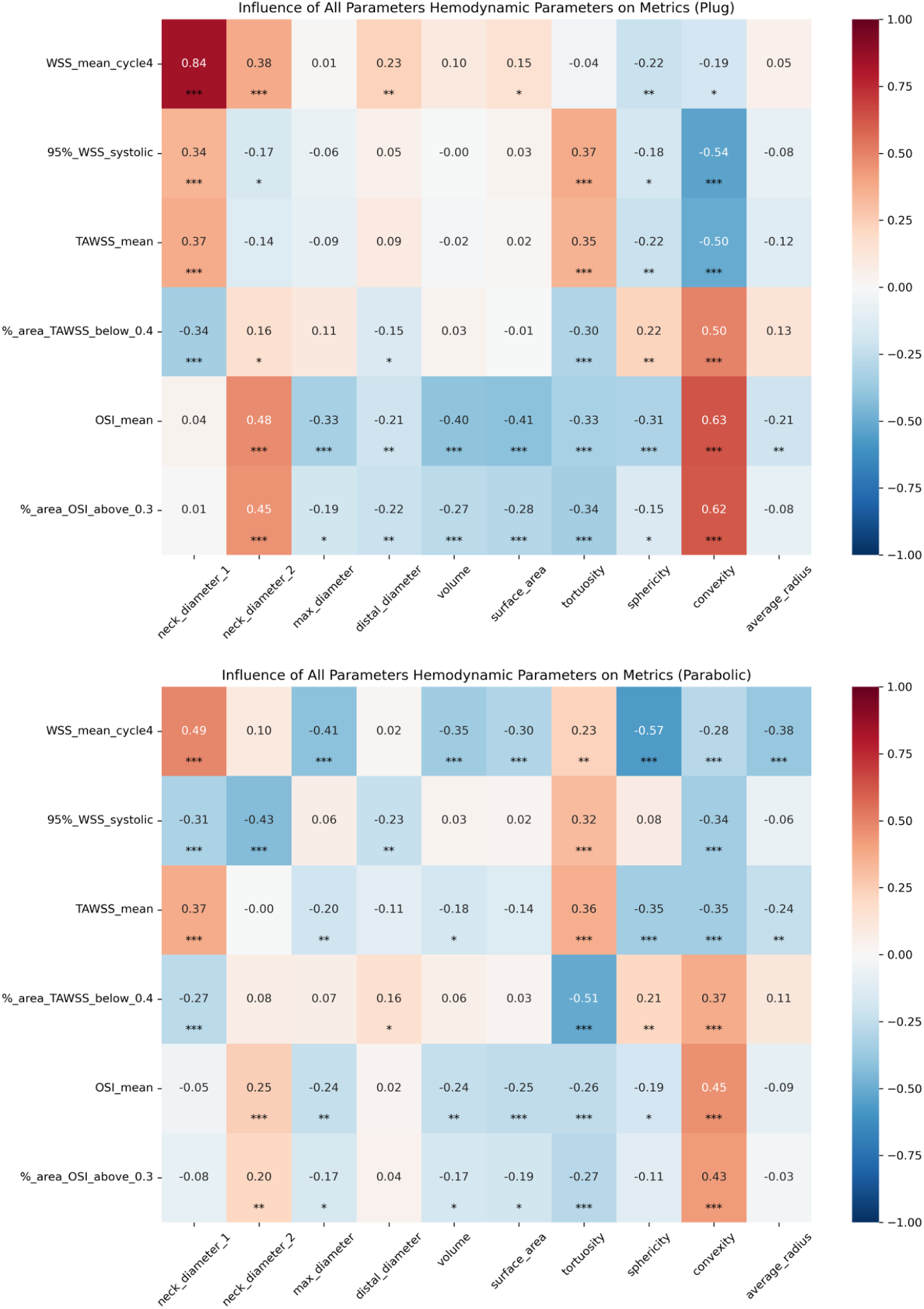
Correlation heat map showing relationships between geometric parameters and haemodynamic metrics using a plug/uniform (top) and parabolic (bottom) velocity inlet profile. Colour intensity indicates correlation strength. Asterisks denote statistical significance.

#### (3(f)(i)) Neck diameters

A widening of neck diameter 1 is the strongest positive modulator of wall shear magnitude. It raises WSS_mean_ (*r* = 0.49, *p*< 0.001), while simultaneously reducing the surface exposed to pro-thrombotic low shear, quantified here as the percentage of wall area where TAWSS < 0.4, Pa (*r* = −0.27, *p*< 0.001). At the same time, it is associated with the modest decrease of WSS_0.95_ (*r* = −0.31, *p*< 0.001).

Similarly, neck diameter 2 is also associated with a reduction in peak shear WSS_0.95_ (*r* = −0.43, *p*< 0.001). However, it correlates positively with the markers of oscillatory flow, including both OSI_mean_(*r* = +0.25) and the percentage of the wall area exposed to high OSI (%area OSI > 0.3, *r* = +0.20).

#### (3(f)(ii)) Global size metrics

Although maximum diameter is the standard clinical trigger for elective repair, the results show that it has little effect on peak WSS (*r* = +0.06, *p*> 0.05). Paradoxically, however, the same maximum diameter is negatively correlated with the mean wall shear WSS_mean_ (*r* = −0.41, *p*< 0.001).

Volume and surface area show weak, non-significant correlations with low TAWSS exposure (|*r*| ≤ 0.06). Average radius shows a negative correlation with TAWSS_mean_, meaning that wider vessels tend to have lower mean cyclic shear. As like maximum diameter, volume, surface area and average radius all are negatively correlated with WSS_mean_.

When the distal diameter is wider, it does not affect overall shear levels across the cardiac cycle, but it reduces shear at peak moments. Specifically, it reduces the upper end of wall shear stress, WSS_0.95_ decreases (*r* = −0.23, *p*< 0.01). However, it has no meaningful effect on WSS_mean_ or OSI.

#### (3(f)(iii)) Shape compactness

Shape compactness was quantified by sphericity (Equation (2.11)) and convexity (Equation (2.12)) indices. Sphericity is observed as the strongest modulator for mean wall stress (*r* = −0.57, *p*< 0.0001). Both sphericity and convexity show the same correlation for TAWSS_mean_ (*r* = −0.35, *p*< 0.001). Oscillatory biomarkers, however, are driven most strongly by the convexity, which is positively correlated with both OSI_mean_ (*r* = +0.46, *p*< 0.001) and the wall area exposed to high OSI (*r* = +0.43, *p*< 0.001).

#### (3(f)(iv)) Centreline tortuosity

Centerline toruosity is significantly correlated with every haemodynamic metric examined, reaching *p*< 0.001 except for WSS_mean_ (*p*< 0.01). The area associated with the low wall shear region shows the strongest correlation with tortuosity (*r* = −0.51). Tortuosity is also inversely associated with oscillatory shear (*r* = −0.26).

#### (3(f)(v)) Sensitivity to inlet profile

One of the most important mechanistic effect on the results relates to the choice of inlet velocity profile and it’s influence on shear metrics. Across the parameter space, plug-flow simulations weaken these correlations. For instance, the correlation between WSS_mean_ and neck diameter 1 drops from *r* = 0.84 to *r* = 0.49. Despite this sensitivity, two key findings hold true across both inlet types: neck diameters along with compactness matrices shows strong correlations with haemodynamic biomarkers.

**Figure 9.**
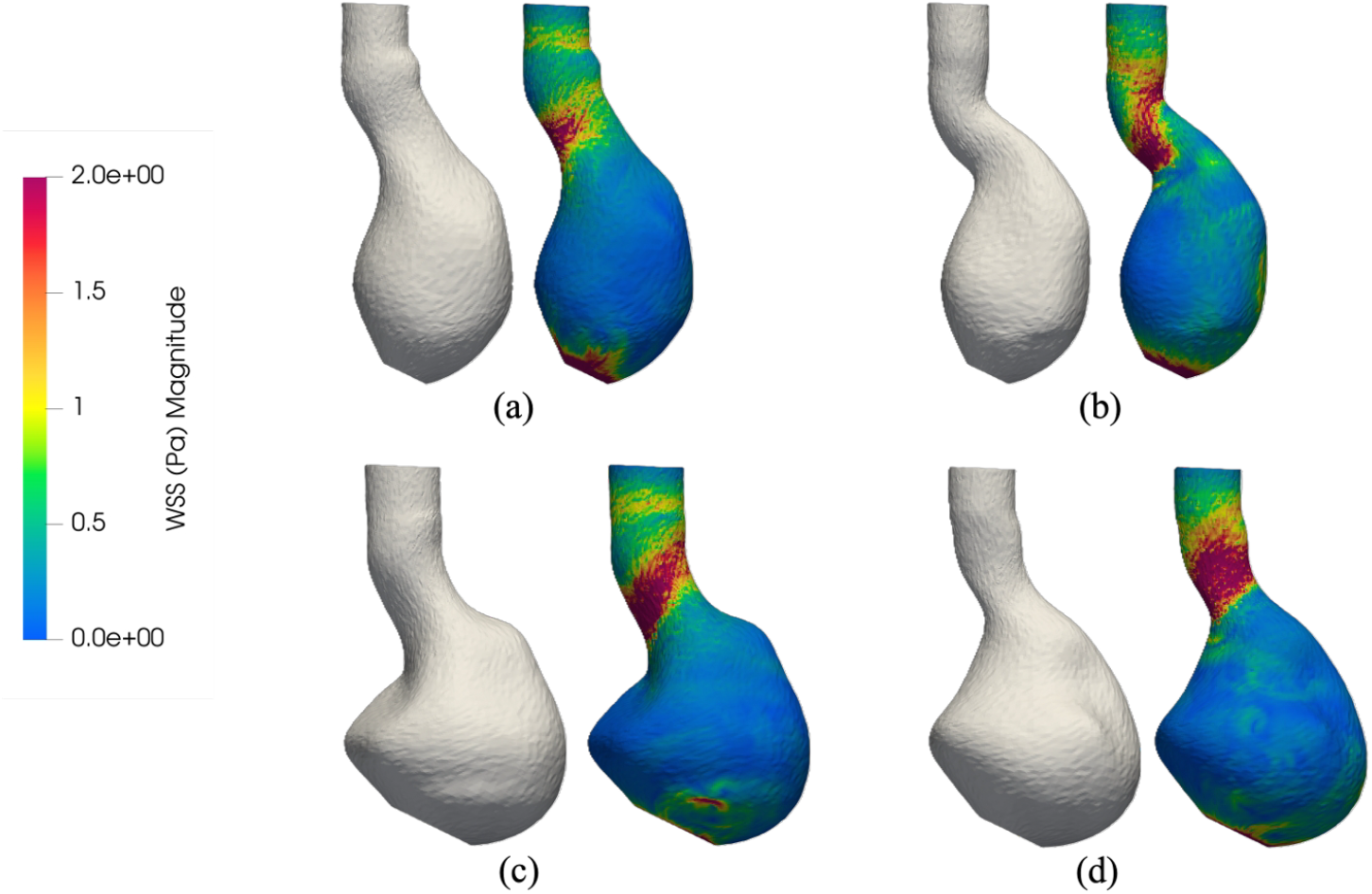
Representative AAA geometries (grey surface, left of each pair) and peak-systolic WSS magnitude contours (right of each pair) for four demographic groups: (a) male, 60-69 year; (b) male, 70-79 year; (c) male, 80+ year; (d) female, 70-79 year. The characteristic low-shear recirculation zone (WSS < 0.4 Pa, blue) dominates the sac in all cases, while the proximal neck sustains elevated shear (>1.5 Pa, red-pink) due to the inflow jet. Panel (c) exhibits a secondary high-shear region at the distal sac, attributable to the locally narrowed geometry at the iliac transition.

## 4. Discussions

This work introduces a constraint-aware automated synthetic cohort generator for AAA that enables scalable in-silico haemodynamics. We demonstrate, for the first time at the cohort scale, that neck diameters and sphericity are the dominant modulator of WSS, while convexity is the leading modulator of oscillatory shear, suppress OSI, and maximum diameter primarily lowers mean wall shear rather than peak stress. These insights provide in this work suggest risk models should move beyond maximum neck diameter, i.e. *D*_max_, to include neck geometry and compactness.

### 4(a) Interpretation of findings

Before interpreting the geometry–haemodynamics correlations, we note that all haemodynamic outputs fell within physiologically reported ranges from patient-specific AAA CFD and *in-vitro* studies [82–84], providing face validity for the simulation framework. The dominance of neck diameters as a predictor of wall shear magnitude is consistent with patient-specific CFD results reported by Pantoja [82] and the longitudinal reconstructions of Bappoo et al. [83], and can be explained by a longer coherent jet that convects high-momentum blood directly onto the sac wall.

Maximum diameter, despite being the standard clinical trigger for elective repair, had little effect on peak wall shear stress; once flow separates into the sac, the wide cross-section acts like a diffuser, slowing and spreading the jet without increasing local stress. As a result, further enlargement simply displaces the recirculation zone downstream. This is consistent with the diameter-peak wall shear stress reported by Vilalta-Alonso et al. [85]. Paradoxically, however, the same maximum diameter is negatively correlated with mean shear values, consistent with CFD studies showing that sac enlargement primarily increases low shear regions [82, 83]. In other words, larger aneurysms do not increase peak shear forces, but they lowers the overall shear which is linked to thrombus formation and inflammatory remodelling. The distal diameter reduces shear at peak systole (WSS_0.95_, *r* = −0.23, *p*< 0.01) without much affecting other metrics.

Shape compactness, neck diameters, and centreline tortuosity all act primarily on both oscillatory rather and magnitude-based metrics. Sphericity was inversely related to mean OSI (*r* = −0.19), while convexity was positively related to it (*r* = +0.46), implying that spherical sacs experience lower shear and less oscillation, whereas highly convex sacs combine lower mean shear with more oscillatory flow. Flow physics considerations explain these findings: compact sacs spread the inflow jet over a larger area, reducing local impingement and allowing vortices to be trapped centrally rather than near the wall. Elongated sacs behave like curved tubes; they concentrate the jet against the wall, increasing shear. This general trend that shape modulates flow is also observed in the literature from CFD analysis of patient-specific AAAs [85, 86]. Neck diameter 2 follows the same family: its positive correlations with mean and high-area OSI (*r* = +0.25, *r* = +0.20) suggest that, while cycle-averaged shear forces remain largely unaffected, disturbed flow becomes more prominent as neck diameter 2 expands, possibly due to more persistent flow separation and oscillatory vortices. Centreline tortuosity raises shear magnitude but is inversely related to OSI (*r* = −0.26) and this runs counter to the findings of Wang et al. [84], who reported higher OSI with increasing tortuosity. It also shows the strong association with a reduction in low shear surface area. Combined with its positive correlation with shear magnitude, this indicates that increasing tortuosity shifts the sac away from the low-shear conditions associated with thrombus formation and expansion.

A final mechanistic observation concerns the inlet velocity profile. The correlation magnitudes were sensitive to the inlet profile, generally strengthening under plug flow. A similar inlet-profile dependence was reported by Wang et al. [87]. Importantly, the identity of the dominant predictors is preserved across both inlet conditions: neck diameters and the compactness descriptors (sphericity, convexity) retain their leading roles, lending robustness to the clinical interpretation of the geometry–haemodynamics relationships reported here.

### 4(b) Clinical implications

This study offers practical insights for usage of CFD simulations to enable personalised AAA risk assessment and treatment planning. The study demonstrates that neck diameters and shape compactness emerges as a strong predictor of both WSS and OSI parameters. This suggests that monitoring neck growth and overall bulge could provide earlier warnings as compared to current practice of monitoring maximum sac diameter alone. Prospective validation against rupture-event endpoints would be required before any such monitoring strategy could be recommended for clinical protocols.

Shape descriptors, particularly sphericity and convexity, were strongly associated with disturbed flow, while increased tortuosity raised WSS. From a clinical perspective, maximum and neck diameters are routinely available from 2D imaging, while convexity may be reasonably approximated from 2D cross-sectional contours, with sphericity and tortuosity typically requiring full 3D reconstruction. These indices can be computed from standard CTA and could complement diameter measurements in future risk scores for AAA. It was particularly noted that the maximum diameter correlated primarily with reduced mean wall shear rather than peak stresses, implying that relying on diameter alone may overlook patients with a low-shear haemodynamic environment at risk of intraluminal thrombus (ILT) formation or progressive degeneration.

By reproducing reported demographic trends, notably, maximum diameter increasing with age and female aneurysms being smaller across parameters – the synthetic cohort highlights the need for age- and gender-specific risk stratification and device design. Female AAA patients in particular exhibit smaller, shorter, and more angulated proximal necks than male patients and have historically poorer EVAR outcomes [26], yet remain under-represented in the clinical-trial datasets on which device sizing and fixation behaviour are typically validated. Gender- and age-stratified synthetic cohorts produced by the present framework are therefore well placed for in-silico evaluation of endograft sealing-zone, fixation, and migration behaviour on under-represented anatomies, in line with the credibility framework set out in ASME V&V40 [88] for context-of-use-driven medical device evaluation. Additionally, the identification of shape compactness (high sphericity and convexity) as a suppressor of oscillatory shear suggests that compact shapes may confer haemodynamic benefits, whereas more tortuous or irregular shapes may accelerate degeneration.

### 4(c) Limitations and future work

The synthetic cohort is shaped by its source data from the Kyriakou et al. [27] dataset. This dataset covers EVAR-planning anatomy, so that the fitted diameter distributions characterise intervention-eligible cases and, hence, should not be extrapolated to surveillance-stage aneurysms or non-EVAR morphologies. The dataset is also male-dominated, and the synthetic cohort inherits that bias. Thus, expanding the training data with female populations is essential for broader generalisability.

A second limitation remains that the aspect of ILT, which is often present in the majority of clinical AAA cases. In the CFD domain considered in this work, the ILT is neglected owing to the source diameters, taken from Kyriakou et al. [27], being measured to the intima. Thus, the synthetic lumen may overestimate the blood-flowing volume where ILT would otherwise reside and hence potentially underestimating WSS magnitudes while overestimating the spatial extent of low-shear regions. Further, the diameter-measurement uncertainty is not formally propagated through the pipeline and a dedicated sensitivity analysis, together with the uncertainty on the tortuosity, is identified as valuable future work.

At the geometry level, the base centreline is held fixed across the cohort. Inter-patient variation in axial curvature and tortuosity is therefore not represented as a sampling dimension, and the geometry-flow correlations reported here may shift once centreline variability is introduced. Per-case centreline substitution requires no structural modification to the pipeline and is a priority extension. Beyond centreline variability, the surface sweep self-intersection constraint additionally precludes synthesis of highly angulated morphologies. Direct case level validation, comparing individual synthetic geometries against manually segmented patient anatomies, is structurally inapplicable to the present framework. The geometry population is entirely synthetic and each geometry is generated by morphing a base geometry using parameter combinations drawn from a patient-derived distribution, so no case corresponds to a specific patient anatomy.

On the CFD side, a single inlet velocity waveform was applied across all cases, and the Newtonian rheology assumption, although standard in large-vessel aortic CFD, may underestimate shear-thinning effects in recirculation zones. Case-level CFD-to-MRI validation was not possible for the same reason as direct geometric validation and hence the synthetic geometries are not matched to individual patient scans. Future work should therefore focus on population-level validation, for example by comparing the distributions of TAWSS, OSI, and WSS with population-level 4D-flow MRI measurements.

Finally, the correlation analysis itself has structural limits. The analysis relies on Pearson coefficients and captures only linear relationships; the inherent collinearity among size-related metrics 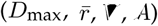 was not explicitly addressed. Future work should apply global sensitivity analysis and non-linear methods to uncover more complex geometry–haemodynamics interactions.

The presented framework provides a pre-cursor for a potential automated image-to-simulation pipeline that can generate large libraries of demographically stratified AAA geometries. Such a dataset can enable systematic training of machine-learning models to provide geometry–flow correlations much faster. Combining this synthetic topology generator with deep-learning surrogates for rapid WSS estimation and generative models for anatomy synthesis could yield comprehensive, real-time pipelines that balance accuracy with computational and operational efficiency.

## Supporting information

Electronic supplementary materials

## Acknowledgment

The authors acknowledge the University of Manchester for financial support via the Dean’s Scholarship (Vijay Nandurdikar).

## Ethics statement

This study is an entirely in-silico, secondary computational analysis of the previously published, anonymised computed-tomography angiography dataset of. No new human or animal data were collected for this work, and no identifiable patient information was accessed. Accordingly, no additional ethics approval was required.

## Credit authorship contribution statement

**Vijay Nandurdikar:** Conceptualization, Data Curation, Formal analysis, Investigation, Methodology, Resources, Software, Validation, Visualization, Writing - Original draft; **Aryan Tyagi:** Software, Formal analysis; **Tejas Canchi:** Methodology, Writing - Review & Editing; **Alistair Revell:** Supervision, Writing - Review & Editing; **Alejandro F. Frangi:** Supervision, Writing - Review & Editing; **Ajay B Harish:** Conceptualization, Funding acquisition, Formal analysis, Investigation, Methodology, Project administration, Supervision, Software, Validation, Resources, Writing - Original draft.

## Data availability statement

Data and relevant code for this research work are stored in GitHub at https://github.com/Harish-Research-Lab/Synthetic-AAA-CFD-framework and have been archived within the Zenodo repository at https://zenodo.org/records/21435232.

## A. Configuration Parameters

**Table 4.**
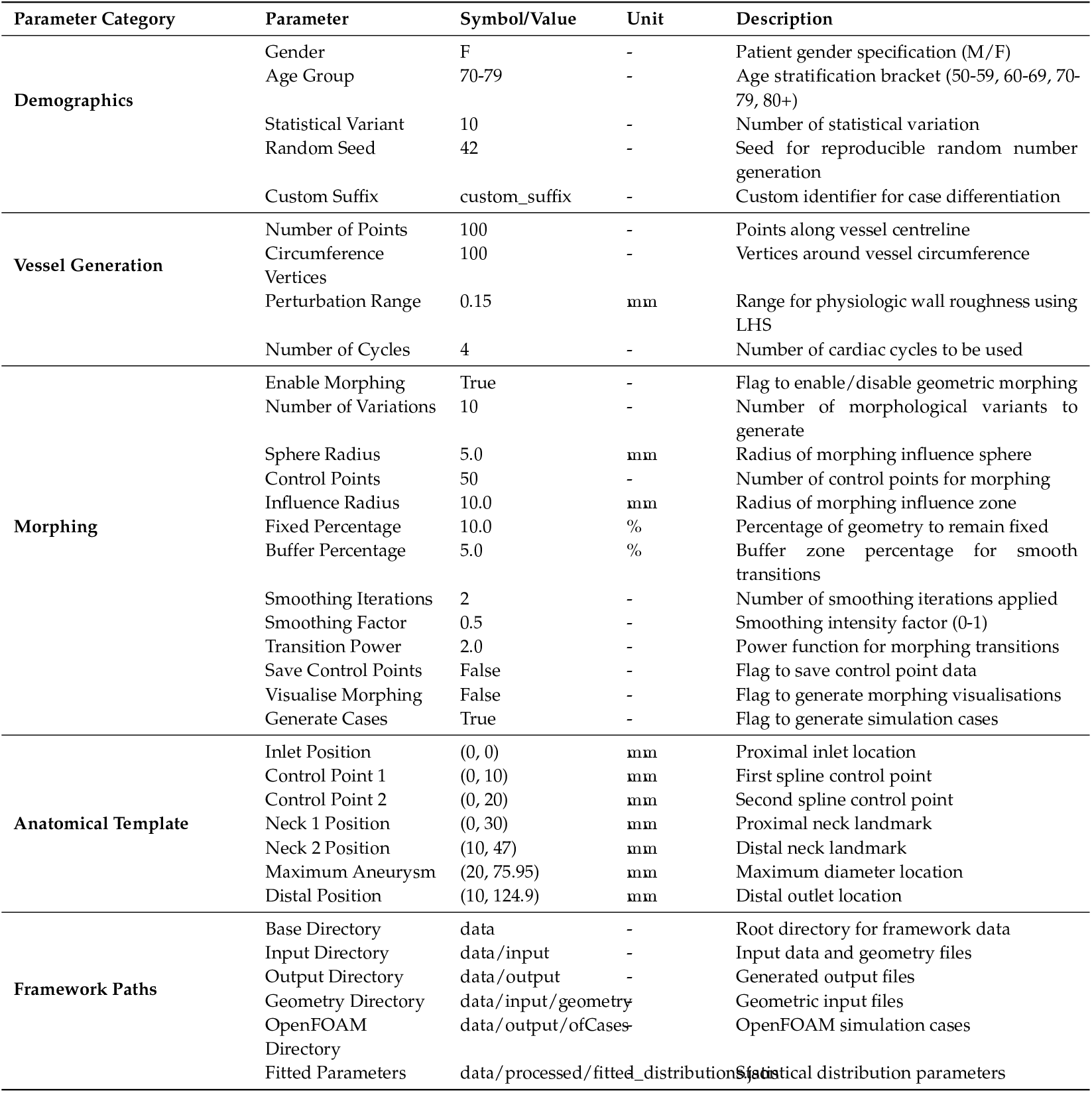
Complete configuration parameters for AAA geometry generation framework.

## B. OpenFOAM^®^ case generation

The transition from geometric models to CFD simulations requires the automated generation of case directories with appropriate boundary conditions and mesh specifications. A pipeline was developed to convert AAA geometries into simulation-ready OpenFOAM cases.

### B(a) Case structure

Each case follows a standardized naming convention that encodes demographic and morphological characteristics:

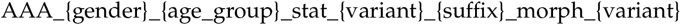

This is useful for tracking geometric variations across demographic cohorts while maintaining organized datasets. For every statistical variant and its associated morphed geometry, the framework builds a OpenFOAM case directory including a dedicated parameters subfolder that records all geometric inputs and configuration settings. The base case configuration employs template that are dynamically populated with case-specific parameters derived from the geometric analysis.

### B(b) Mesh processing

A background Cartesian grid is generated with `blockMesh` with appropriate padding; and then mesh generation is done with snappyHexMesh.

Prior to meshing, the surface is decomposed into `inlet, outlet`, and `wall` STL regions. Surface normals are computed, and a single interior seed point is automatically determined.

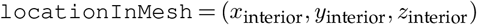

where (*x*_interior_, *y*_interior_, *z*_interior_) is the automatically detected point inside the vessel lumen. The same point is passed to snappyHexMeshDict to ensure proper domain marking during the mesh refinement process. Geometry-specific parameters, including mesh refinement controls, are adapted to each model according to geometry dimensions.

### B(c) Inlet velocity profile specification

The simulation framework accepts any prescribed scalar velocity waveform as a two-column CSV file (*t, u*) and converts it into a vector field aligned with the inlet surface normal. The time-dependent inlet velocity is therefore

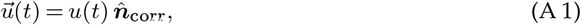

where *u*(*t*) is the waveform magnitude and 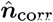 is the corrected unit normal.

Surface normals 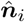 are computed for every inlet triangle using cross-products of STL face edges. An inlet reference normal is computed by area-weighted averaging of these face normals and normalised to unit length. Orientation is validated using a dot-product test with the interior direction vector 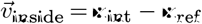, where **x**_ref_ lies on the inlet and **x**_int_ is the same interior point used in mesh generation. If 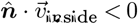, the normal is flipped.

Element-wise multiplication of the scalar waveform with the corrected normal yields the three-component velocity profile:

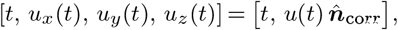

which enforces proper inflow direction while preserving the intended velocity magnitude. The resulting vector profile is exported in a format compatible with OpenFOAM.

Simulation consistency is maintained by enforcing uniform settings for solver tolerances, discretisation schemes, and time stepping across all cases. This ensures that differences in simulation output arise solely from geometric variation, not solver configuration.

## C. Centreline-sweep surface construction

The abdominal aortic surface is generated by sweeping circular cross-sections along the centreline using a parallel-transport frame.

At each centreline station *s*_*i*_, the local unit tangent 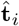 is estimated from the finite-difference of adjacent stations. An initial orthonormal pair 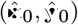, both perpendicular to 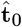, is established at the inlet via a null-space construction.

At each subsequent station the frame is updated iteratively

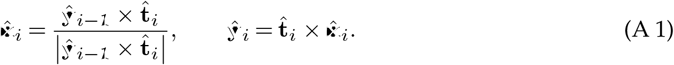

This parallel-transport update propagates the frame without accumulating twist, so cross-sections remain consistently oriented even along curved centreline paths. Vertices on the *i*-th cross-section of radius *r*_*i*_ = *D*_*i*_/2 are placed at

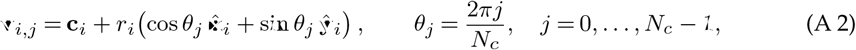

where **c**_*i*_ is the *i*-th centreline point and *N*_*c*_ is the number of circumferential vertices. Adjacent rings are connected by triangles to form the closed wall surface.

### Geometric validity condition

The construction produces a non-self-intersecting surface if and only if the local centreline radius of curvature exceeds the local vessel radius at every arc-length station

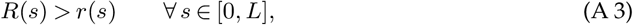

where *L* is the total centreline arc length. If this condition is violated at any station, adjacent cross-sectional rings overlap and the surface self-intersects.

### Capability illustration

The framework generates angulated and tortuous centreline paths by assigning non-zero lateral offsets to the centreline control points. Figure 10 shows an S-shaped geometry produced by prescribing a sinusoidal lateral offset profile over the full vessel centreline length (*L* = 125 mm, amplitude 28 mm, one full period).

**Figure 10.**
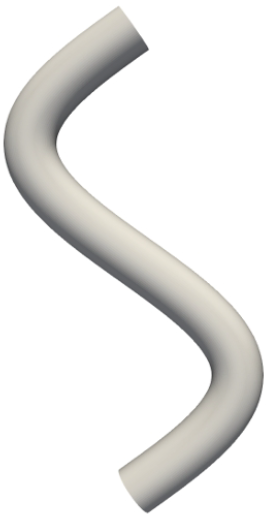
S-shaped capability geometry. The smooth, artefact-free surface confirms no self-intersection (minimum *R*(*s*)/*r*(*s*)= 2.02, well above the threshold of 1).

The smooth, non-self-intersecting surface confirms that the parallel-transport sweep handles extreme multi-directional curvature robustly. This geometry serves as a capability demonstration and is not a clinical AAA.

## D. Cycle-to-cycle convergence of the inlet pressure waveform

Since the inlet velocity waveform is prescribed as a boundary condition, the computed area-averaged inlet pressure is a free quantity that integrates the global hydrodynamic state of the domain. Its cycle-to-cycle variation therefore provides a sensitive diagnostic for periodic steady-state convergence. Figure 11 shows the area-averaged inlet pressure waveform across all four simulated cardiac cycles for a representative case. Cycles 1 and 2 show observable departure from the converged waveform, most prominently during the diastolic deceleration phase (0.35-0.45 s). Cycles 3 and 4 are visually indistinguishable throughout the full waveform; quantitatively, the maximum absolute difference between cycles 3 and 4 is 1.41 Pa, corresponding to a relative difference of 0.34% of the mean waveform amplitude, well within the conventional < 1% convergence criterion.

**Figure 11.**
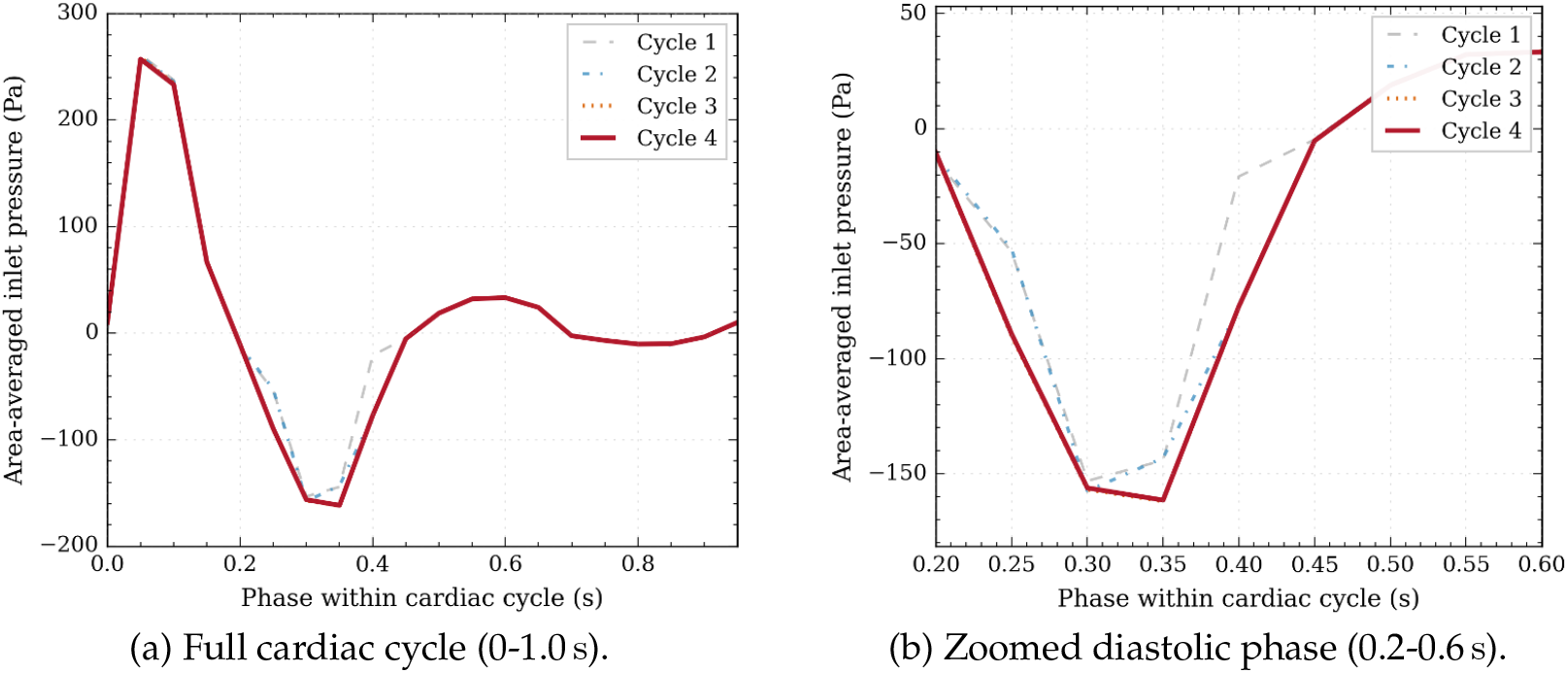
Cycle-to-cycle convergence of the area-averaged inlet pressure waveform for a representative case (male, 60-69, median geometry). (a) Full cardiac cycle across all four simulated cycles. (b) Zoomed view of the diastolic deceleration phase (0.2-0.6 s) where transient effects are most prominent.

## E. Mesh independence assessment

A synthetic AAA model representing a 60–69-year old male was discretised at five successive grid levels with blockMesh and snappyHexMesh. The cell count *N* approximately doubled from one level to the next. All meshes share the same boundary layer topology: five prismatic wall layers grown from the lumen surface (first-layer thickness 0.3 mm, expansion ratio 1.1), yielding *y*^+^≲1 on the adopted grid. This keeps near-wall resolution constant so that only the core flow cells change with refinement.

Computational cost rose nearly linearly with cell count: the finest mesh (7.72 × 10^6^ cells) required 17h 18 min for four cardiac cycles, whereas the adopted mesh (1.29 × 10^6^ cells) finished in 2h 26 min, about a seven times faster and also keeping the centre line velocity error below 2.3% (Table 5).

Figure 12 shows that the peak-systolic centreline velocity curves for all five grids are virtually indistinguishable at the inlet and outlet; the only noticeable deviations occur inside the aneurysm bulge (25<*z*<40 mm), where the coarsest mesh overestimate velocity by up to ∼0.05 m s^*−*1^. Those differences vanish once the cell count exceeds 1.29 × 10^6^. Figure 13 zooms in on the adopted mesh versus the finest mesh and adds a 95% bootstrap confidence interval. Richardson extrapolation with the three finest grids (*N* = 2.75, 4.68, 7.72 M) yields an observed order of accuracy *p* = 1.94 and a fine-grid GCI of 2.2%.

**Figure 12.**
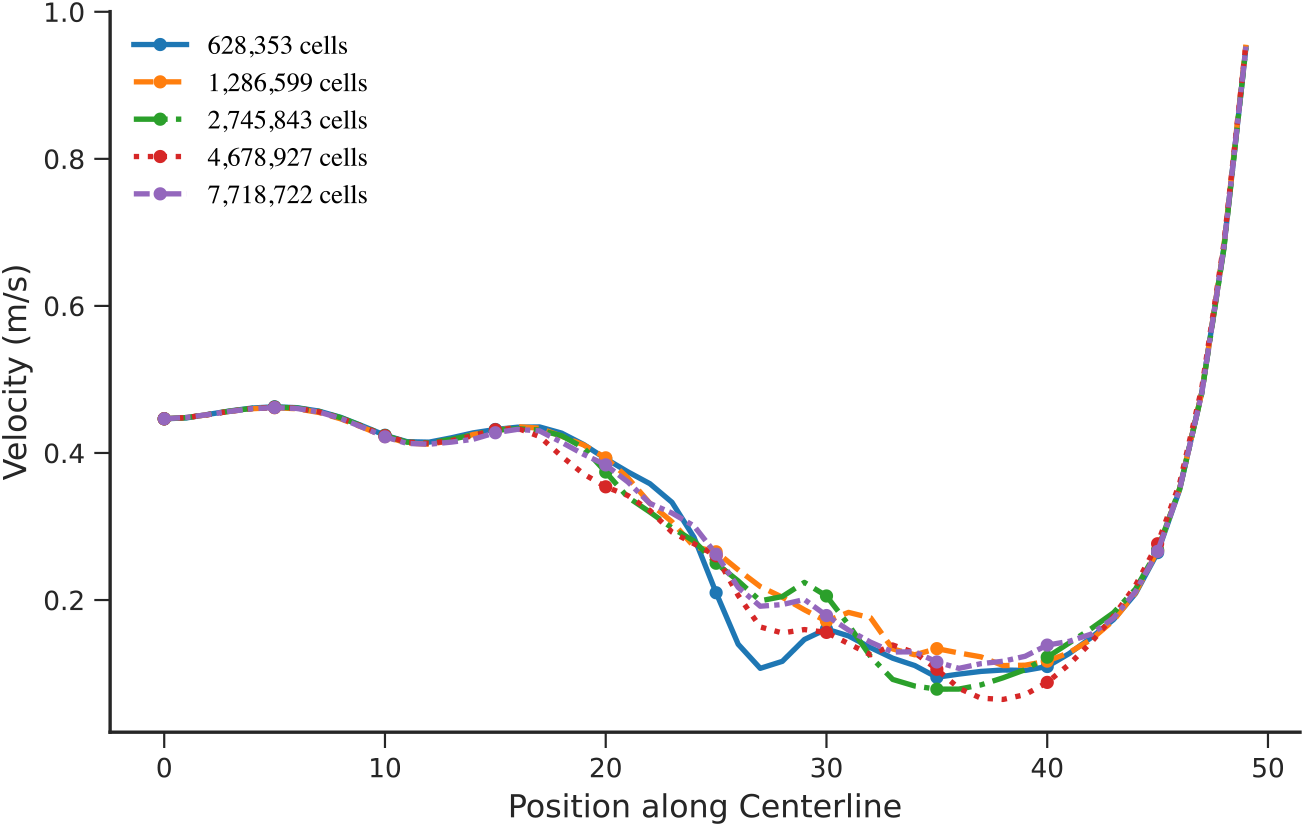
Centreline velocity for all five mesh levels. Upstream of the aneurysm (*z* < 15 mm) and downstream (*z* > 45 mm) the meshes collapse onto a single curve. The only mesh-sensitive region is the aneurysm sac (25<*z*<40 mm), where the coarsest grid depart from the converged solution. These differences disappear for *N* ≥ 1.29 × 10^6^.

**Figure 13.**
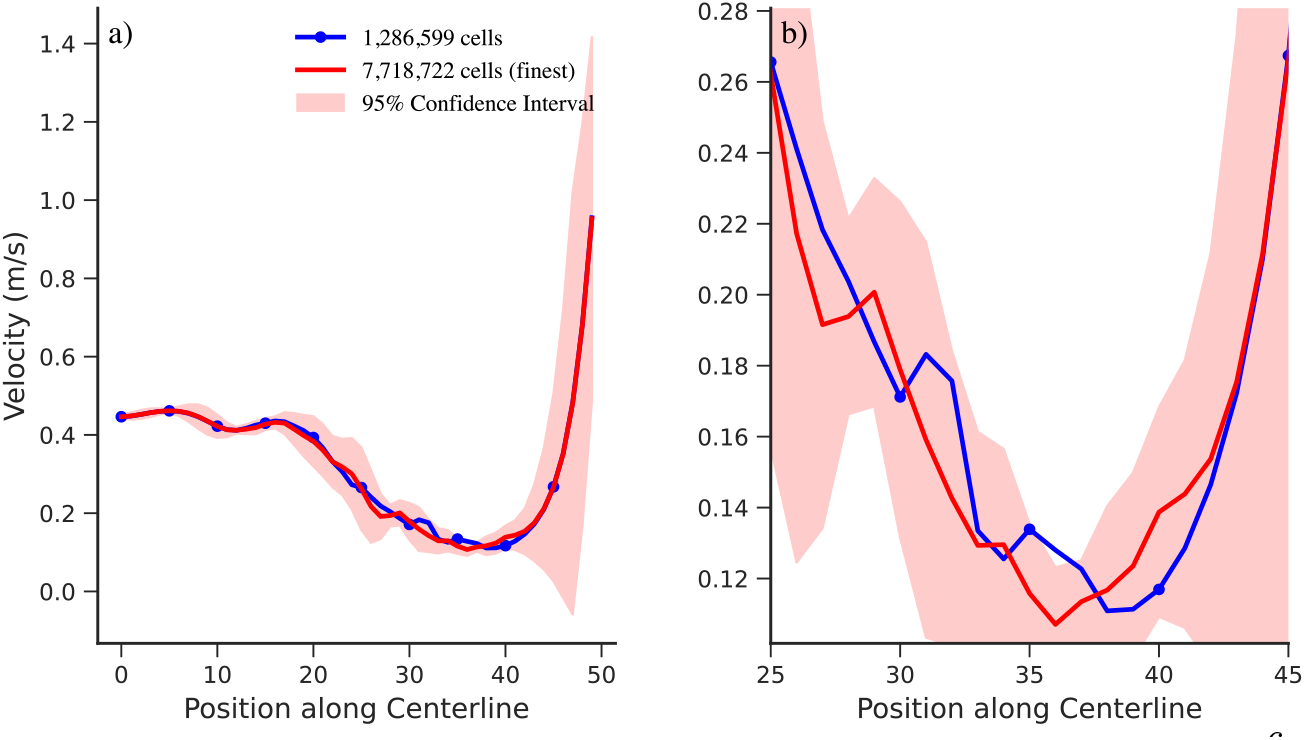
Mesh convergence of centreline velocity for the adopted mesh (1.29 × 10^6^ cells, blue) vs. the finest mesh (7.72 × 10^6^ cells, red). Shaded band: 95% bootstrap confidence interval on the finest mesh.

Selecting the mesh parameters associated with the 1.29 M-cell mesh therefore reduces the run time without compromising accuracy, and it is used for all rest of the simulations.

### Algorithm 1

Virtual AAA cohort generation framework

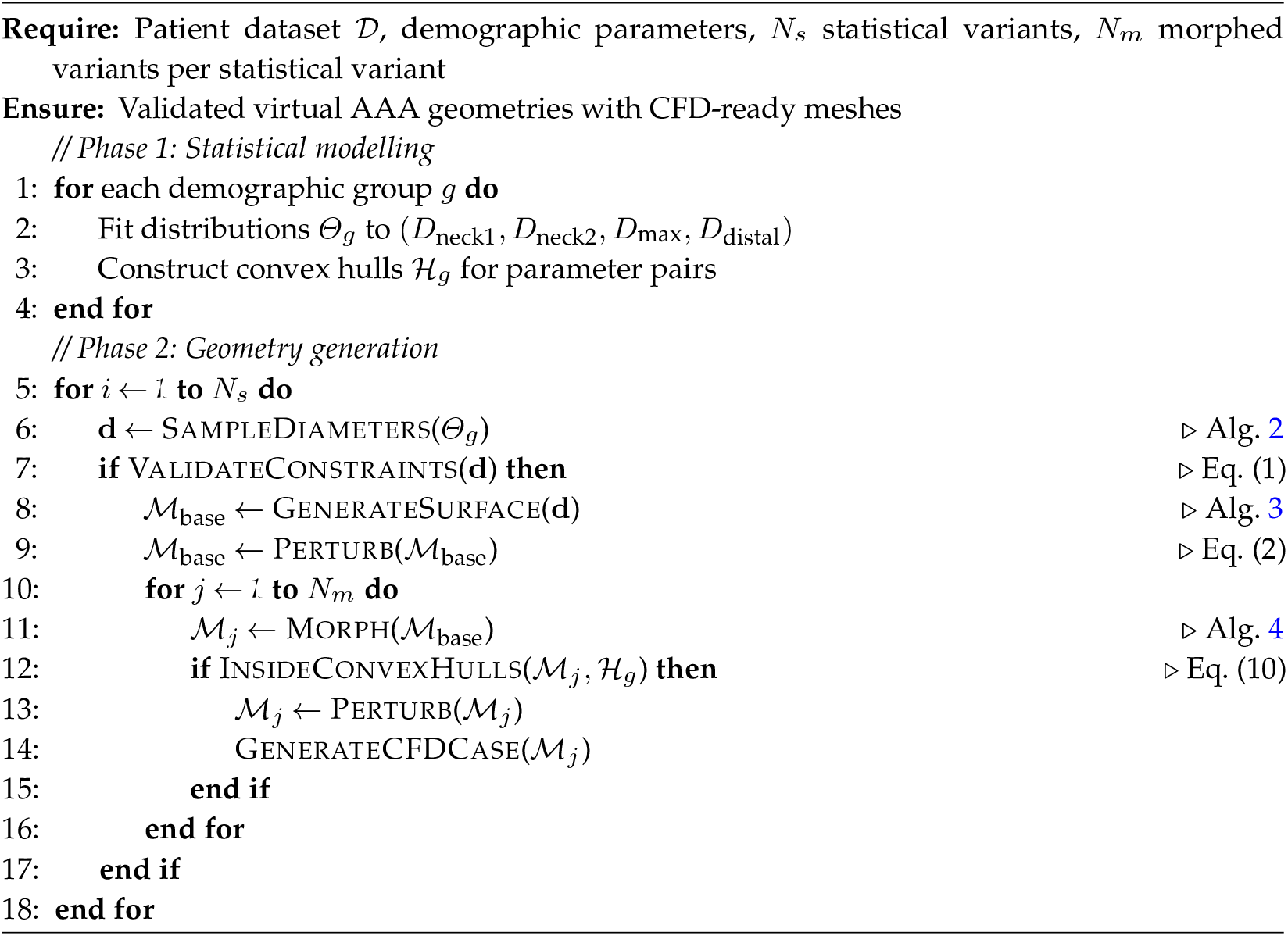

### Algorithm 2

Rejection-based diameter sampling

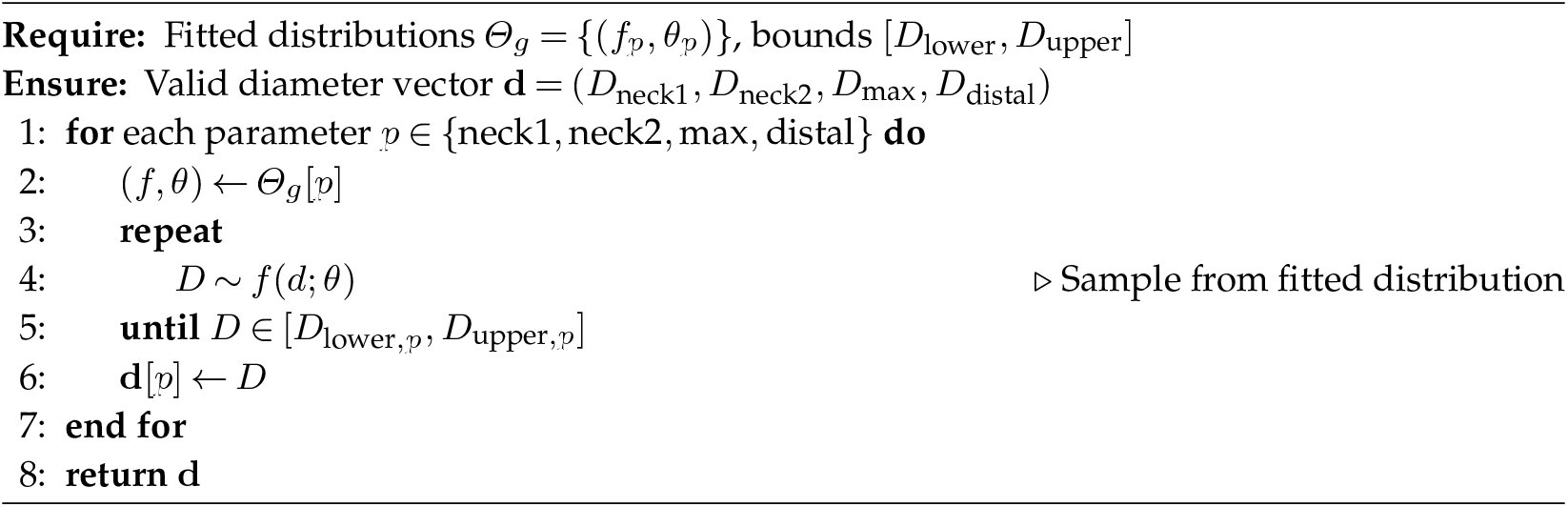

**Table 5.** Wall-clock time for four cardiac cycles on each mesh (32 cores).

|  |  |  |  |  |  |
| --- | --- | --- | --- | --- | --- |
| Cells ($N$) | 628 353 | 1 286 599 | 2 745 843 | 4 678 927 | 7 718 722 |
| Time (h:mm:ss) | 01:08:13 | 02:26:27 | 06:04:46 | 09:42:23 | 17:18:06 |

## F. Algorithm Pseudocode

## G. Distribution Fitting Details

### Algorithm 3

Base geometry and surface mesh generation

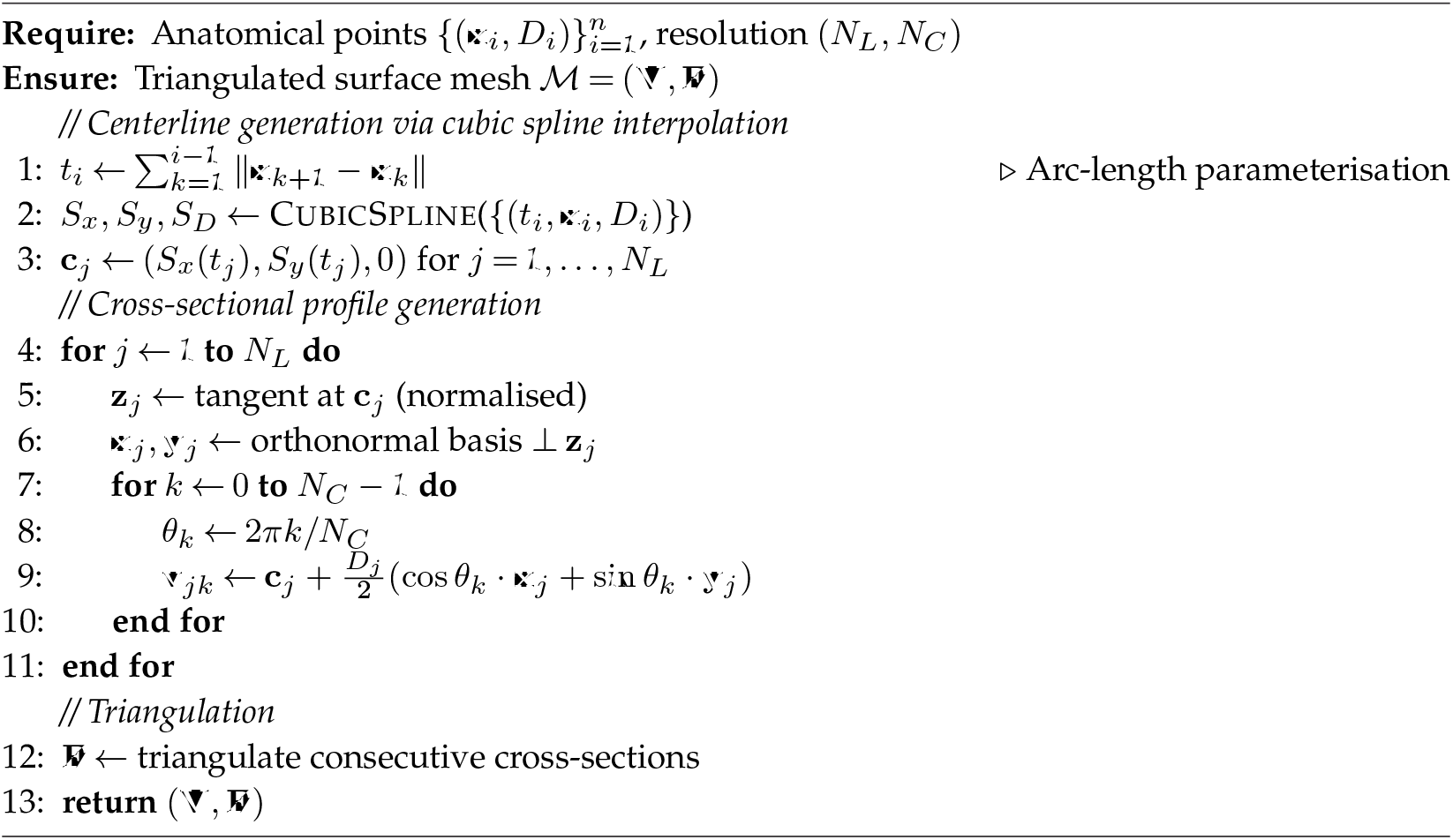

### G(a) Maximum Aneurysm Diameter

**Table 6.**
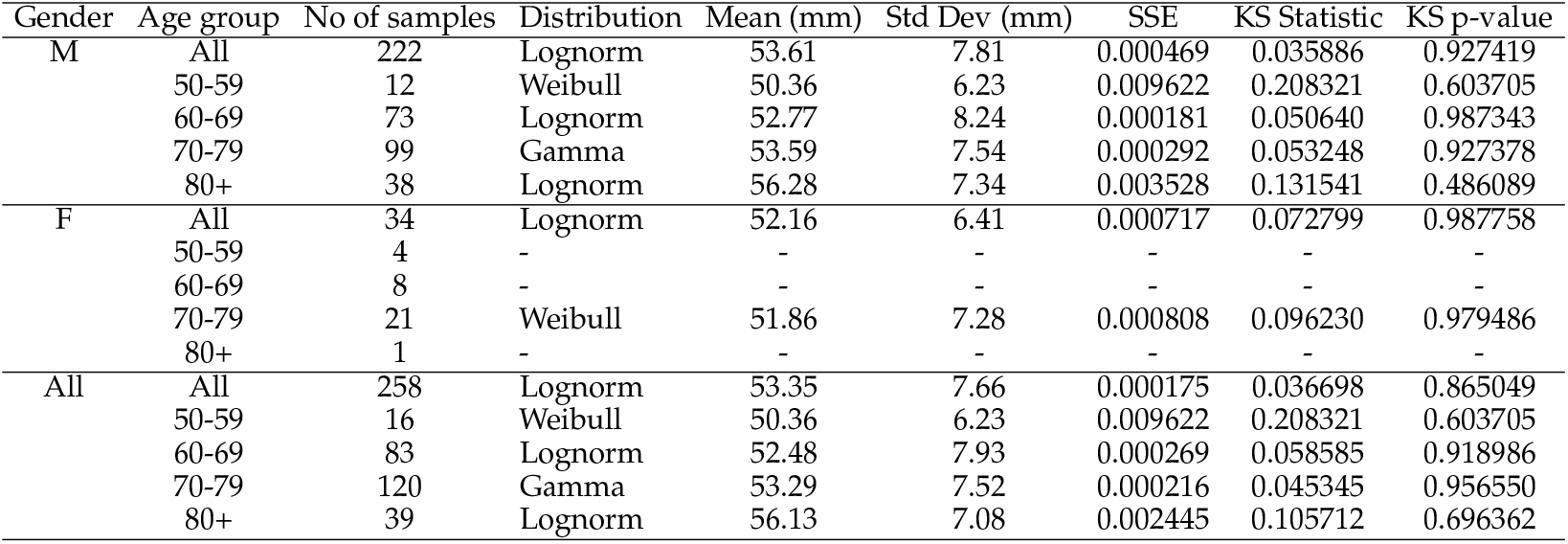
Mean, standard deviation, SSE, KS Statistic, and KS p-value for maximum aneurysm diameter across demographics.

#### (G(a)(i)) Summary Statistics

##### Algorithm 4

Spherical control point morphing

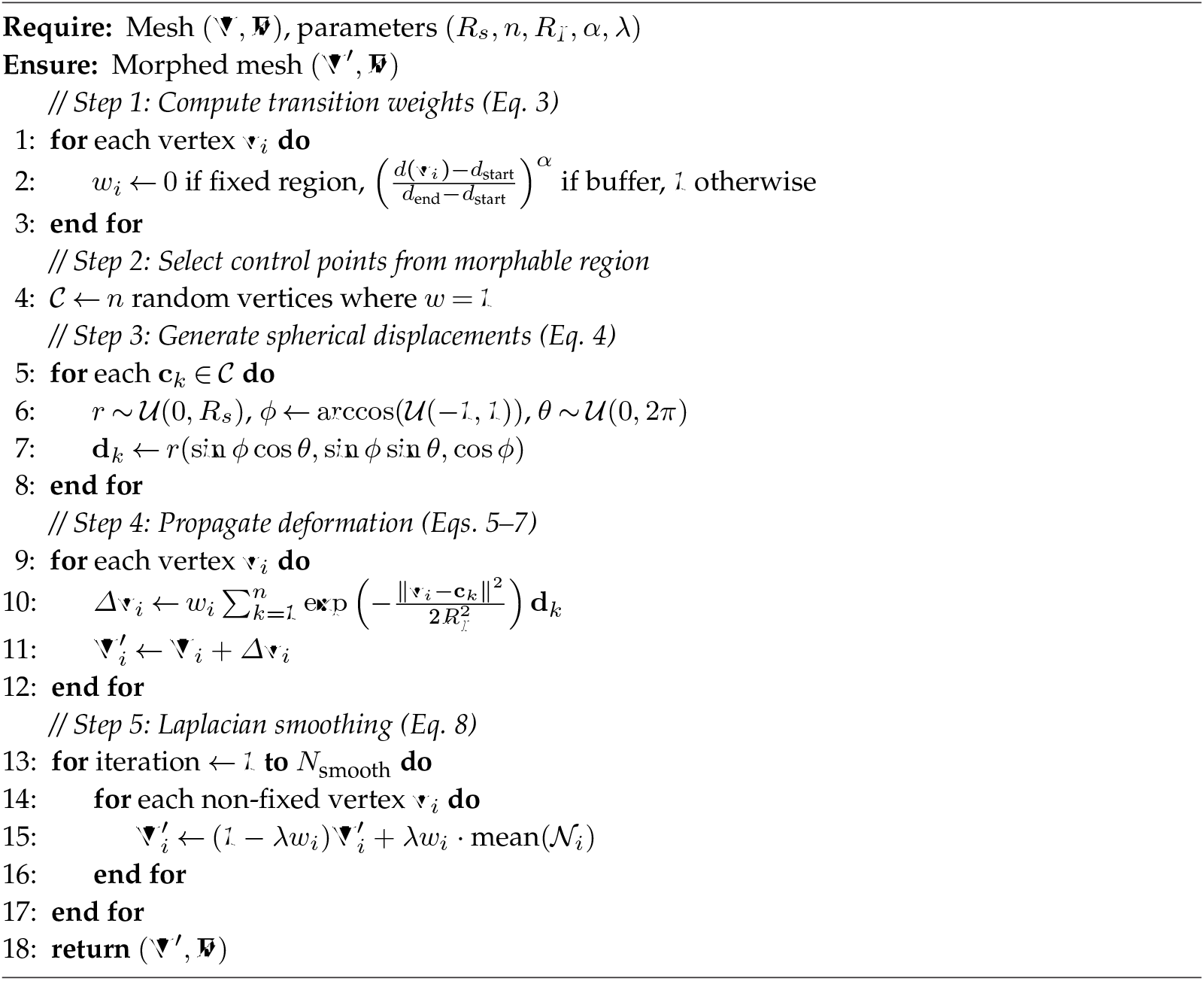

##### Algorithm 5

Population bounds validation

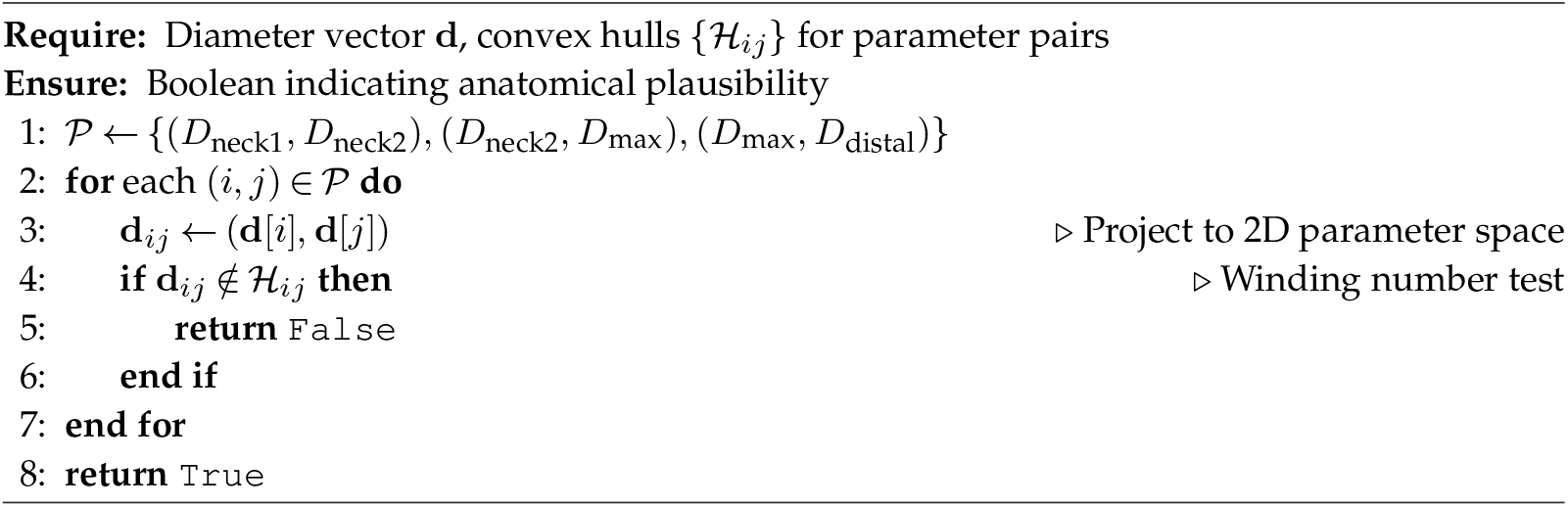

#### (G(a)(ii)) Fitted Parameters

##### Algorithm 6

Hemodynamic biomarker extraction

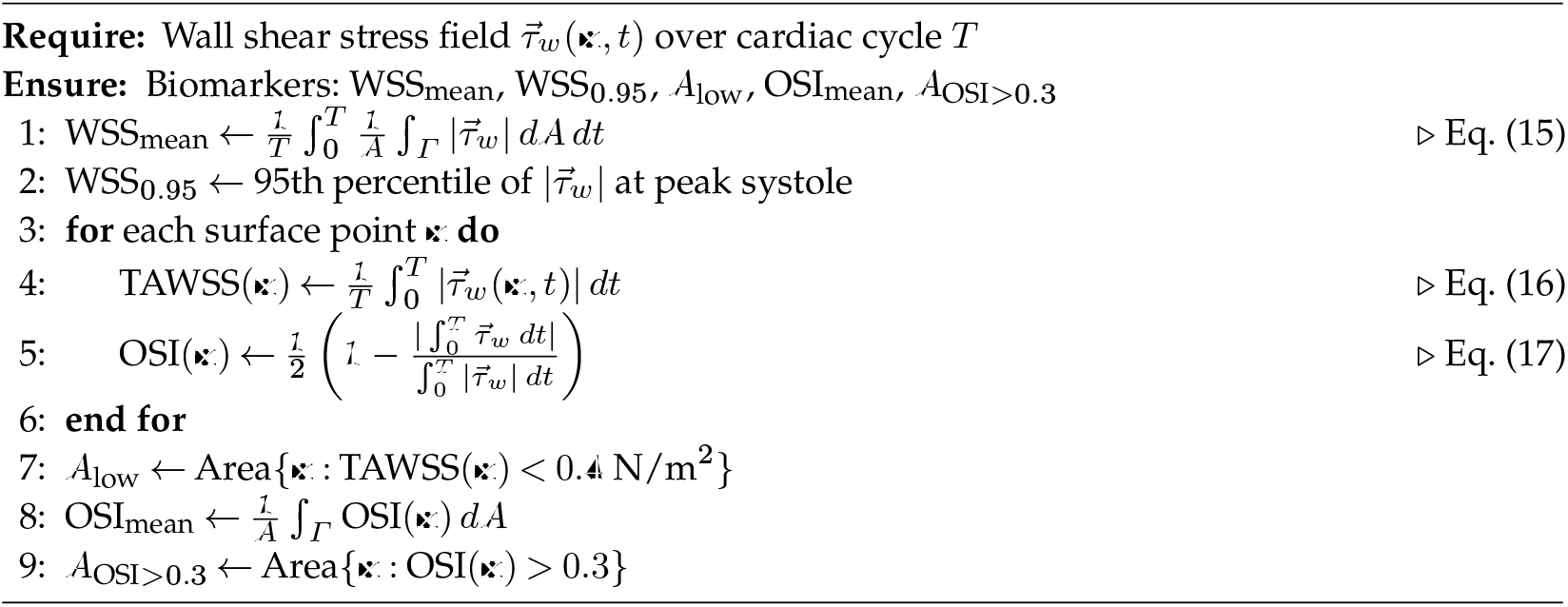

**Table 7.** Distribution parameters for maximum aneurysm diameter across demographics, including SSE, KS Statistic, and KS p-value.

(G(a)(ii)) Fitted Parameters
| Gender | Age group | No of samples | Distribution | Parameters | SSE | KS Statistic | KS p-value |
| --- | --- | --- | --- | --- | --- | --- | --- |
| M | All | 222 | Lognorm | $s = 0.374, \mu = 33.787, \sigma = 18.474$ | 0.000469 | 0.035886 | 0.927419 |
| | 50-59 | 12 | Weibull | $k = 1.164, \lambda = 8.277, \theta = 42.463$ | 0.009622 | 0.208321 | 0.603705 |
| | 60-69 | 73 | Lognorm | $s = 0.474, \mu = 36.627, \sigma = 14.410$ | 0.000181 | 0.050640 | 0.987343 |
| | 70-79 | 99 | Gamma | $\alpha = 5.477, \theta = 3.103, \mu = 36.592$ | 0.000292 | 0.053248 | 0.927378 |
| | 80+ | 38 | Lognorm | $s = 0.358, \mu = 36.674, \sigma = 18.383$ | 0.003528 | 0.131541 | 0.486089 |
| F | All | 34 | Lognorm | $s = 0.240, \mu = 25.745, \sigma = 25.670$ | 0.000717 | 0.072799 | 0.987758 |
|  | 50-59 | - | - | - | - | - | - |
|  | 60-69 | 9 | - | - | - | - | - |
| | 70-79 | 21 | Weibull | $k = 1.368, \lambda = 8.540, \theta = 11.904$ | 0.000808 | 0.096230 | 0.979486 |
|  | 80+ | 4 | - | - | - | - | - |
| All | All | 258 | Lognorm | $s = 0.364, \mu = 33.341, \sigma = 18.716$ | 0.000175 | 0.036698 | 0.865049 |
| | 50-59 | 12 | Weibull | $k = 1.164, \lambda = 8.277, \theta = 42.463$ | 0.009622 | 0.208321 | 0.603705 |
| | 60-69 | 84 | Lognorm | $s = 0.469, \mu = 36.863, \sigma = 13.973$ | 0.000269 | 0.058585 | 0.918986 |
| | 70-79 | 120 | Gamma | $\alpha = 5.421, \theta = 3.148, \mu = 36.220$ | 0.000216 | 0.045345 | 0.956550 |
| | 80+ | 42 | Lognorm | $s = 0.346, \mu = 36.668, \sigma = 18.312$ | 0.002445 | 0.105712 | 0.696362 |

### G(b) Neck Diameter 1

#### (G(b)(i)) Summary Statistics

**Table 8.** Mean, standard deviation, SSE, KS Statistic, and KS p-value for neck diameter 1 across demographics.

| Gender | Age group | No of samples | Distribution | Mean (mm) | Std Dev (mm) | SSE | KS Statistic | KS p-value |
| --- | --- | --- | --- | --- | --- | --- | --- | --- |
| M | All | 222 | Weibull | 22.78 | 2.93 | 0.002806 | 0.041973 | 0.813131 |
|  | 50-59 | 12 | Weibull | 21.63 | 1.76 | 0.013337 | 0.148904 | 0.917750 |
|  | 60-69 | 73 | Gamma | 22.93 | 3.15 | 0.001281 | 0.075181 | 0.775390 |
|  | 70-79 | 99 | Normal | 22.46 | 2.53 | 0.001254 | 0.059292 | 0.856699 |
|  | 80+ | 38 | Weibull | 23.71 | 3.43 | 0.001655 | 0.093975 | 0.859387 |
| F | All | 34 | Lognorm | 21.01 | 3.16 | 0.007559 | 0.083950 | 0.953903 |
|  | 50-59 | - | - | - | - | - | - | - |
|  | 60-69 | - | - | - | - | - | - | - |
|  | 70-79 | 21 | Lognorm | 20.90 | 2.72 | 0.008791 | 0.110572 | 0.934843 |
|  | 80+ | - | - | - | - | - | - | - |
| All | All | 258 | Weibull | 22.57 | 3.03 | 0.004977 | 0.048758 | 0.555093 |
|  | 50-59 | 12 | Weibull | 21.63 | 1.76 | 0.013337 | 0.148904 | 0.917750 |
|  | 60-69 | 84 | Weibull | 22.70 | 3.32 | 0.002349 | 0.061693 | 0.886779 |
|  | 70-79 | 120 | Weibull | 22.19 | 2.63 | 0.006303 | 0.050052 | 0.909638 |
|  | 80+ | 42 | Weibull | 23.67 | 3.44 | 0.001667 | 0.082747 | 0.913150 |

**Table 9.** Distribution parameters for neck diameter 1 across demographics, including SSE, KS Statistic, and KS p-value.

| Gender | Age group | No of samples | Distribution | Parameters | SSE | KS Statistic | KS p-value |
| --- | --- | --- | --- | --- | --- | --- | --- |
| M | All | 222 | Weibull | $k = 2.195, \lambda = 6.898, \theta = 16.671$ | 0.002806 | 0.041973 | 0.813131 |
| | 50-59 | 12 | Weibull | $k = 2.565, \lambda = 4.708, \theta = 17.445$ | 0.013337 | 0.148904 | 0.917750 |
| | 60-69 | 73 | Gamma | $\alpha = 8.343, \theta = 1.106, \mu = 13.706$ | 0.001281 | 0.075181 | 0.775390 |
| | 70-79 | 99 | Normal | $\mu = 22.460, \sigma = 2.532$ | 0.001254 | 0.059292 | 0.856699 |
| | 80+ | 38 | Weibull | $k = 2.148, \lambda = 7.903, \theta = 16.704$ | 0.001655 | 0.093975 | 0.859387 |
| F | All | 34 | Lognorm | $s = 0.368, \mu = 12.450, \sigma = 8.011$ | 0.007559 | 0.083950 | 0.953903 |
|  | 50-59 | - | - | - | - | - | - |
|  | 60-69 | 9 | - | - | - | - | - |
| | 70-79 | 21 | Lognorm | $s = 0.351, \mu = 13.228, \sigma = 7.217$ | 0.008791 | 0.110572 | 0.934843 |
|  | 80+ | 4 | - | - | - | - | - |
| All | All | 258 | Weibull | $k = 2.419, \lambda = 7.790, \theta = 15.662$ | 0.004977 | 0.048758 | 0.555093 |
| | 50-59 | 12 | Weibull | $k = 2.565, \lambda = 4.708, \theta = 17.445$ | 0.013337 | 0.148904 | 0.917750 |
| | 60-69 | 84 | Weibull | $k = 2.402, \lambda = 8.434, \theta = 15.221$ | 0.002349 | 0.061693 | 0.886779 |
| | 70-79 | 120 | Weibull | $k = 2.520, \lambda = 6.997, \theta = 15.976$ | 0.006303 | 0.050052 | 0.909638 |
| | 80+ | 42 | Weibull | $k = 2.088, \lambda = 7.717, \theta = 16.837$ | 0.001667 | 0.082747 | 0.913150 |

#### (G(b)(ii)) Fitted Parameters

### G(c) Neck Diameter 2

#### (G(c)(i)) Summary Statistics

**Table 10.** Mean, standard deviation, SSE, KS Statistic, and KS p-value for neck diameter 2 across demographics.

| Gender | Age group | No of samples | Distribution | Mean (mm) | Std Dev (mm) | SSE | KS Statistic | KS p-value |
| --- | --- | --- | --- | --- | --- | --- | --- | --- |
| M | All | 222 | Lognorm | 24.76 | 3.33 | 0.003620 | 0.052950 | 0.544399 |
|  | 50-59 | 12 | Gamma | 23.89 | 2.02 | 0.004664 | 0.103162 | 0.997874 |
|  | 60-69 | 73 | Gamma | 25.19 | 3.86 | 0.001214 | 0.071028 | 0.829508 |
|  | 70-79 | 99 | Weibull | 24.33 | 2.71 | 0.002732 | 0.057226 | 0.883420 |
|  | 80+ | 38 | Lognorm | 25.33 | 3.79 | 0.003080 | 0.097301 | 0.830580 |
| F | All | 34 | Lognorm | 22.42 | 3.18 | 0.013024 | 0.101076 | 0.843759 |
|  | 50-59 | - | - | - | - | - | - | - |
|  | 60-69 | - | - | - | - | - | - | - |
|  | 70-79 | 21 | Gamma | 22.36 | 3.25 | 0.008291 | 0.161290 | 0.589799 |
|  | 80+ | - | - | - | - | - | - | - |
| All | All | 258 | Lognorm | 24.48 | 3.42 | 0.002856 | 0.054696 | 0.408440 |
|  | 50-59 | 12 | Gamma | 23.89 | 2.02 | 0.004664 | 0.103162 | 0.997874 |
|  | 60-69 | 84 | Gamma | 24.88 | 3.95 | 0.001136 | 0.066796 | 0.823384 |
|  | 70-79 | 120 | Weibull | 23.99 | 2.91 | 0.003287 | 0.056820 | 0.812062 |
|  | 80+ | 42 | Lognorm | 25.22 | 3.66 | 0.001197 | 0.071120 | 0.973537 |

#### (G(c)(ii)) Fitted Parameters

**Table 11.** Distribution parameters for neck diameter 2 across demographics, including SSE, KS Statistic, and KS p-value.

| Gender | Age group | No of samples | Distribution | Parameters | SSE | KS Statistic | KS p-value |
| --- | --- | --- | --- | --- | --- | --- | --- |
| M | All | 222 | Lognorm | $s = 0.194, \mu = 7.782, \sigma = 16.663$ | 0.003620 | 0.052950 | 0.544399 |
| | 50-59 | 12 | Gamma | $\alpha = 1989.588, \theta = 0.045, \mu = -66.513$ | 0.004664 | 0.103162 | 0.997874 |
| | 60-69 | 73 | Gamma | $\alpha = 28.085, \theta = 0.729, \mu = 4.725$ | 0.001214 | 0.071028 | 0.829508 |
| | 70-79 | 99 | Weibull | $k = 2.128, \lambda = 6.223, \theta = 18.816$ | 0.002732 | 0.057226 | 0.883420 |
| | 80+ | 38 | Lognorm | $s = 0.321, \mu = 13.608, \sigma = 11.133$ | 0.003080 | 0.097301 | 0.830580 |
| F | All | 34 | Lognorm | $s = 0.236, \mu = 9.066, \sigma = 12.992$ | 0.013024 | 0.101076 | 0.843759 |
|  | 50-59 | - | - | - | - | - | - |
|  | 60-69 | 9 | - | - | - | - | - |
| | 70-79 | 21 | Gamma | $\alpha = 8.293, \theta = 1.120, \mu = 13.067$ | 0.008291 | 0.161290 | 0.589799 |
|  | 80+ | 4 | - | - | - | - | - |
| All | All | 258 | Lognorm | $s = 0.181, \mu = 5.815, \sigma = 18.356$ | 0.002856 | 0.054696 | 0.408440 |
| | 50-59 | 12 | Gamma | $\alpha = 1989.588, \theta = 0.045, \mu = -66.513$ | 0.004664 | 0.103162 | 0.997874 |
| | 60-69 | 84 | Gamma | $\alpha = 20.200, \theta = 0.883, \mu = 7.043$ | 0.001136 | 0.066796 | 0.823384 |
| | 70-79 | 120 | Weibull | $k = 2.311, \lambda = 7.001, \theta = 17.837$ | 0.003287 | 0.056820 | 0.812062 |
| | 80+ | 42 | Lognorm | $s = 0.320, \mu = 13.910, \sigma = 10.750$ | 0.001197 | 0.071120 | 0.973537 |

### G(d) Distal Diameter

#### (G(d)(i)) Summary Statistics

**Table 12.** Mean, standard deviation, SSE, KS Statistic, and KS p-value for distal diameter across demographics.

| Gender | Age group | No of samples | Distribution | Mean (mm) | Std Dev (mm) | SSE | KS Statistic | KS p-value |
| --- | --- | --- | --- | --- | --- | --- | --- | --- |
| M | All | 222 | Lognorm | 20.09 | 6.12 | 0.002399 | 0.091277 | 0.046414 |
|  | 50-59 | 12 | Weibull | 19.90 | 5.86 | 0.007914 | 0.166667 | 0.839863 |
|  | 60-69 | 73 | Lognorm | 19.71 | 6.15 | 0.001681 | 0.096837 | 0.471175 |
|  | 70-79 | 99 | Lognorm | 20.02 | 5.96 | 0.001173 | 0.087687 | 0.408394 |
|  | 80+ | 38 | Weibull | 21.04 | 6.45 | 0.000914 | 0.083978 | 0.930959 |
| F | All | 34 | Gamma | 17.74 | 4.46 | 0.001505 | 0.072225 | 0.988764 |
|  | 50-59 | - | - | - | - | - | - | - |
|  | 60-69 | - | - | - | - | - | - | - |
|  | 70-79 | 21 | Lognorm | 18.02 | 5.16 | 0.002393 | 0.079638 | 0.997780 |
|  | 80+ | - | - | - | - | - | - | - |
| All | All | 258 | Lognorm | 19.79 | 5.96 | 0.000935 | 0.083522 | 0.051574 |
|  | 50-59 | 12 | Weibull | 19.90 | 5.86 | 0.007914 | 0.166667 | 0.839863 |
|  | 60-69 | 84 | Lognorm | 19.44 | 5.90 | 0.000821 | 0.074251 | 0.715024 |
|  | 70-79 | 120 | Weibull | 19.67 | 5.88 | 0.000692 | 0.085512 | 0.325394 |
|  | 80+ | 42 | Gamma | 20.79 | 6.23 | 0.000122 | 0.083510 | 0.907752 |

#### (G(d)(ii)) Fitted Parameters

**Table 13.**
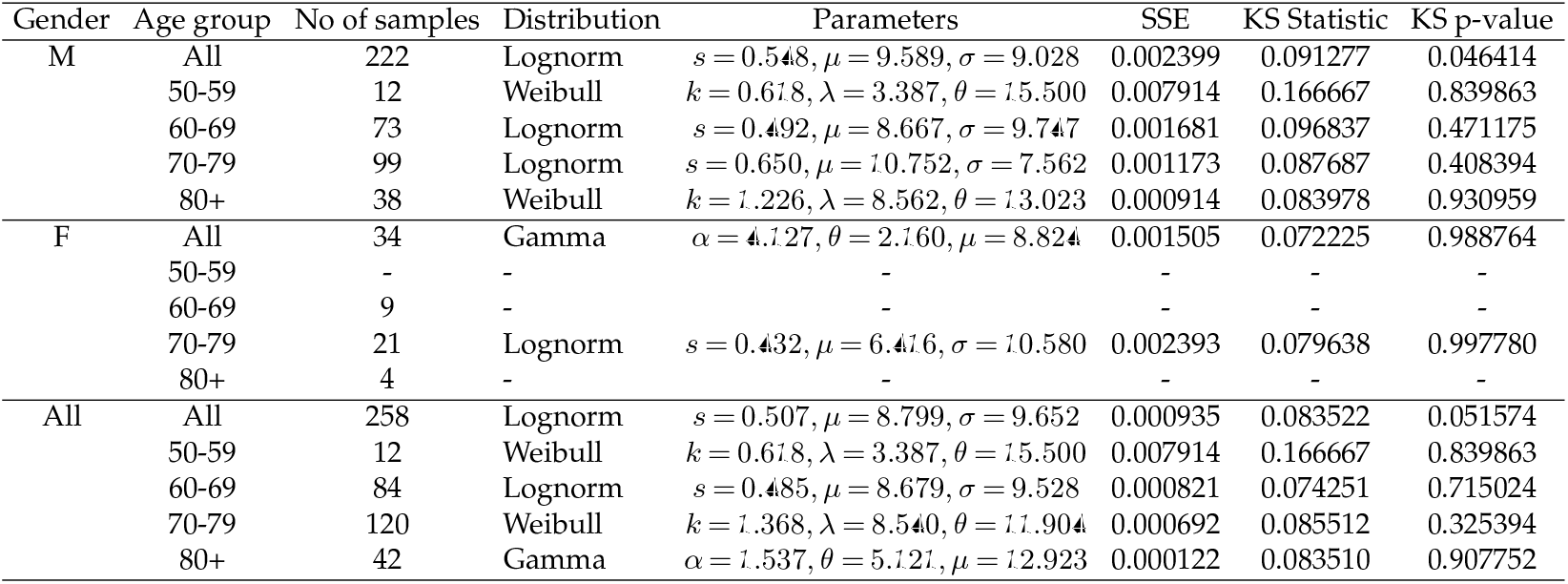
Distribution parameters for distal diameter across demographics, including SSE, KS Statistic, and KS p-value.

## H. Distribution Fitting Visualisations

**Figure 14.**
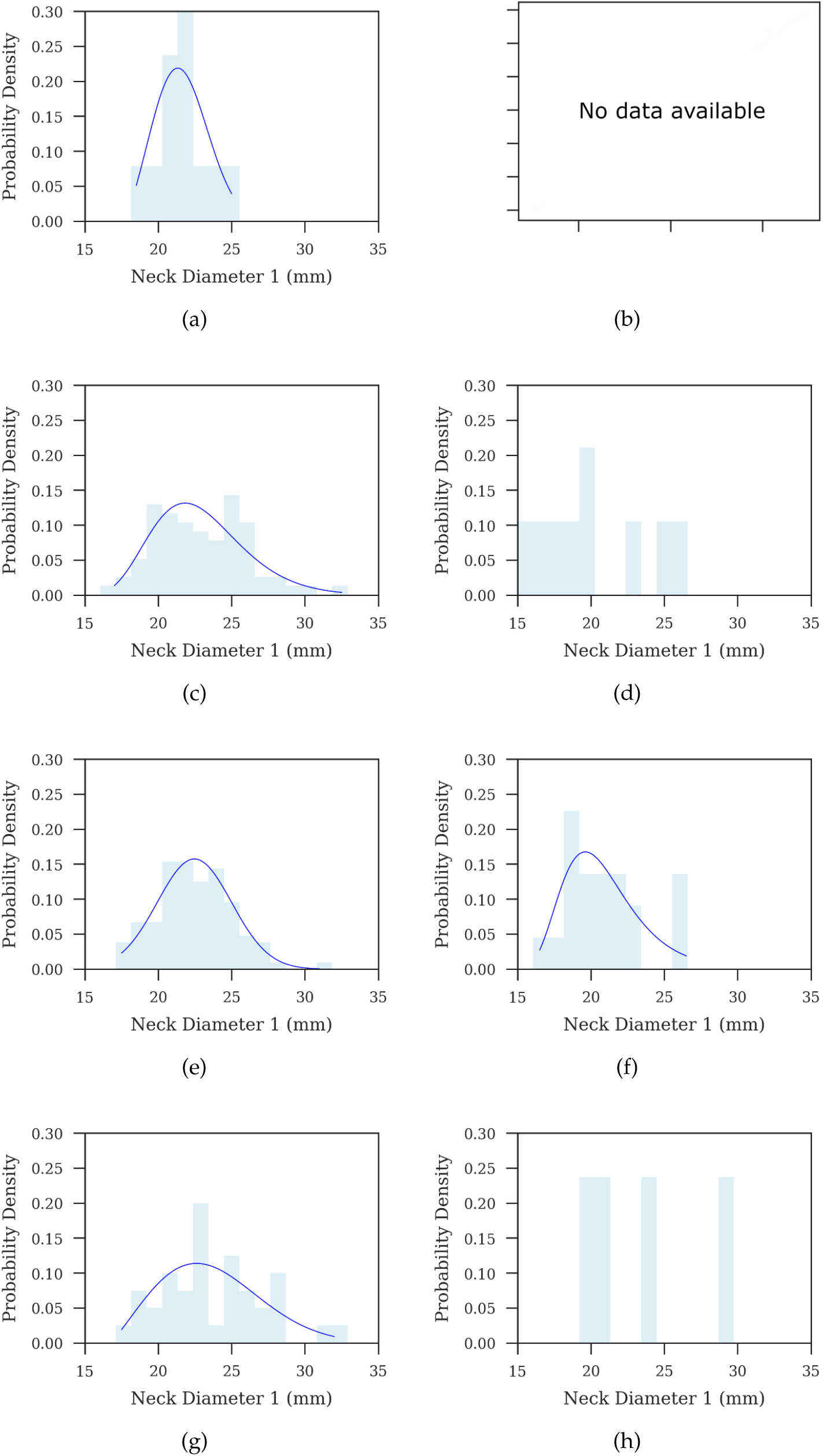
Proximal neck diameter (Neck Diameter 1) distribution grouped by age and gender. Each row corresponds to an age group: 50-59, 60-69, 70-79, and 80+. Each column represents gender, with the left column for males and the right column for females.

**Figure 15.**
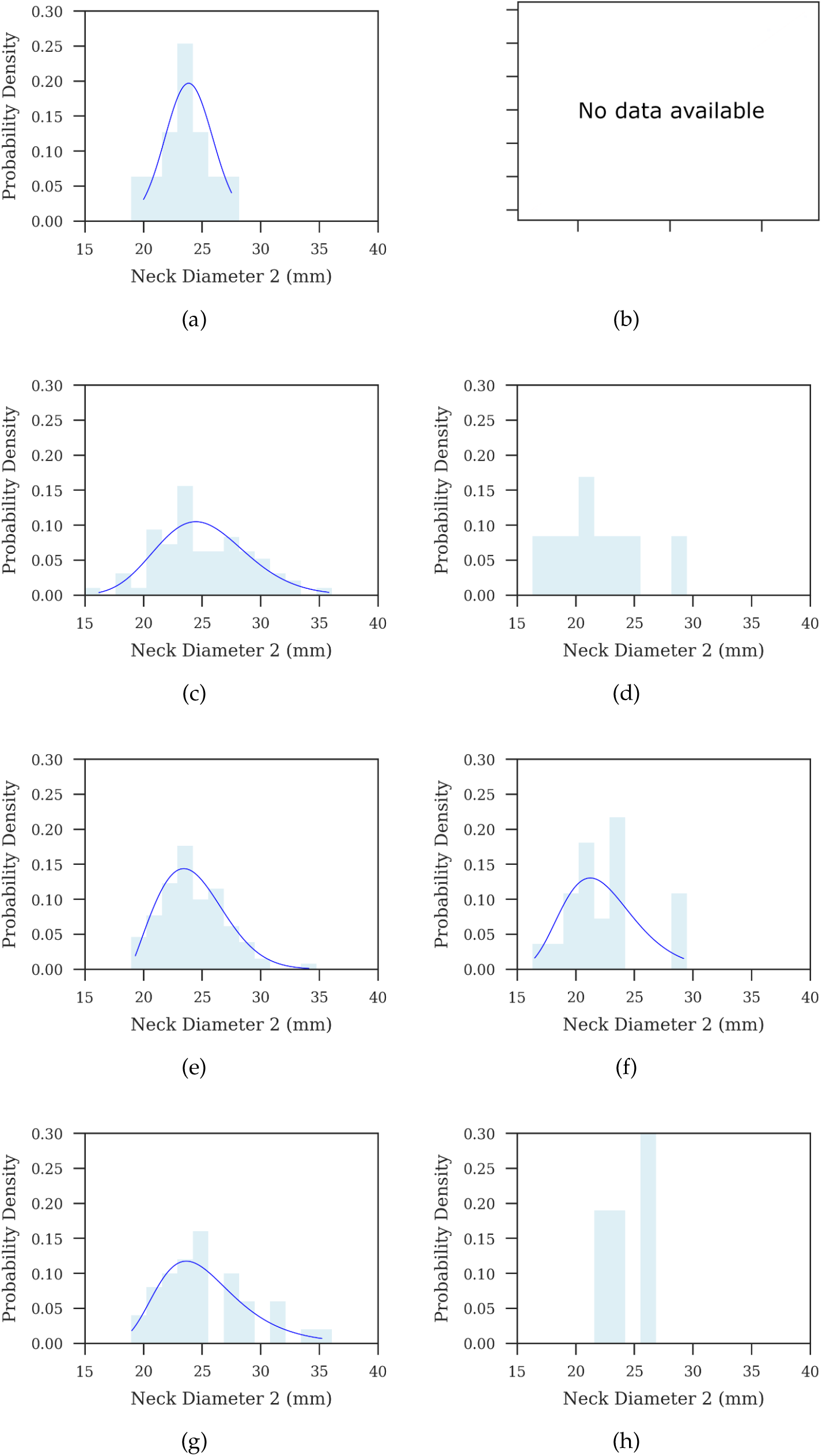
Distal neck diameter (Neck Diameter 2) distribution grouped by age and gender. Each row corresponds to an age group: 50-59, 60-69, 70-79, and 80+. Each column represents gender, with the left column for males and the right column for females.

**Figure 16.**
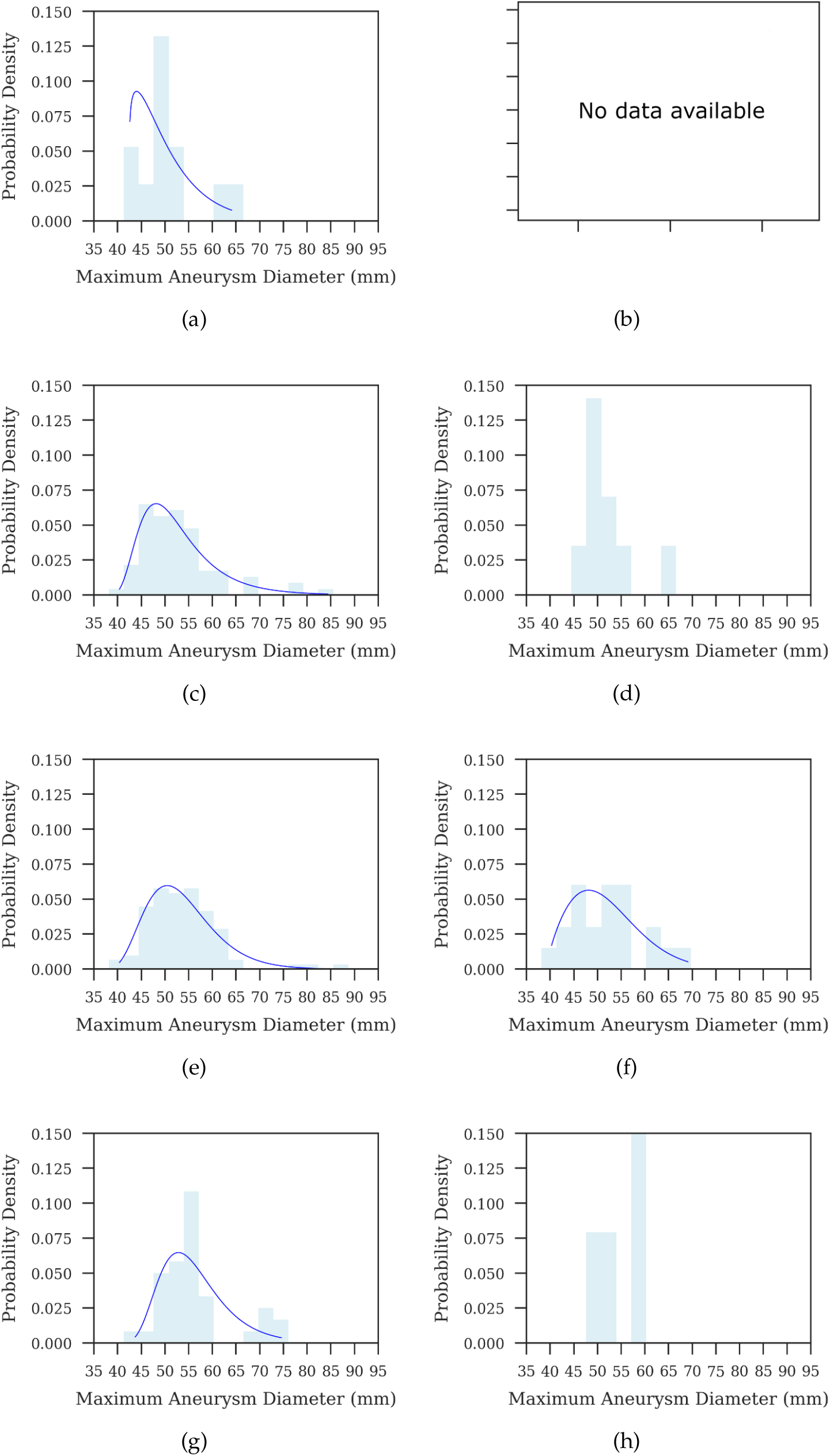
Maximum aneurysm diameter distribution grouped by age and gender. Each row corresponds to an age group: 50-59, 60-69, 70-79, and 80+. Each column represents gender, with the left column for males and the right column for females.

**Figure 17.**
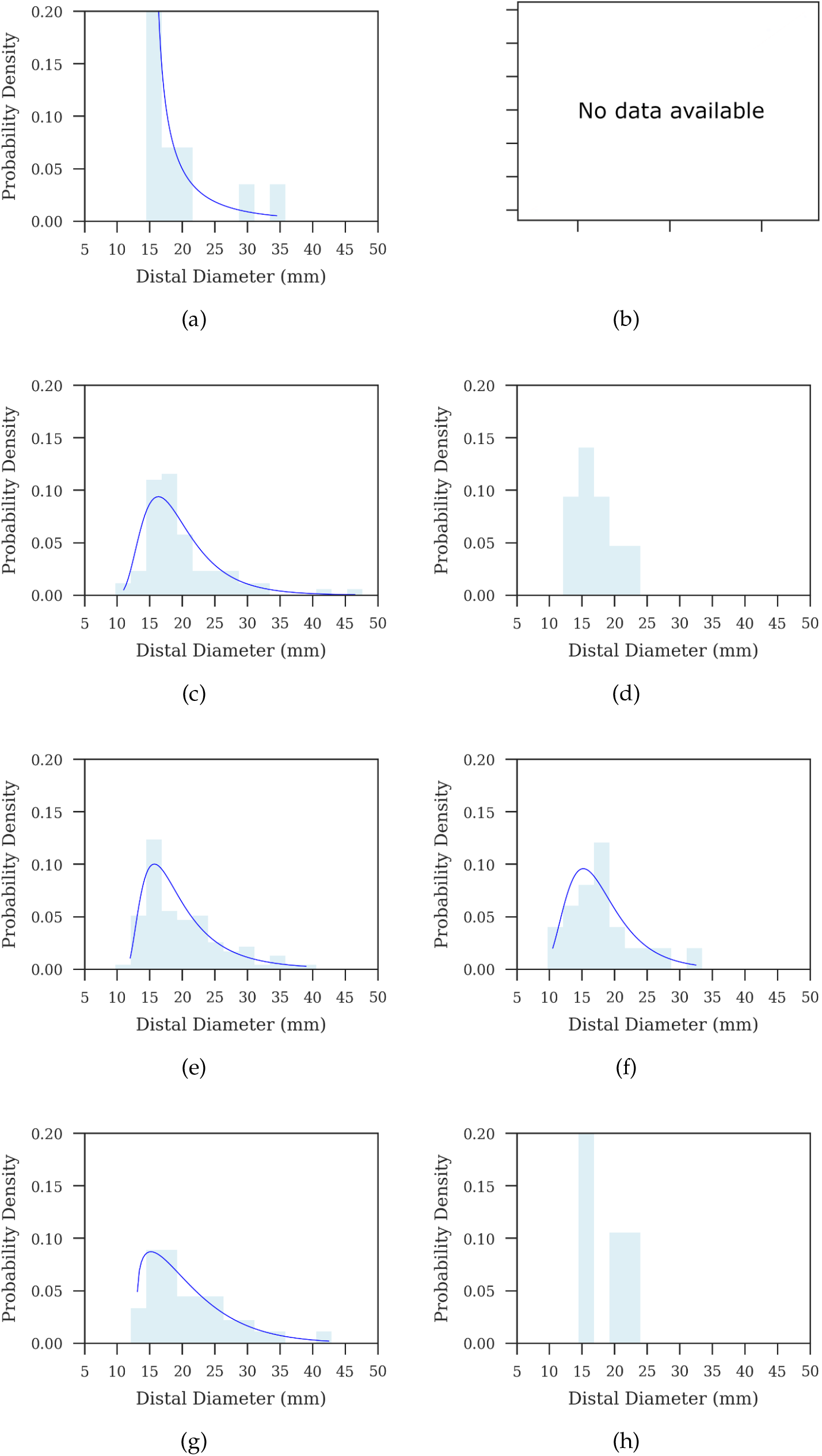
Distal diameter distribution grouped by age and gender. Each row corresponds to an age group: 50-59, 60-69, 70-79, and 80+. Each column represents gender, with the left column for males and the right column for females.

**Figure 18.**
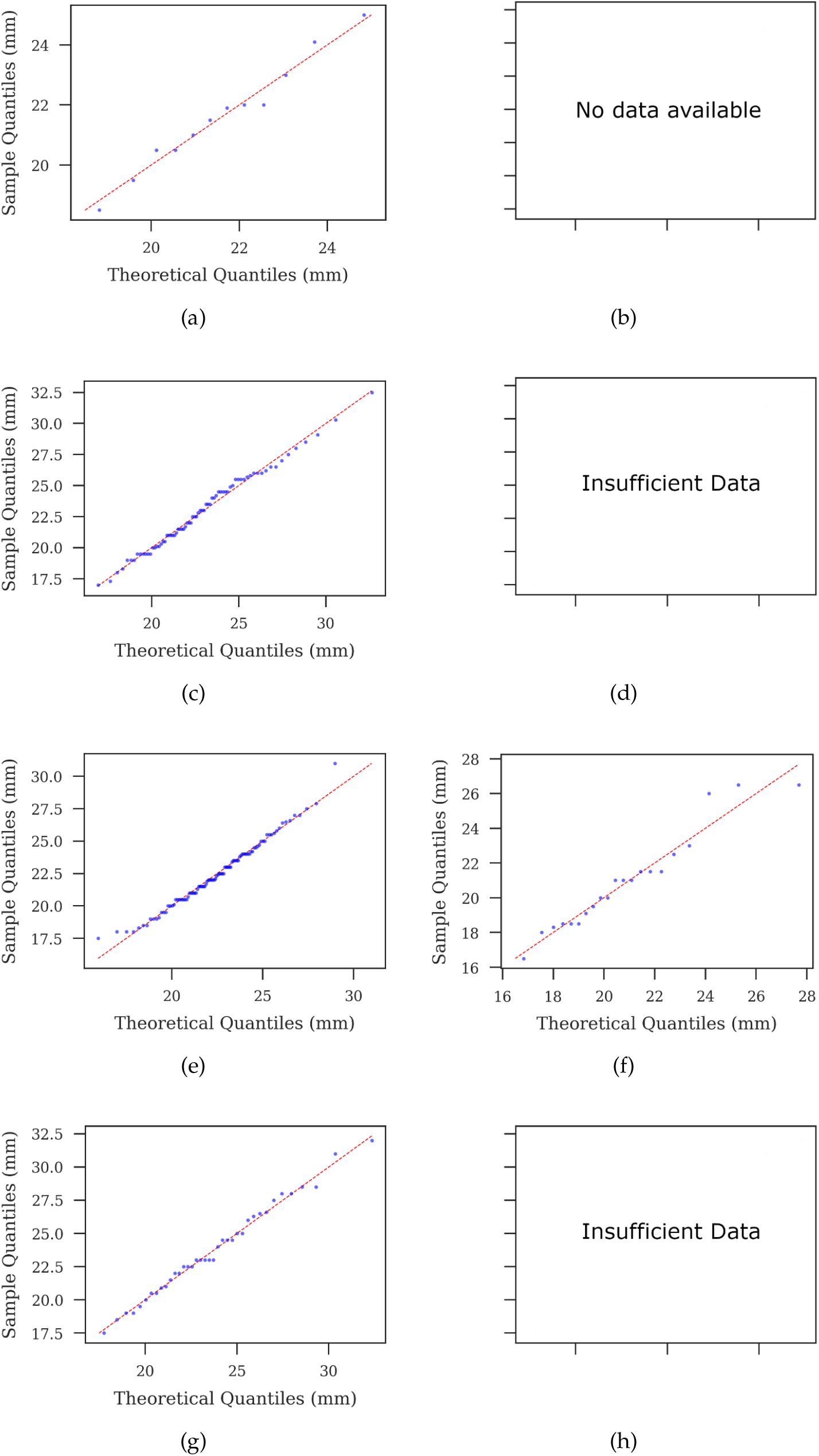
Q-Q plots for proximal neck diameter (Neck Diameter 1) grouped by age and gender. Each row corresponds to an age group: 50-59, 60-69, 70-79, and 80+. Each column represents gender, with the left column for males and the right column for females.

**Figure 19.**
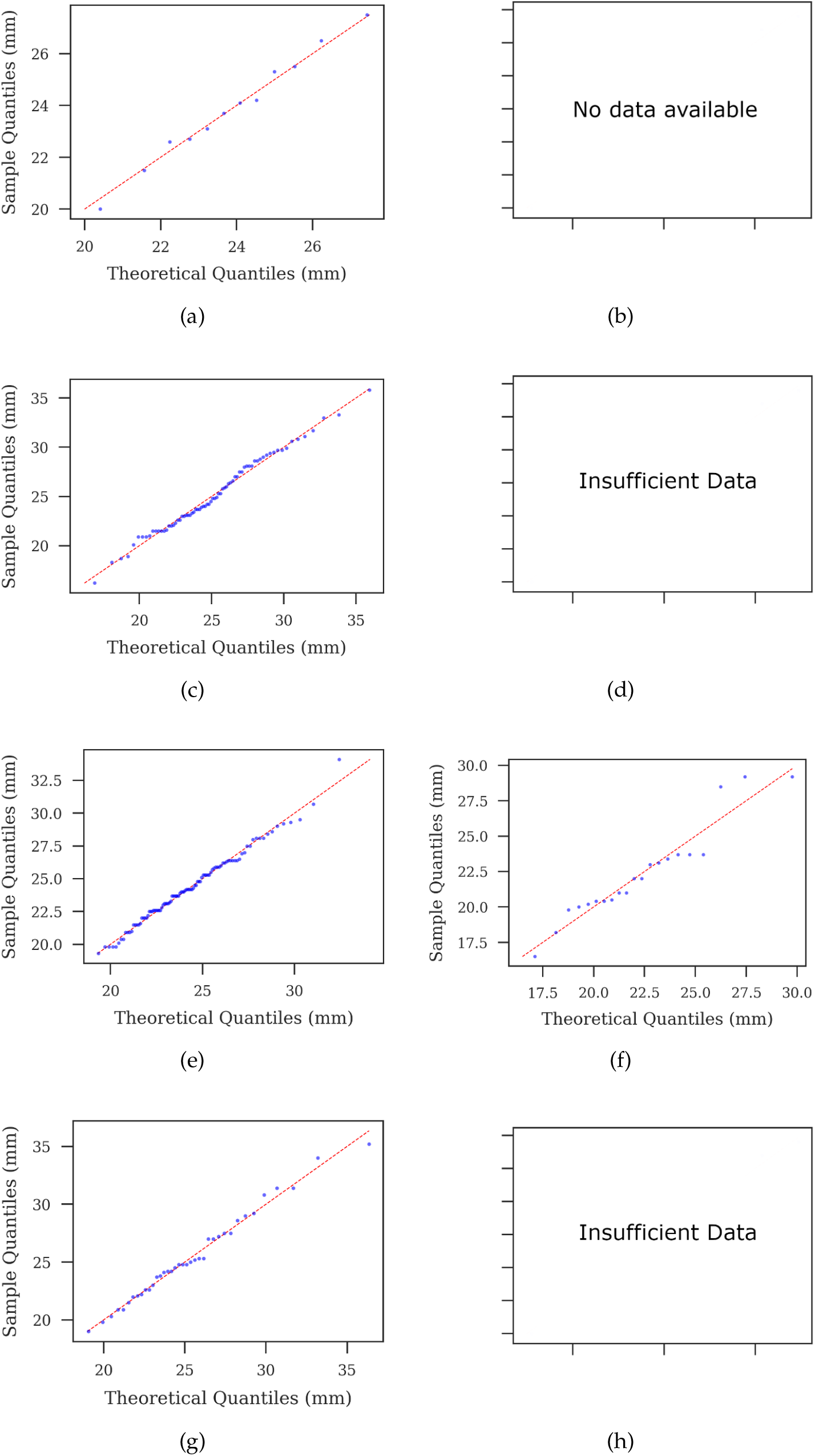
Q-Q plots for distal neck diameter (Neck Diameter 2) grouped by age and gender. Each row corresponds to an age group: 50-59, 60-69, 70-79, and 80+. Each column represents gender, with the left column for males and the right column for females.

**Figure 20.**
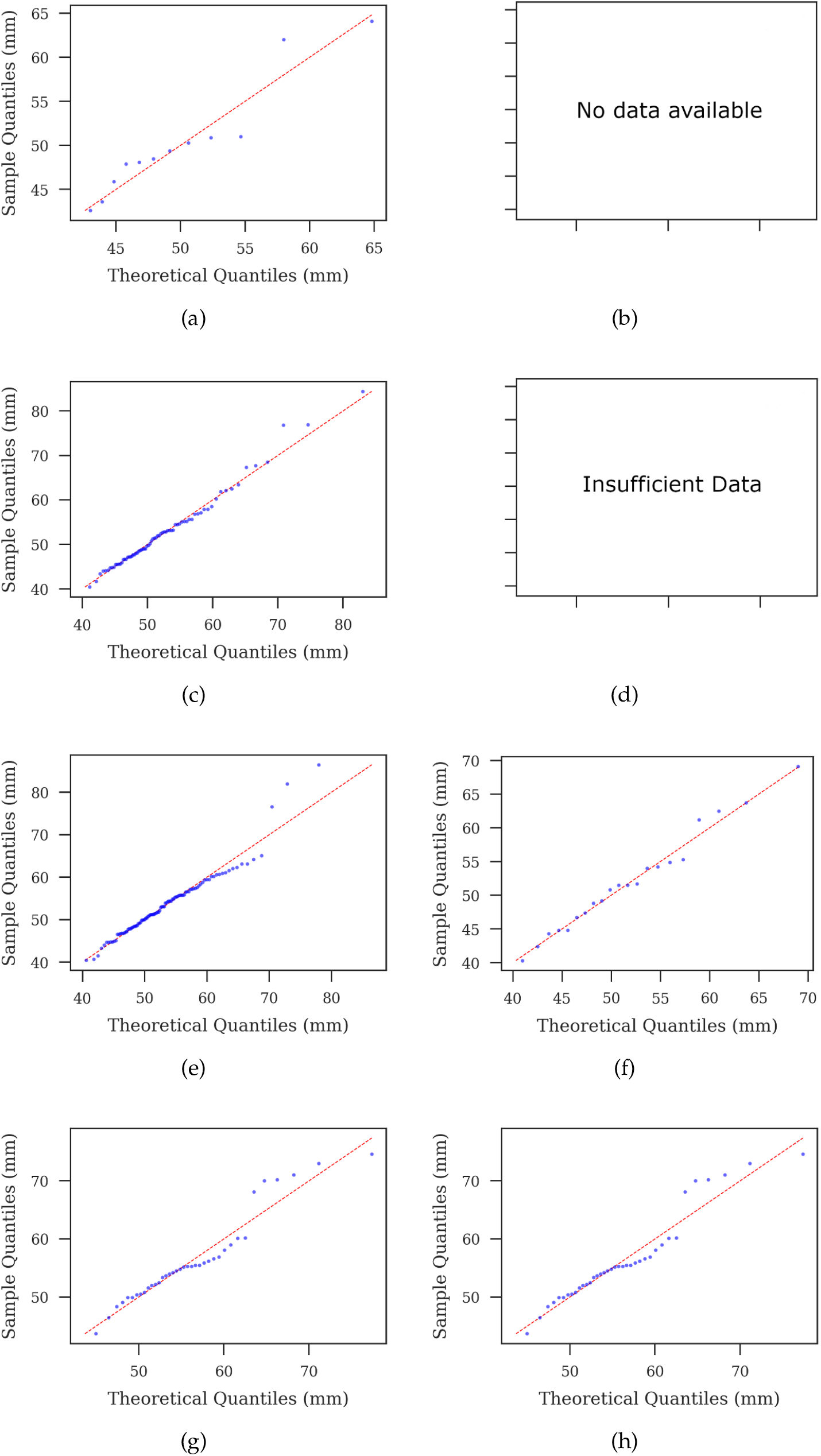
Q-Q plots for maximum aneurysm diameter grouped by age and gender. Each row corresponds to an age group: 50-59, 60-69, 70-79, and 80+. Each column represents gender, with the left column for males and the right column for females.

**Figure 21.**
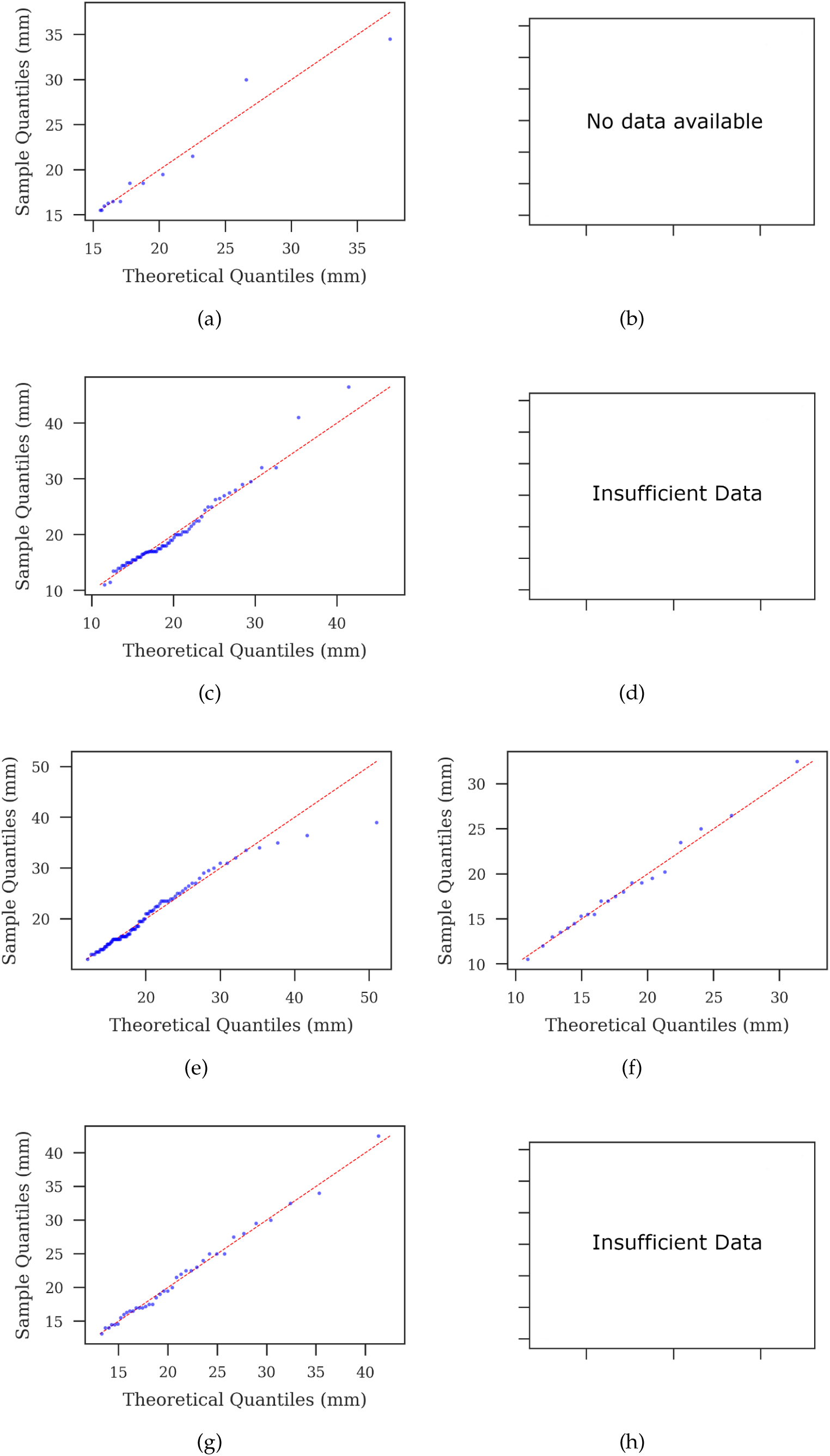
Q-Q plots for distal diameter grouped by age and gender. Each row corresponds to an age group: 50-59, 60-69, 70-79, and 80+. Each column represents gender, with the left column for males and the right column for females.

## I. Demographic Parameter Relationships

**Figure 22.**
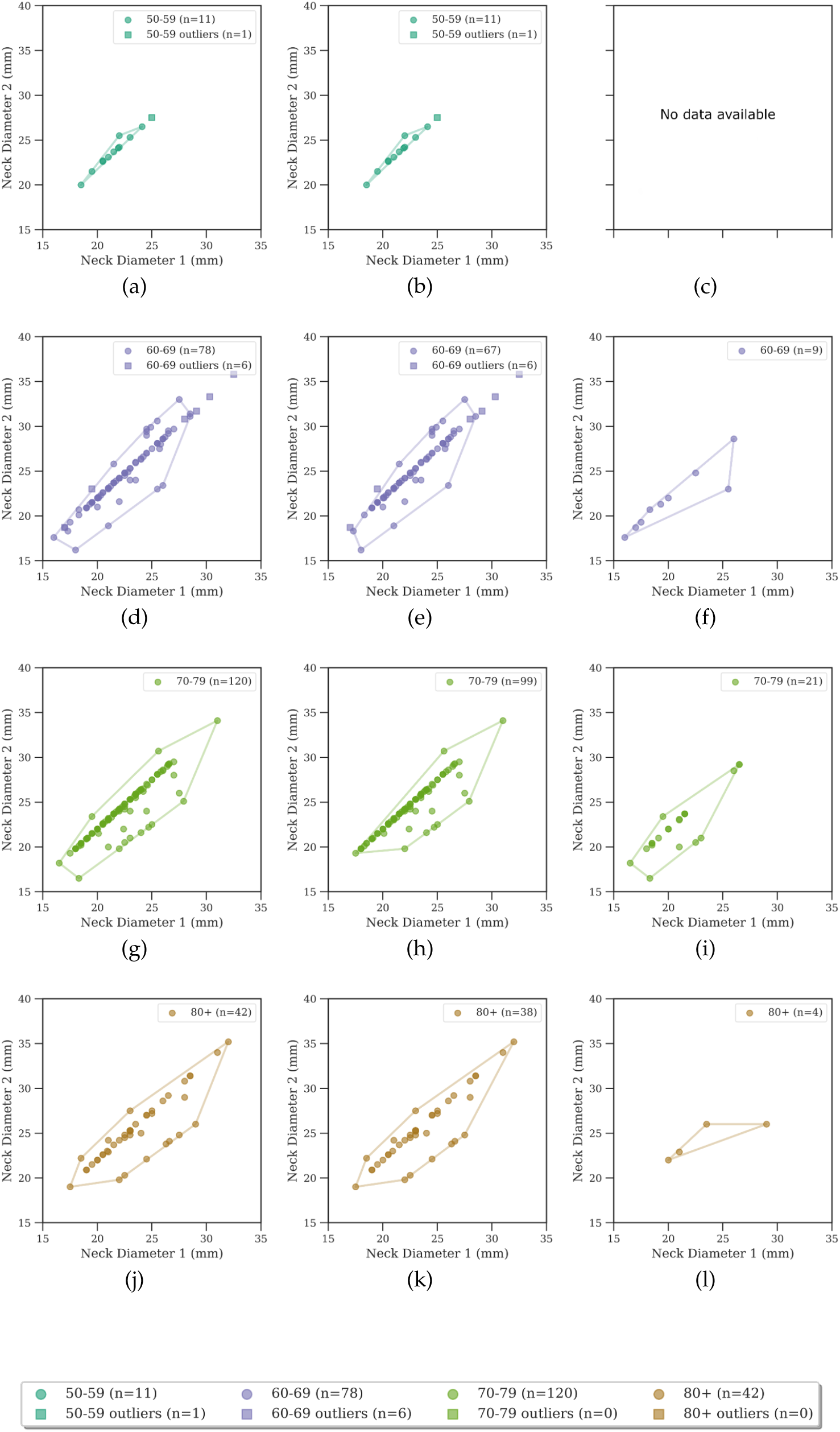
Neck diameter relationships across age groups and genders

**Figure 23.**
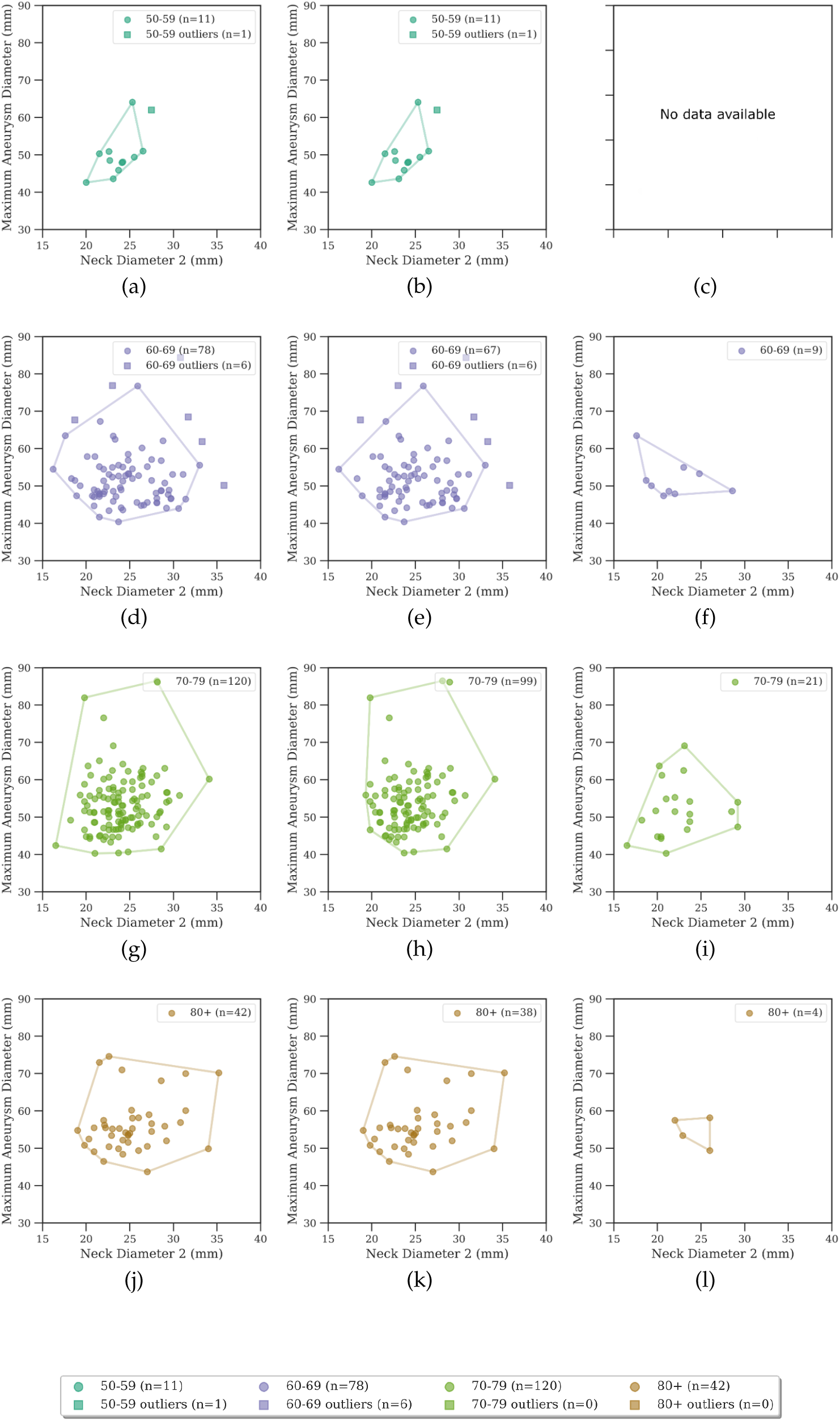
Neck diameter 2 vs maximum aneurysm diameter relationships across age groups and genders

**Figure 24.**
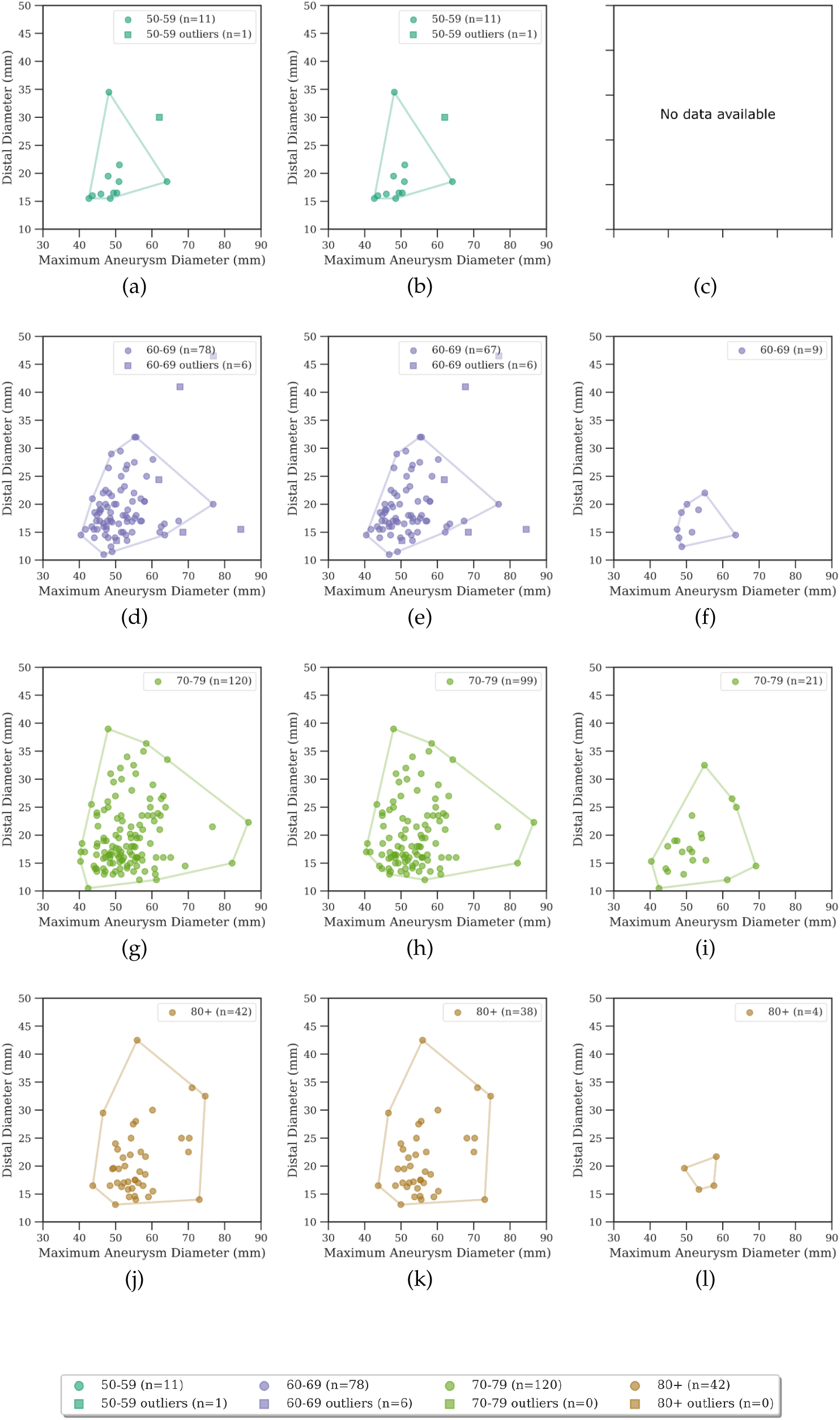
Maximum aneurysm diameter vs distal diameter relationships across age groups and genders

## J. Morphing Effectiveness

### J(a) KL Divergence Comparisons

**Table 14.** KL divergence values comparing morphed variants and patient data across key parameters and demographic groups.

| Parameter | Gender | Age Group | KL Divergence |
| --- | --- | --- | --- |
| Neck 1 Diameter | Female | 70–79 | 0.9224 |
|  | Male | 60–69 | 0.8218 |
|  | Male | 70–79 | 1.0421 |
|  | Male | 80+ | 0.7670 |
| Neck 2 Diameter | Female | 70–79 | 0.1784 |
|  | Male | 60–69 | 0.1135 |
|  | Male | 70–79 | 0.6666 |
|  | Male | 80+ | 0.2357 |
| Maximum Diameter | Female | 70–79 | 0.2632 |
|  | Male | 60–69 | 0.0180 |
|  | Male | 70–79 | 0.0121 |
|  | Male | 80+ | 0.1532 |

**Figure 25.**
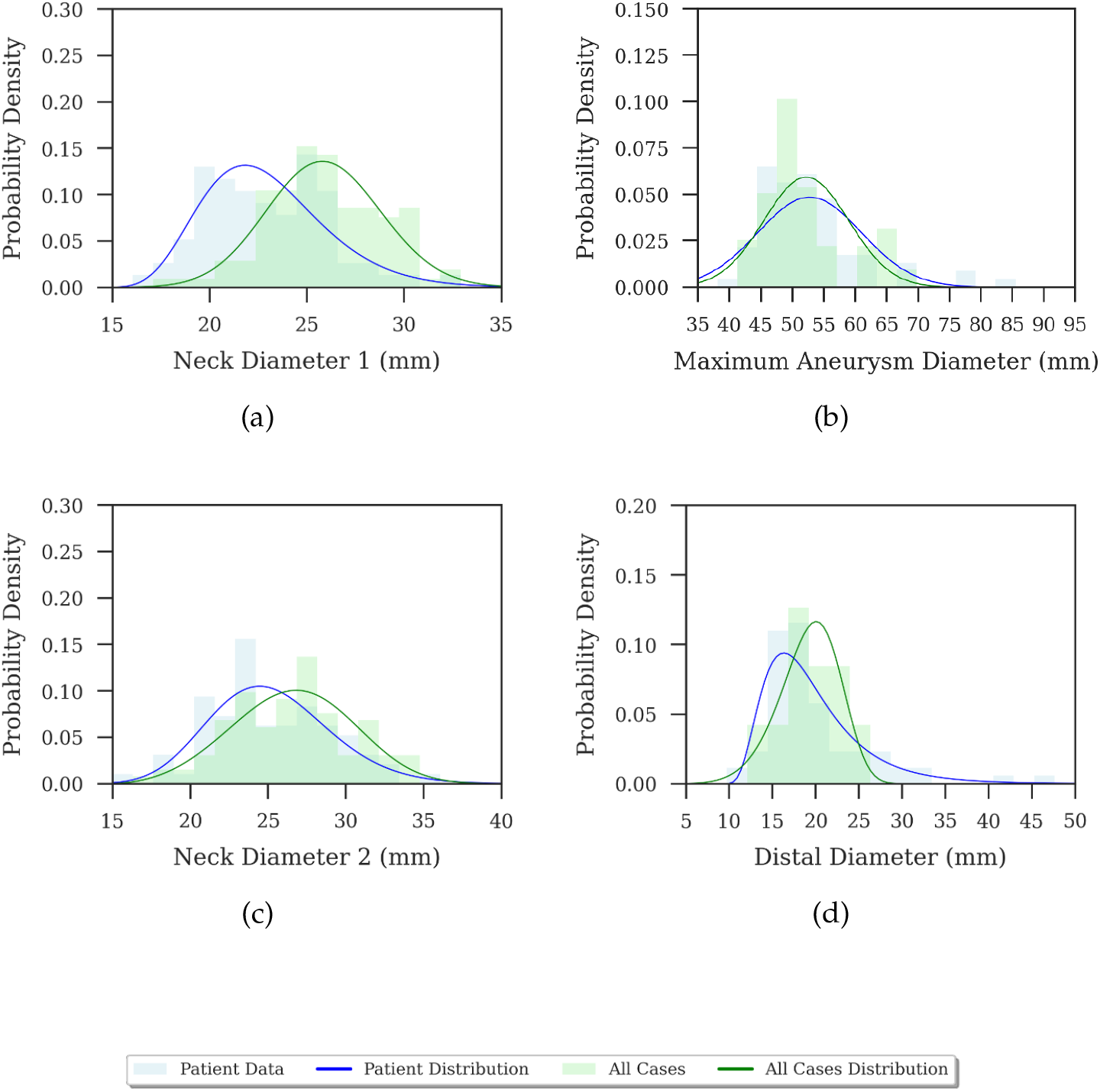
Comparison of patient data vs. generated statistical variants for the male 60-69 age group. (a) Neck 1 diameter comparison. (b) Maximum aneurysm diameter comparison. (c) Neck 2 diameter comparison. (d) Distal diameter comparison. Each subfigure shows the probability density function of patient data (blue) alongside the morphed statistical variants (green). The third row provides an enlarged legend for the plots.

**Figure 26.**
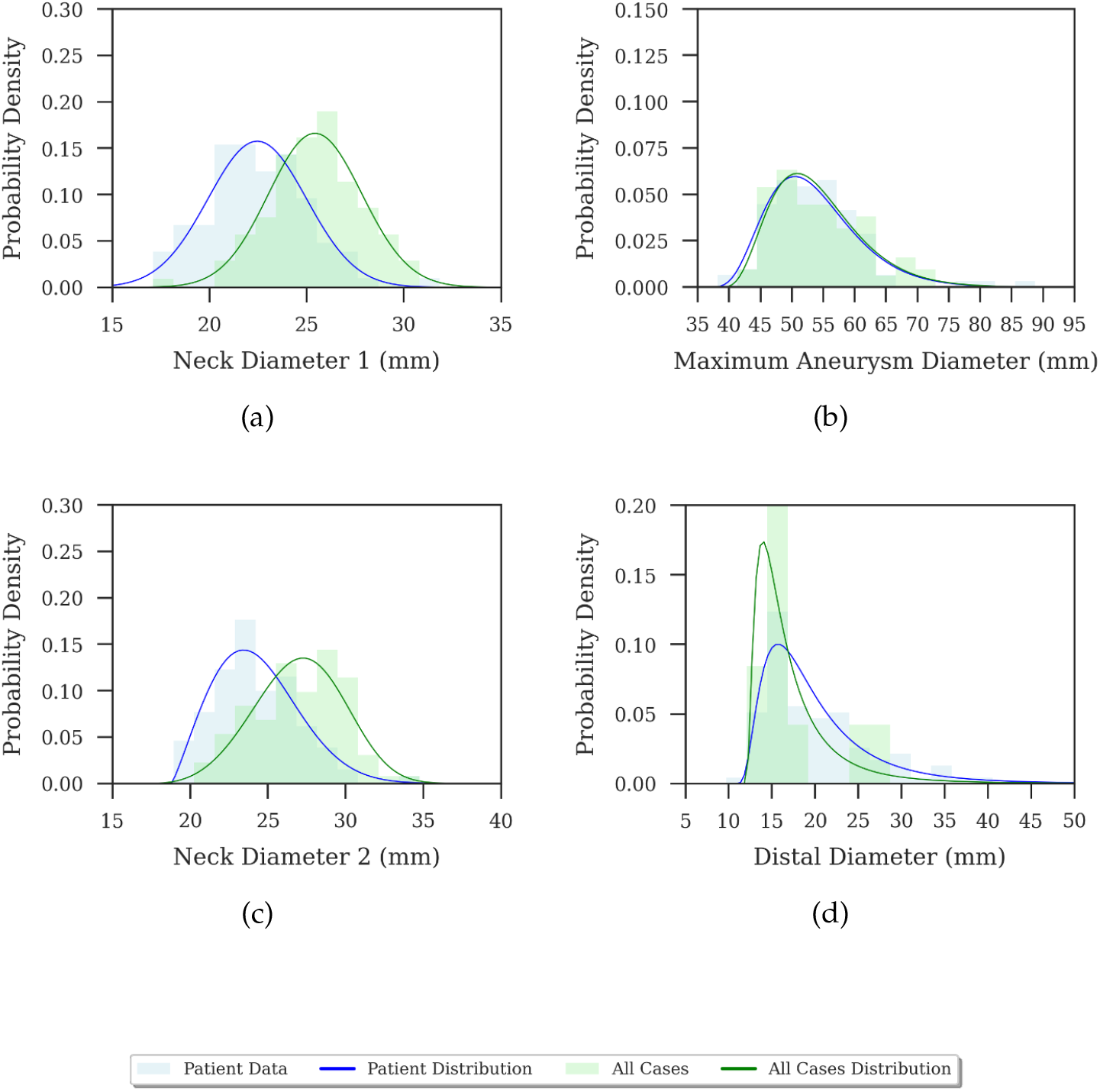
Comparison of patient data vs. generated statistical variants for the male 70-79 age group. (a) Neck 1 diameter comparison. (b) Maximum aneurysm diameter comparison. (c) Neck 2 diameter comparison. (d) Distal diameter comparison. Each subfigure shows the probability density function of patient data (blue) alongside the morphed statistical variants (green). The third row provides an enlarged legend for the plots.

**Figure 27.**
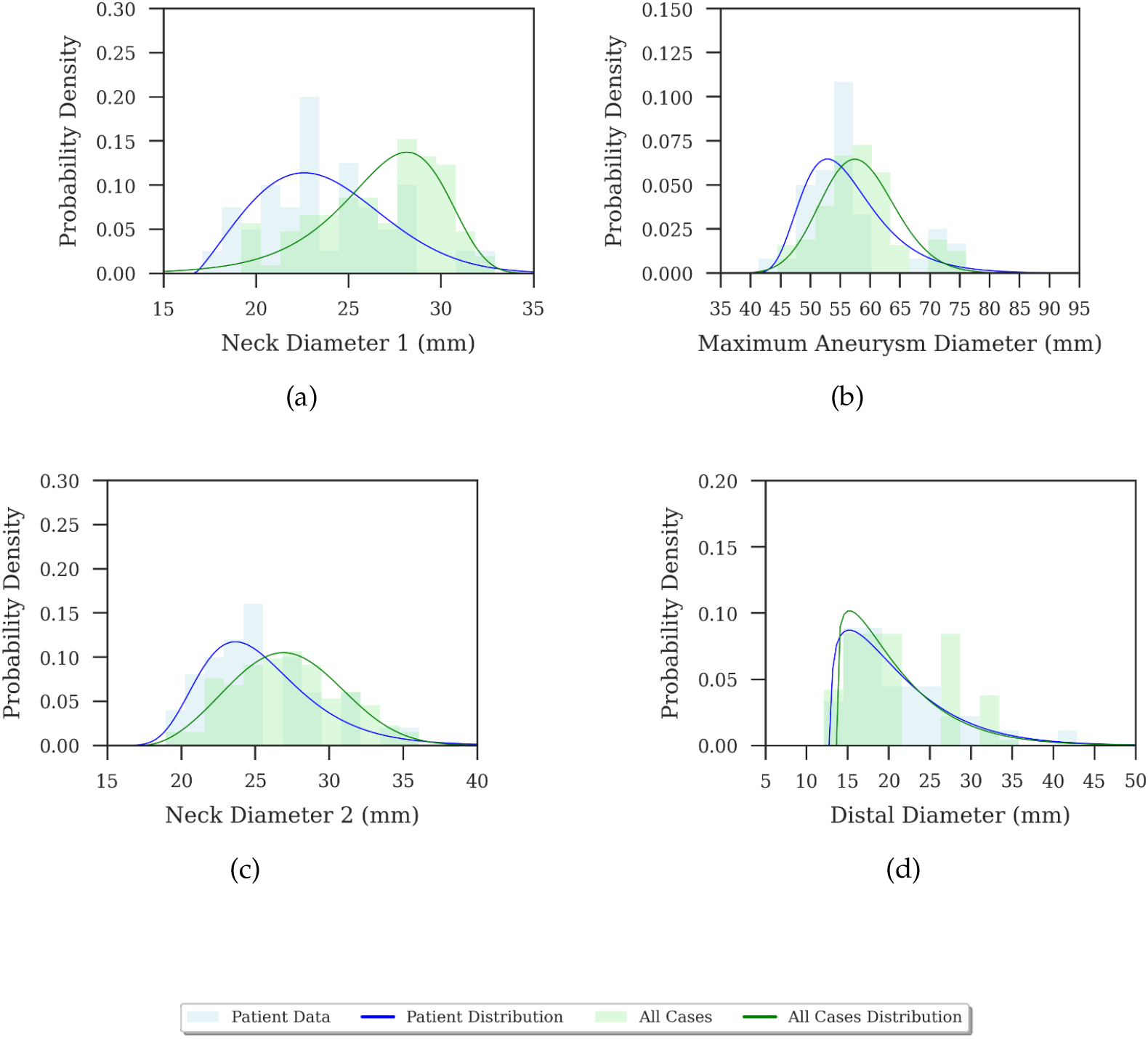
Comparison of patient data vs. generated statistical variants for the male 80+ age group. (a) Neck 1 diameter comparison. (b) Maximum aneurysm diameter comparison. (c) Neck 2 diameter comparison. (d) Distal diameter comparison. Each subfigure shows the probability density function of patient data (blue) alongside the morphed statistical variants (green). The third row provides an enlarged legend for the plots.

**Figure 28.**
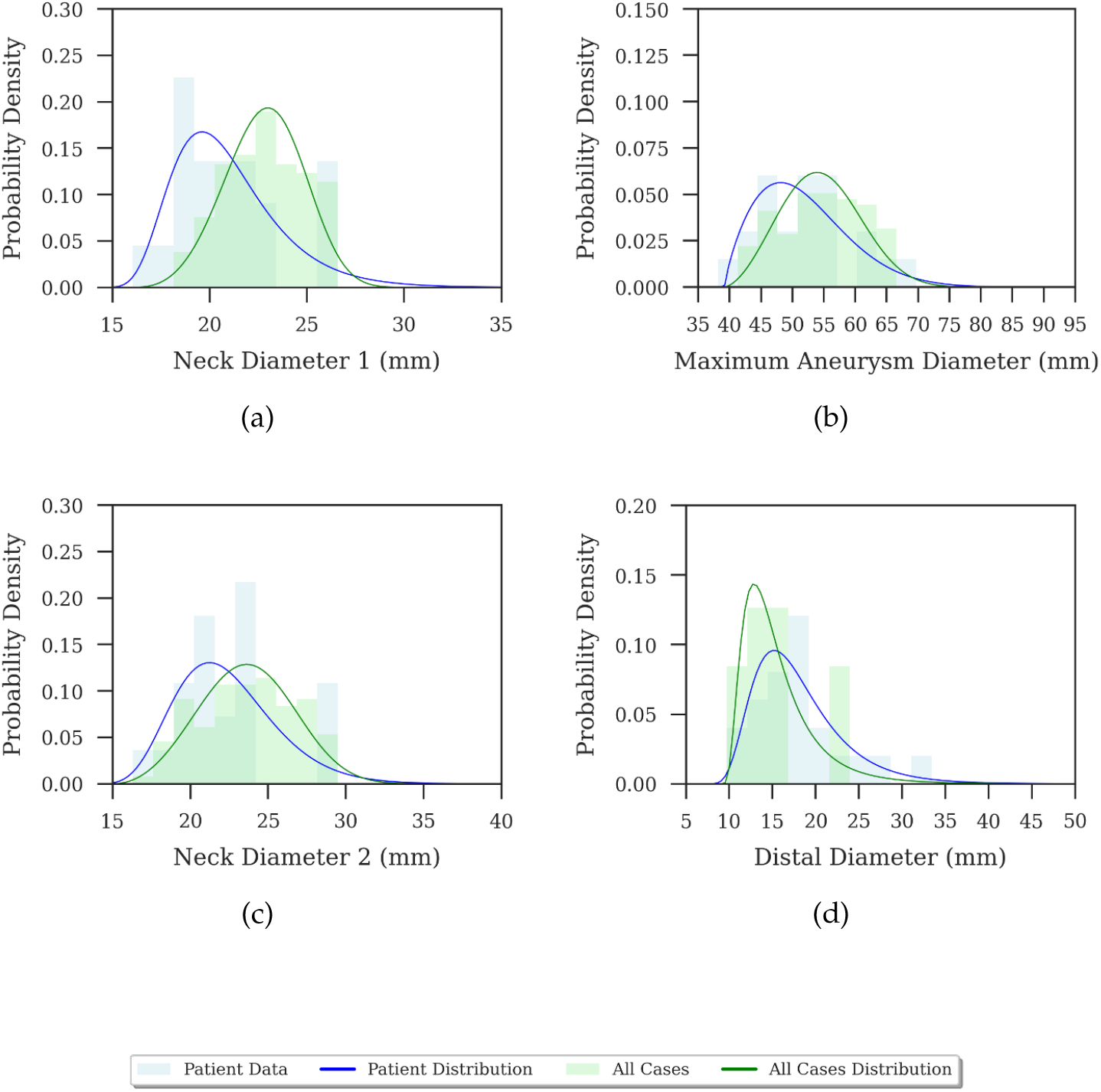
Comparison of patient data vs. generated statistical variants for the female 70-79 age group. (a) Neck 1 diameter comparison. (b) Maximum aneurysm diameter comparison. (c) Neck 2 diameter comparison. (d) Distal diameter comparison. Each sub-figure shows the probability density function of patient data (blue) alongside the morphed statistical variants (green). The third row provides an enlarged legend for the plots.

### J(b) Outlier Analysis

Seven patients (IDs: 40, 42, 71, 72, 78, 109, 163) has been considered as outliers for this study.

### J(c) Universal Interior Set Analysis

**Figure 29.**
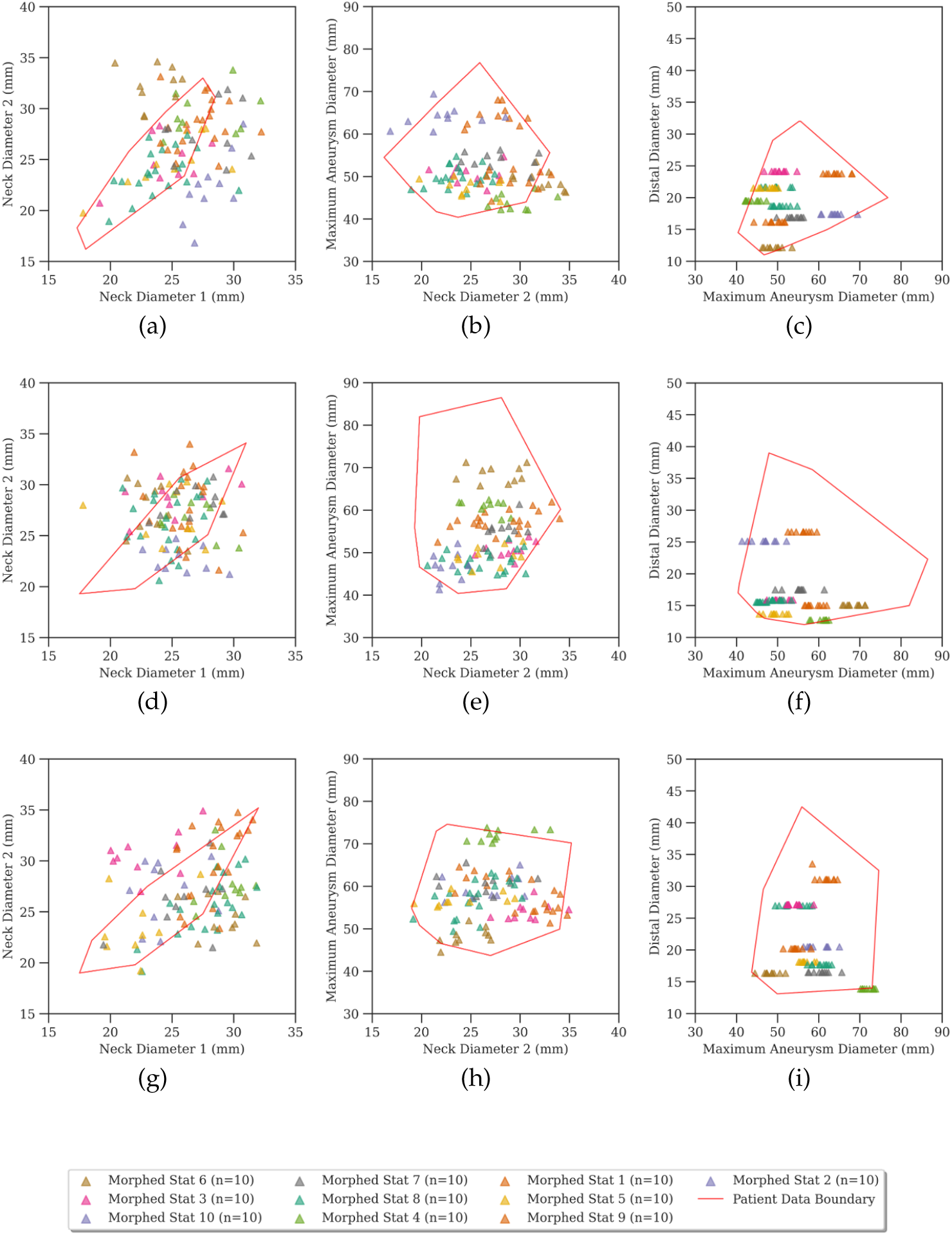
Parameter space comparison between patient data and morphed geometries for male patients aged 60-69, 70-79, and 80+. The convex hull of patient data (red) is overlaid with morphed geometry points. Subplots in each row represent relationships between neck diameter points, neck-to-aneurysm transitions, and aneurysm-to-distal transitions. The legend in the last row remains constant for all age groups.

**Figure 30.**
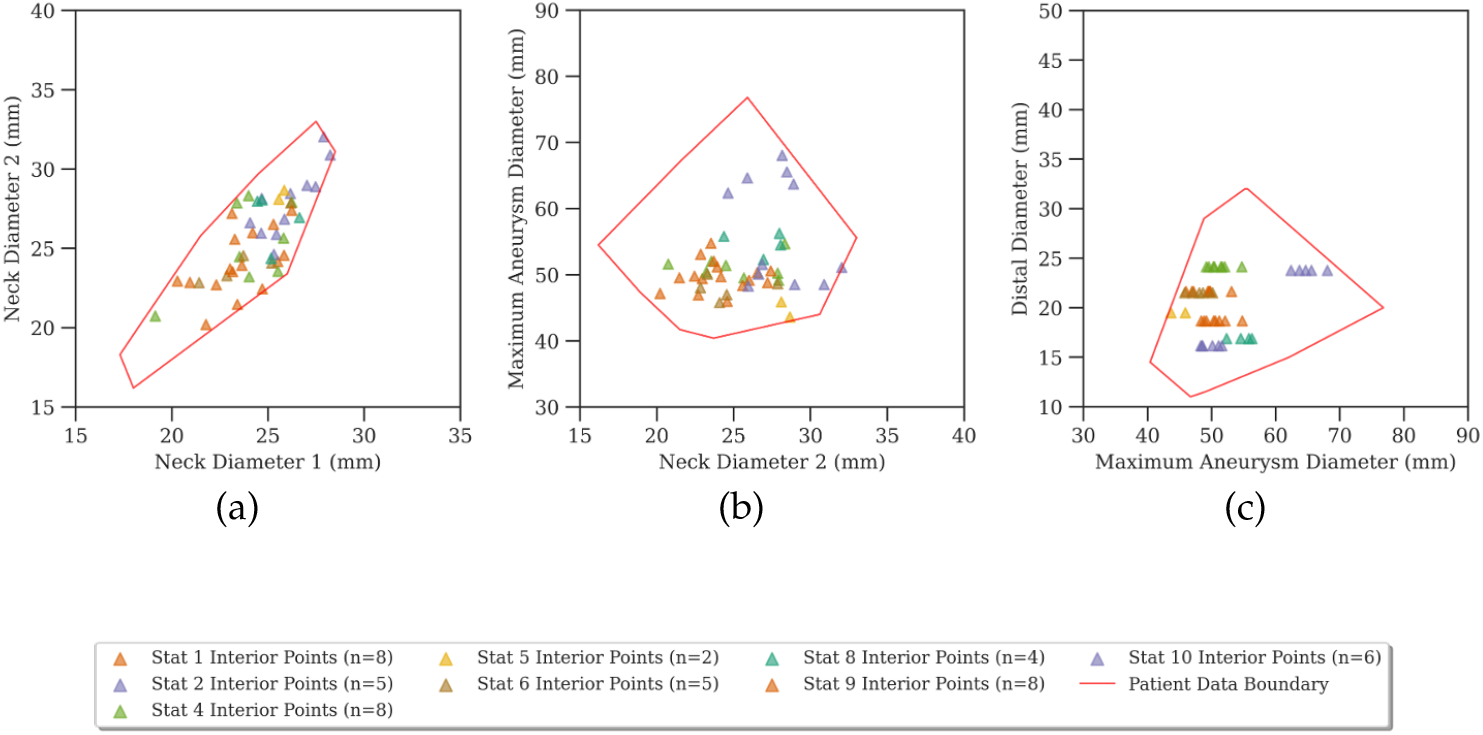
Universal interior set analysis for male patients aged 60-69. Valid morphed geometries are shown within the convex hull of patient data (a) Neck Diameter 1 vs. Neck Diameter 2, (b) Neck Diameter 2 vs. Maximum Aneurysm Diameter, (c) Maximum Aneurysm Diameter vs. Distal Diameter.

**Figure 31.**
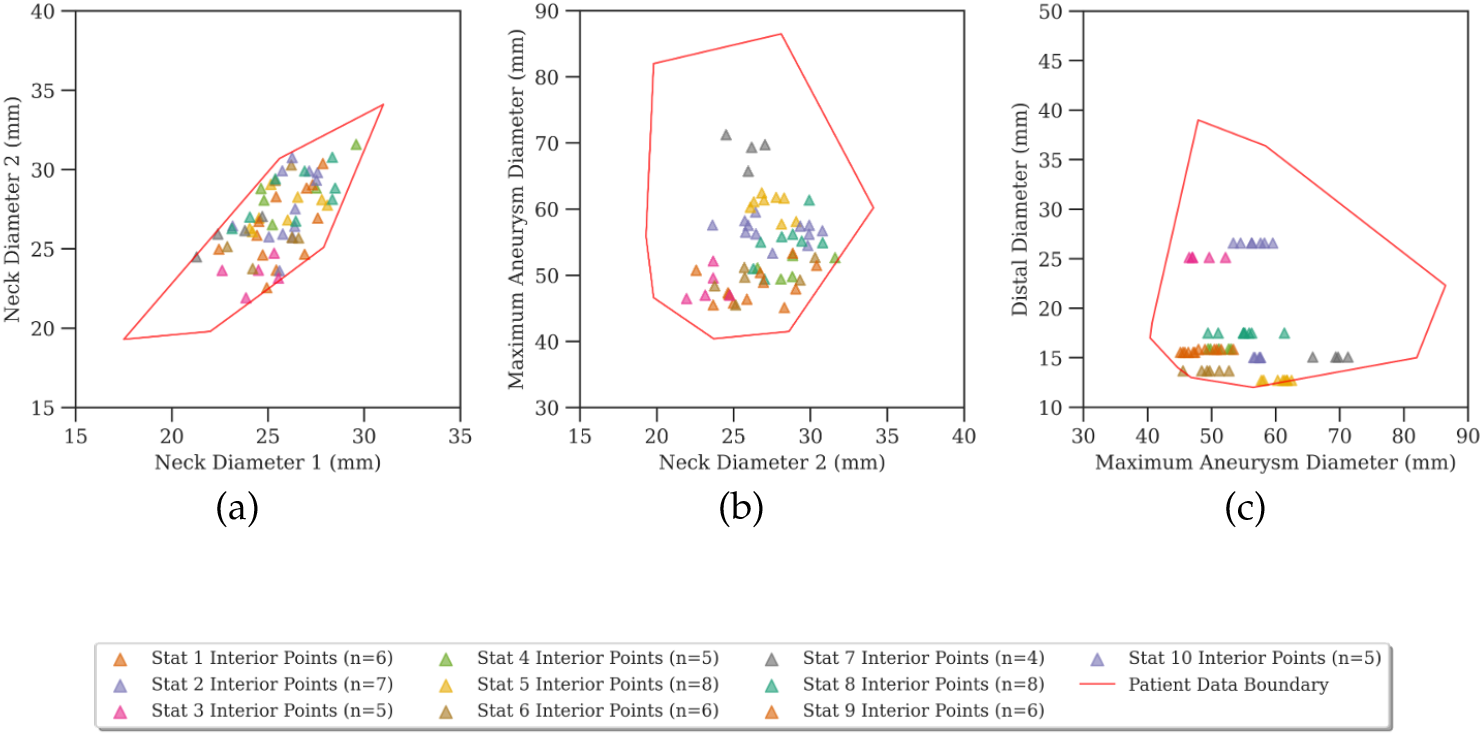
Universal interior set analysis for male patients aged 70-79. Valid morphed geometries are shown within the convex hull of patient data (a) Neck Diameter 1 vs. Neck Diameter 2, (b) Neck Diameter 2 vs. Maximum Aneurysm Diameter, (c) Maximum Aneurysm Diameter vs. Distal Diameter.

**Figure 32.**
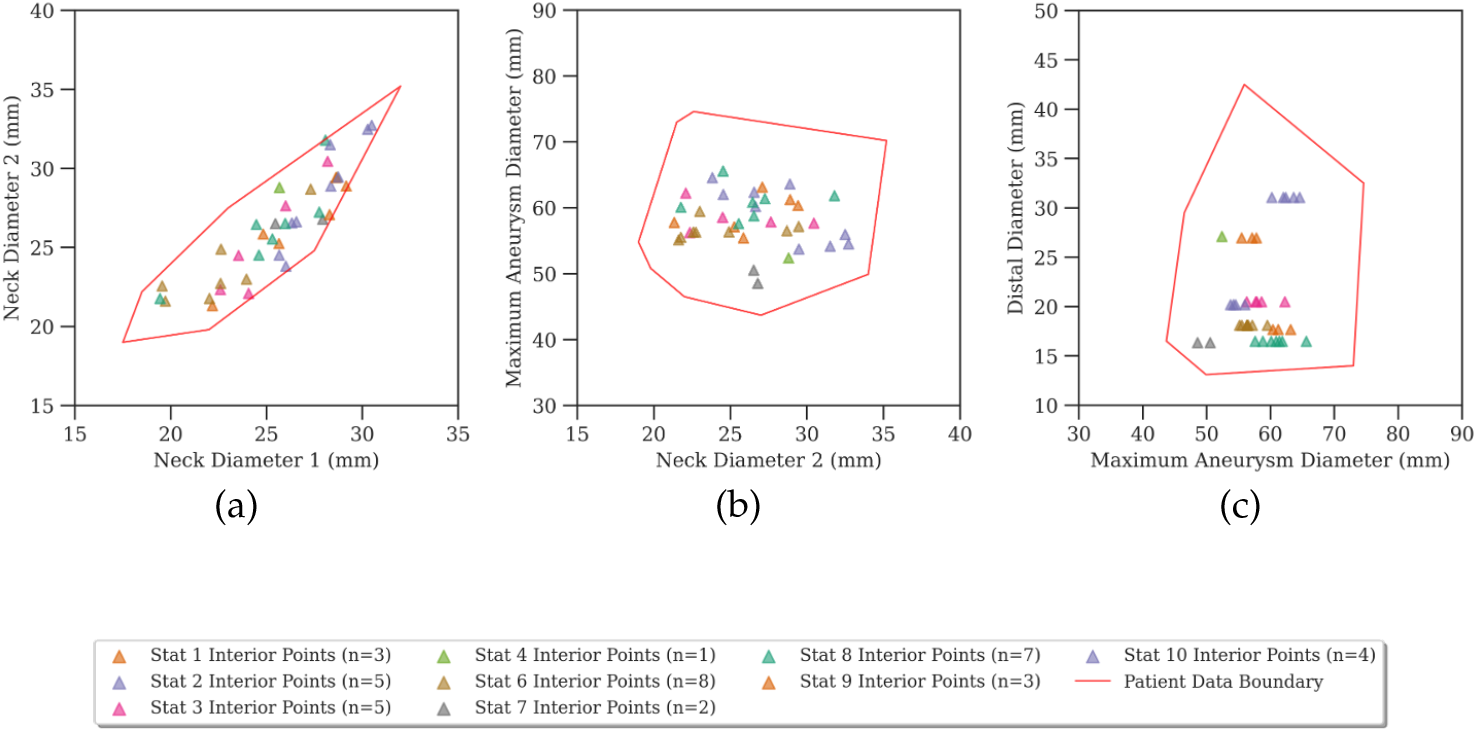
Universal interior set analysis for male patients aged 80+. Valid morphed geometries are shown within the convex hull of patient data (a) Neck Diameter 1 vs. Neck Diameter 2, (b) Neck Diameter 2 vs. Maximum Aneurysm Diameter, (c) Maximum Aneurysm Diameter vs. Distal Diameter

## Major abbreviations

AAA: Abdominal Aortic Aneurysm
CTA: Computed Tomography (CT) Angiography
EVAR: Endovascular Aneurysm Repair
VQI: Vascular Quality Initiative
CFD: Computational Fluid Dynamics
WSS: Wall Shear Stress
TAWSS: Time-Averaged Wall Shear Stress
OSI: Oscillatory Shear Index

