## Supplementary material for "Neck Geometry and Shape Compactness as Additional Haemodynamic Modulators in Abdominal Aortic Aneurysms": Electronic supplementary materials

### 1. GOVERNING EQUATIONS AND QUANTITIES RELATED TO CFD

This section provides the mathematical definitions of the governing equations and haemodynamic biomarkers used throughout the main manuscript. All biomarkers are computed from CFD simulations using OpenFOAM v9® with the `pisoFoam` transient solver.

#### A. Governing equations

Blood flow in the abdominal aorta is modelled as an incompressible Newtonian fluid. The governing equations are the continuity (mass conservation) and Navier-Stokes (momentum conservation) equations in their incompressible form

$$\nabla \cdot \mathbf{u} = 0, \tag{S1}$$

$$\frac{\partial \mathbf{u}}{\partial t} + (\mathbf{u} \cdot \nabla)\mathbf{u} = -\frac{1}{\rho}\nabla p + \nu \nabla^2 \mathbf{u} + \mathbf{g}, \tag{S2}$$

where $\mathbf{u}$ is the velocity vector $(\mathrm{m\,s^{-1}})$, $p$ is the kinematic pressure Pa, $\rho = 1060\ \mathrm{kg\,m^{-3}}$ is density of blood, $\nu = \mu/\rho$ is the kinematic viscosity with dynamic viscosity $\mu = 3.5\ \mathrm{mPa\,s}$, and $\mathbf{g}$ is gravitational acceleration. The full derivation and physical interpretation can be found in standard references such as, for example by Batchelor [1].

#### B. Wall shear stress quantities

Let $\vec{\tau}_w(x,t)$ denote the instantaneous wall shear stress vector at surface location $x$ and time $t$, $A$ the total lumen wall surface area, and $T = 1.0$ s the cardiac cycle period.

**Mean cycle WSS ($\mathrm{WSS_{mean}}$)** The mean cycle WSS ($\mathrm{WSS_{mean}}$) is the spatially and temporally averaged shear stress magnitude over a full cardiac cycle and given as

$$\mathrm{WSS_{mean}} = \frac{1}{T}\int_0^T \left( \frac{1}{A}\int_A |\vec{\tau}_w(x,t)|\,\mathrm{d}A \right) \mathrm{d}t. \tag{S3}$$

**Peak-systolic 95th-percentile WSS ($\mathrm{WSS}_{95\%}$)** The 95th percentile of the instantaneous wall shear stress magnitude ($\mathrm{WSS}_{95\%}$) evaluated at peak systole and used to capture peak shear conditions. This value highlights areas of the aneurysm wall exposed to high shear forces during maximum flow. This statistic is further preferred over the absolute maximum because isolated numerical spikes arising from local mesh irregularities can inflate the maximum WSS values without reflecting genuine flow physics. Thus, reporting the 95th percentile instead of the maximum helps avoid these outliers and provides a more stable and meaningful estimate of peak shear stress.

**Time-averaged WSS (TAWSS)** Unlike $\mathrm{WSS}_{\mathrm{mean}}$, which aggregates a single value over the entire surface, time-averaged wall shear stress (TAWSS) retains spatial resolution and indicates the sustained shear environment at each location on the vessel wall. The TAWSS is computed pointwise across the surface as

$$\mathrm{TAWSS}(x) = \frac{1}{T}\int_0^T |\vec{\tau}_w(x,t)|\,\mathrm{d}t. \tag{S4}$$

Regions where $\mathrm{TAWSS} < 0.4\,\mathrm{Pa}$ are identified as prone to thrombus formation and low-shear associated endothelial dysfunction [2]. Further, the TAWSS is obtained by integrating over the full cardiac cycle and hence the transient WSS spikes (e.g. from mesh artefacts) contribute negligibly to the spatial patterns of sustained shear.

### C. Oscillatory Shear Index (OSI)

OSI quantifies the directional variability of wall shear stress over the cardiac cycle at each surface point:

$$\mathrm{OSI} = \frac{1}{2}\left(1 - \frac{\left|\int_0^T \vec{\tau}_w(t)\,\mathrm{d}t\right|}{\int_0^T |\vec{\tau}_w(t)|\,\mathrm{d}t}\right), \tag{S5}$$

where OSI is zero for purely unidirectional flow and 0.5 for completely oscillatory (zero net) flow. The fraction of the wall area where $\mathrm{OSI} > 0.3$ is used to quantify the spatial extent of disturbed, multidirectional flow associated with adverse vascular remodelling [3]. Since OSI is a ratio of time-integrated quantities, it is comparatively insensitive to brief, localised numerical excursions in $\vec{\tau}_w$, provided these spikes are not persistent over a substantial fraction of the cycle.

Together, these metrics complement traditional magnitude-based shear stress measures by highlighting regions of complex, multidirectional flow that may contribute to aneurysm progression [3].

### 2. SOFTWARE VERIFICATION REPORT

An audit of the entire framework was considered, in two steps, to enhance the trustworthiness of the developed framework and software. First, it was decomposed into its constituent computations and each computation was identified as using either a standard library or custom code. The correctness check for each computation is documented in Figure S1. Second, the custom shape measurement modules, where other potential bugs could be hidden, was re-verified against analytical reference geometries.

Figure S1 maps every computational step of the framework onto the four stages described in the manuscript (dataset description, geometric modeling, CFD framework, correlation analysis). For each step, the figure names the utility or library used and its verification basis. The majority of steps rely on standard, widely adopted scientific libraries such as `SciPy`, `NumPy` and `OpenFOAM`, whose correctness is established through extensive community testing and upstream validation. For these, our verification concerns correct usage rather than implementation and is documented per block (e.g. `checkMesh` and mesh convergence for meshing, solver logs and Courant diagnostics for the transient solver, KL-divergence checks for morphing).

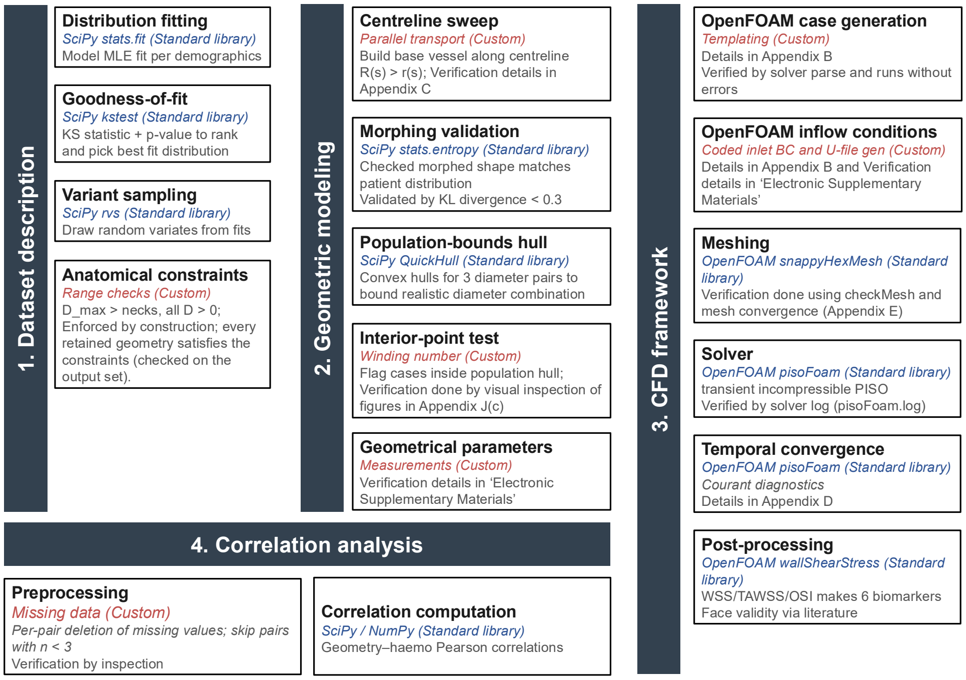


**Fig. S1.** Verification map of the framework. Each step lists the library or code used. The standard libraries in blue, custom code in red with its verification basis.

Custom code is highlighted in red. It is confined to a small number of components including anatomical range checks, the parallel transport centreline sweep, the winding number interior point test, geometrical parameter measurement, OpenFOAM case templating and inlet boundary condition generation, and missing data handling in preprocessing. Each red block states the verification or validation performed for that component, with pointers to the relevant appendix or to this report. This decomposition ensures that every custom computation in the framework is covered by a verification activity.

### A. Shape descriptors mapped to reference geometries

The shape descriptors used in this work was extracted and compared for standard geomertric shapes, where these metrics are well known. The Figure S2 shows each descriptor evaluated on reference geometries with known or analytically derivable values.

**Morphed diameter (a-c)** Geometries were generated with prescribed morphing amplitudes (22, 50, and 70 mm). The framework's measured maximum diameter recovered the prescribed value to machine precision with errors $\leq 1.4 \times 10^{-14}$.

**Surface area and volume (d-f)** The surface area was evaluated on a pyramid, cube, and torus for which closed-form values are known. The agreement is exact to mesh resolution (e.g. cube is 9600 $mm^2$, 64000$mm^3$).

**Tortuosity (g-i)** A straight cylinder, quarter-circle arc, and $S$-shaped vessel with analytical tortuosities of 1, 1.110721, and 1.666667 were recovered with relative errors below $3 \times 10^{-6}$%.

**Sphericity (j-l)** Evaluated on ellipsoids of decreasing aspect ratio. The computed value increases monotonically toward the theoretical maximum of 1, reaching 0.9999 for the sphere.

**Convexity (m-o)** Evaluated on shapes of increasing concavity, from a convex ellipsoid (1.000) through mild (0.9000) to pronounced (0.7634) waisting.

**Average radius (p-r)** Compared against analytical values for a sphere, cube, and cylinder, with relative errors below $10^{-6}$%.

Together with the framework decomposition, these results demonstrate that the shape-measurement modules agrees well with analytical references across all descriptors.

### B. Convexity and sphericity

Particular attention is required for sphericity and convexity as they are both derived from mesh volume and used in correlation metrics. The pipeline computes that volume by the discrete divergence theorem in

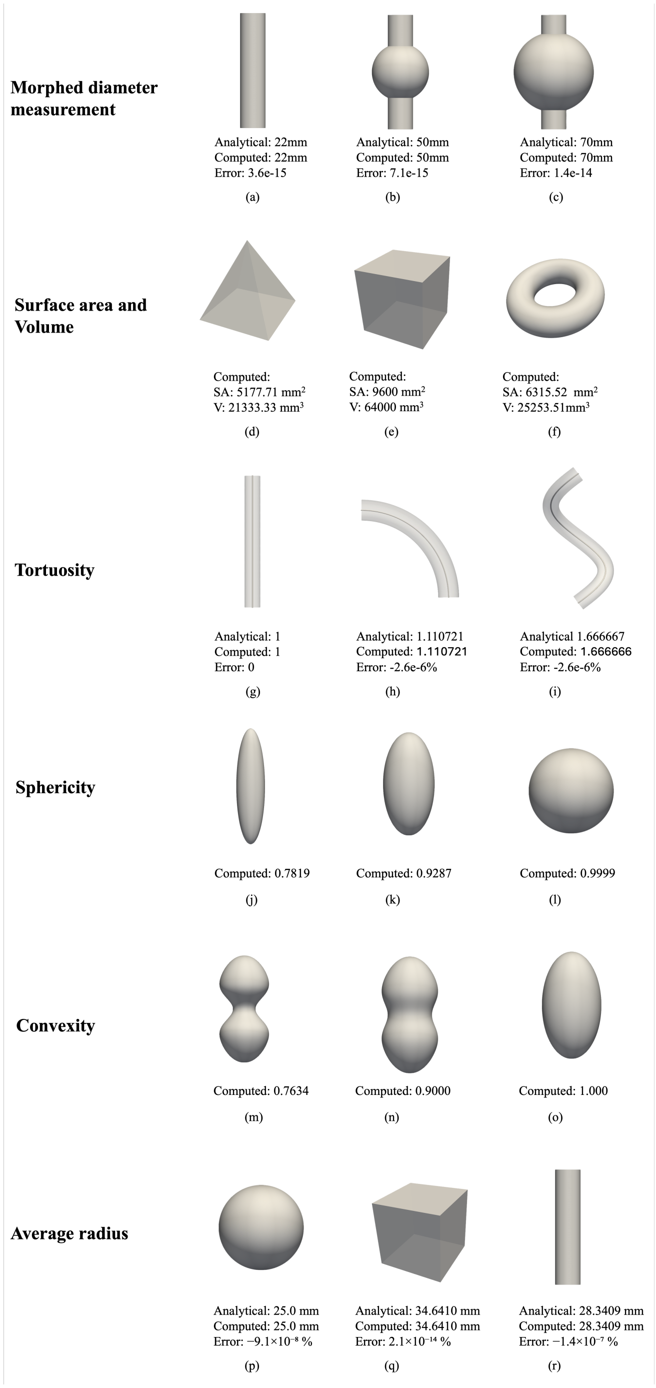


**Fig. S2.** Shape descriptors evaluated on reference geometries. Computed values are compared against analytical solutions where available which each triangular face of the surface mesh forms a tetrahedron with the coordinate origin and the signed volumes of all such tetrahedra are summed,

$$V = \frac{1}{3} \sum_{i=\text{faces}} \left( \mathbf{c}_i \cdot \hat{\mathbf{n}}_i \right) A_i, \tag{S6}$$

where $\mathbf{c}_i$ is the position vector of face $i$'s centroid, $\hat{\mathbf{n}}_i$ its outward unit normal and $A_i$ is its area. Thus, for faces whose outward normal points away from the origin this term is positive and for faces whose normal points toward the origin (the side of the surface nearer the origin or any concave region facing it) the term is negative. The negative contributions cancel the volume of the region between the origin and the near side of the surface, leaving the required enclosed volume once all terms are summed.

The volume routine is written to accumulate the signed per-face contributions first and then apply the absolute value once to the final total

```
# signed, no per-face abs()
vol += np.dot(centroid, n_hat) * area / 3

# abs() taken once on the total
mesh_volume = abs(vol)
```

This is equivalent to the signed-volume routine used internally by `trimesh` and the pipeline calls directly as well. Automatic range checks are also added on sphericity and convexity to ensure that the values outside the physically valid range $(0, 1]$ is flagged immediately rather than passing through unnoticed.

The verification considered four primitive shapes since their analytical solutions are well known. The STL files were generated and processed to extract the metrics for comparison.

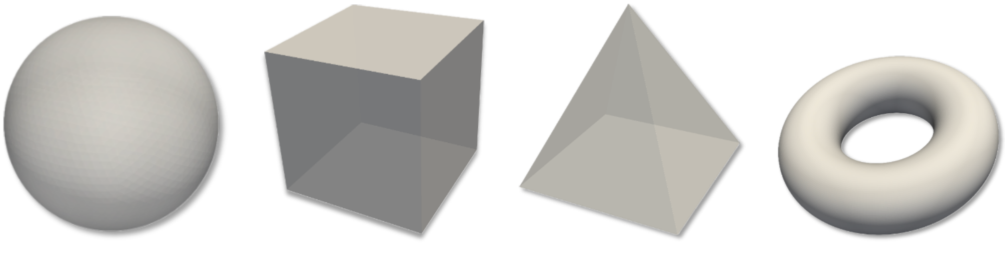

**(a)** **(b)** **(c)** **(d)**

**Fig. S3.** Shapes for ground truth verification: (a) Sphere ($r = 25\,\text{mm}$), (b) Cube ($a = 40\,\text{mm}$), (c) Pyramid ($a = h = 40\,\text{mm}$), (d) Torus ($R = 20$, $r = 8\,\text{mm}$). The torus is the only non-convex shape (Table S1).

The common bug is for the non-convex shapes (like torus) where there is a flip in the normal direction. This is also commonly the case of AAA geometries. This can inflate the volume since the negative contributions can be added rather than subtracted. Table S1 compares analytical and computed values for each shape and metric.

**Table S1.** Shape metrics on analytical primitives

| Shape | Metric | Analytical | Computed | %Error |
|---|---|---|---|---|
| **Sphere** | Volume | 65449.8 | 65308.4 | −0.22 |
| | Surface | 7854.0 | 7844.6 | −0.12 |
| | Sphericity | 1.0000 | 0.9998 | −0.02 |
| | Convexity | 1.0000 | 1.0000 | 0.00 |
| **Cube** | Volume | 64000.0 | 64000.0 | 0.00 |
| | Surface | 9600.0 | 9600.0 | 0.00 |
| | Sphericity | 0.8060 | 0.8060 | 0.00 |
| | Convexity | 1.0000 | 1.0000 | 0.00 |
| **Pyramid** | Volume | 21333.3 | 21333.3 | 0.00 |
| | Surface | 5177.7 | 5177.7 | 0.00 |
| | Sphericity | 0.7184 | 0.7184 | 0.00 |
| | Convexity | 1.0000 | 1.0000 | 0.00 |
| **Torus** | Volume | 25266.2 | 25253.5 | −0.05 |
| | Surface | 6316.5 | 6315.5 | −0.02 |
| | Sphericity | 0.6592 | 0.6591 | −0.02 |
| | Convexity | 0.7243 | 0.7241 | −0.02 |

The framework is applied to real AAA geometries, to confirm that this applies through for all cases. Figure S4 illustrates a representative AAA surface. In order to verify its applicability to AAA geometries, four AAA cases were selected from the virtual population to represent the extreme values of sphericity and convexity in the dataset. Together, the representative samples span the full geometric range of the cohort in these two shape descriptors.

### C. Inlet waveform boundary condition

The inlet boundary condition imposes a parabolic, time-varying velocity profile at the model's inlet patch. The waveform itself is supplied to

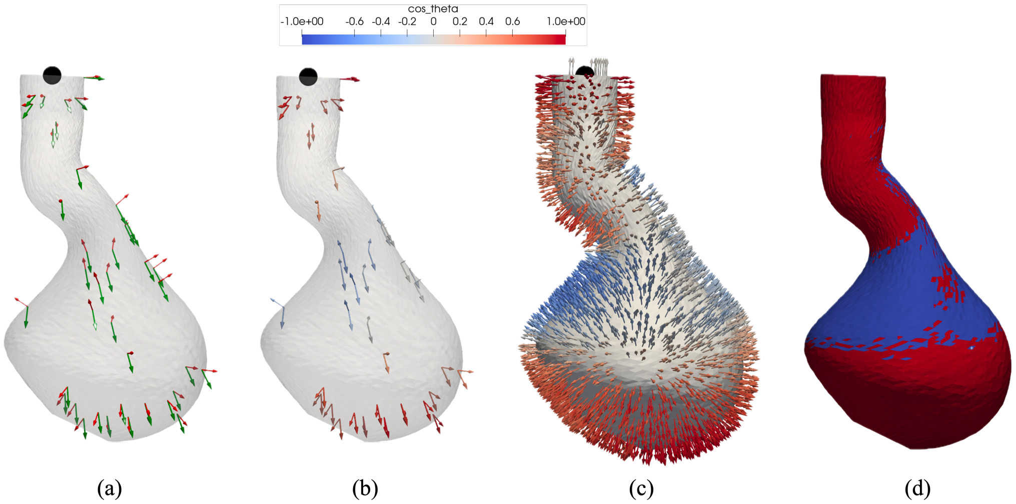


**Fig. S4.** Face-normal visualisation on AAA case: (a) Position vector **c** (green) and outward normal $\hat{\mathbf{n}}$ (red) as arrow pairs; (b) the same pairs coloured by $\cos\theta$; (c) the normal field over many more faces; (d) surface coloured by the sign of the volume contribution. Blue faces represent ($\cos\theta < 0$) non-convex faces.

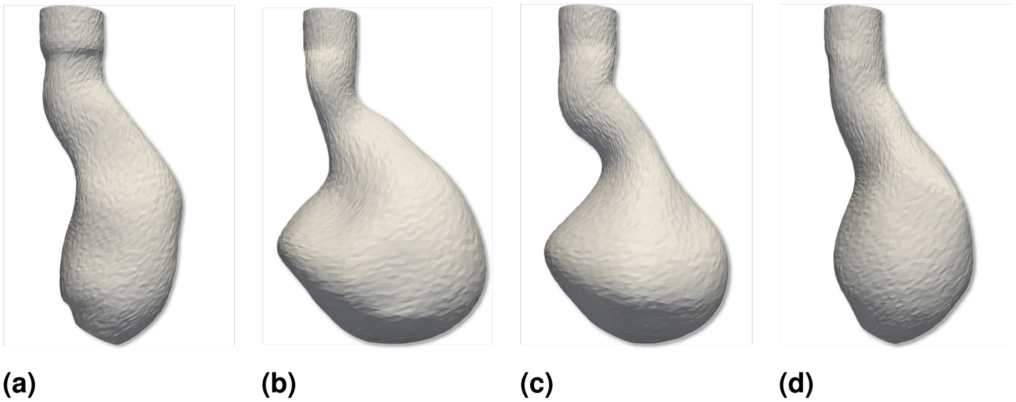


**Fig. S5.** Four AAA geometries used for end to end verification. (a) Min. sphericity (M, 60-69, stat-5, morph 8); (b) Max. sphericity (M, 70–79, stat-7, morph 2); (c) Min. convexity (F, 70–79, stat-3. morph 3); (d) Max. convexity (M, 60–69, stat-10, morph 3

**Table S2.** Shape metrics and geometry for the four extreme AAA cases

| Case | Selection | Sphericity | Convexity | Volume ($mm^3$) | Surface area ($mm^2$) |
|---|---|---|---|---|---|
| **M 60-69, morph∼8** | Min sphericity | 0.7665 | 0.7697 | 115693 | 14980 |
| **M 70-79, morph∼2** | Max sphericity | 0.8518 | 0.7468 | 252844 | 22702 |
| **F 70-79, morph∼3** | Min convexity | 0.8069 | 0.6854 | 187820 | 19656 |
| **M 60-69, morph∼3** | Max convexity | 0.8000 | 0.8143 | 135928 | 15982 |

OpenFOAM as a list of (time, velocity) pairs in the `U` file, preceded by a declared count, `numPoints`, that tells the solver how many time-velocity pairs follow. The cycle period is computed internally from the first and last declared time points (`timePoints.last() - timePoints.first()`).

The waveform used to simulating the blood flow for the AAA case (M, 60-69, stat-10, morph-4). The inlet centreline velocity is extracted from the solver written field files at every saved timestep ($t = 0$ to 3.8,s, $\Delta t = 0.05$,s). The results shown in Figure S6 shows that the simulation tracks the input exactly across all four cardiac cycles.

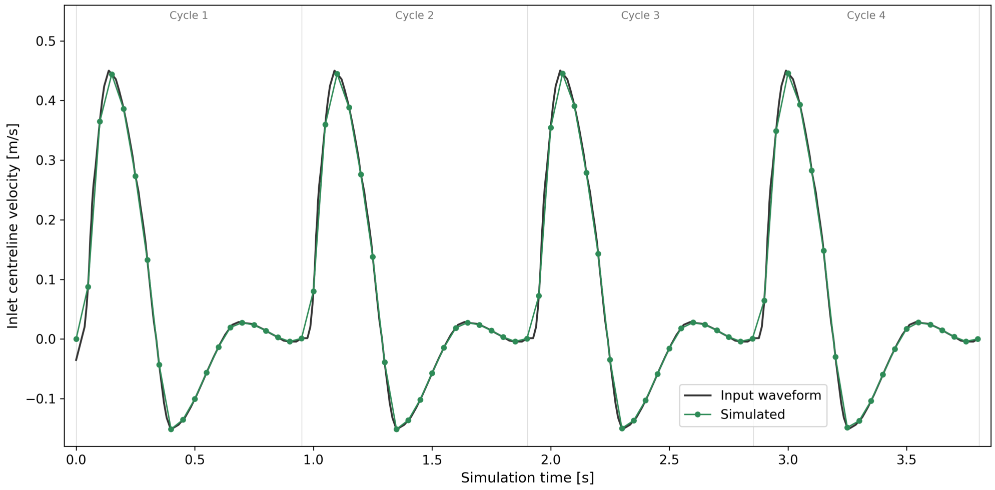


**Fig. S6.** Inlet velocity waveform (4 cardiac cycles)

The waveform is further confirmed by examining inlet velocity vectors at four characteristic instants of the cardiac cycle using ParaView's Glyph filter (Figure S7). The result represents the expected physiological pattern with maximum positive velocity at peak systole, flow reversal at incisura, maximum negative velocity at peak diastole and near-zero velocity at end diastole.

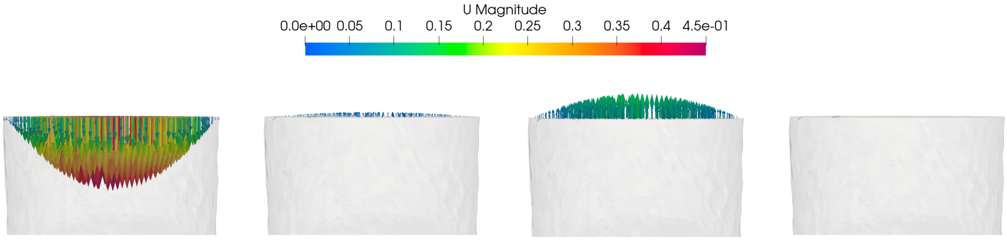


(a) (b) (c) (d)

**Fig. S7.** Inlet velocity vectors at four key cardiac phases: (a) peak systole (b) incisura (c) peak diastole (d) end diastole

### D. Morphed centerline

The morphed centerline is, by definition, the locus of cross-sectional centroids along the morphed geometry. Each centroid can be estimated by plane-slab slicing. At every point of the centerline, all mesh vertices lying within a thin slab perpendicular to the local tangent can be collected and their mean taken as the centroid. While workable, this approach is sensitive to how the slab thickness is chosen, because morphing displaces vertices axially as well as radially. A slab thin enough to isolate a single cross-section can omit vertices that have moved out of it, whereas a thicker slab can include vertices from neighbouring cross-sections and bias the centroid laterally. Both effects introduce small errors into the centroid, and hence into the centerline length and tortuosity derived from it.

Alternative, this work groups the mesh vertices into cross-sectional rings using the known structure of the wall mesh, rather than by geometric slicing. It uses the already present mesh generation routine `Cylinder_Triangulation_Continuous_N`, which produces a fixed number of vertices per ring in a fixed order. Morphing only moves these vertices but never adds, removes or reorders them. Thus, each ring therefore always consists of the same set of vertices, regardless of how the geometry is deformed. The centroid of each ring is then computed directly as the mean of its vertices, with no slicing, no nearest-neighbour search, and no dependence on a slab-thickness parameter.

The method is quantified through the Pearson correlation between tortuosity and centerline length across the full cohort. By construction,

$$\text{tortuosity} = \frac{\text{centerline length}}{\text{straight length}} \tag{S7}$$

where straight length is the inlet to outlet distance. The inlet and outlet rings are held fixed by the morphing boundary conditions, the straight length is constant across all cases. Thus, the tortuosity and centerline length are exact linear rescalings of one another and their correlation is expected to be 1.

The simplistic plane-slab method yielded $r = 0.954$, the small deflection reflecting its sensitivity to slab thickness. The topology-based method recovers the expected $r = 1.000$, confirming that the morphed centerline is computed consistently and exactly.
